# Renal Coenzyme A (CoA) Production from VB5 Fuels Stem Cell Proliferation and Tumor Growth

**DOI:** 10.1101/2025.08.08.669325

**Authors:** Ting Miao, Ying Liu, Mujeeb Qadiri, Amaury Dasseux, John M Asara, Yanhui Hu, Xiaomei Sun, Luz del Carmen Pliego-Alcántara, Christian C. Dibble, Norbert Perrimon

## Abstract

Coenzyme A (CoA), derived from Vitamin B5 (VB5; also called pantothenate), is essential for lipid metabolism, energy production, and cell proliferation. While the intracellular functions of CoA are well-characterized, much less is known about its tissue-specific regulation and systemic physiological roles. Here, using *Drosophila melanogaster*, we uncover a gut-renal circuit in which dietary VB5 fuels CoA biosynthesis specifically in the Malpighian tubules (MTs, the fly kidney), non-autonomously impacting gut homeostasis. We show that, in the MTs, Myc boosts renal CoA production by directly upregulating the pantothenate kinase *Fbl* (human *PANK1-3* ortholog) and downregulating *CG5828*, which we characterize as the functional ortholog of the metabolite phosphatase and CoA synthesis suppressor PANK4 (*dPANK4*). Elevated CoA biosynthesis enhances the mevalonate-isoprenoid pathway activity in the gut, promoting intestinal stem cell proliferation. We further demonstrate that renal CoA production is required for gut tumor growth in a fly model. Consistently, *MYC* and genes within the CoA-isoprenoid axis display strong association with clinical outcomes in human cancers. Together, our findings establish that Myc-driven CoA metabolism generates an inter-organ signal that couples VB5 availability to stem cell control and tumor growth, and identify the CoA-isoprenoid axis as a targetable metabolic vulnerability in cancer.

## Introduction

Dietary supplementation with B vitamins is a common practice worldwide [1], valued for their beneficial roles in fundamental biological processes such as energy production, nervous system function, and immune responses [2, 3]. While all B vitamins are essential nutrients for metazoans and function as critical cofactors or cofactor precursors in central metabolism, their dietary effects are not uniformly advantageous. Under certain conditions, they can produce unexpected and even adverse outcomes, for example, promoting leukemic cell proliferation [4]. These findings underscore that our understanding of B vitamin metabolism in physiological and pathological states remains incomplete.

A particularly important member of this family is Vitamin B5 (VB5), or pantothenic acid/ pantothenate, which exclusively serves as a precursor of Coenzyme A (CoA) in all organisms [5]. CoA is a central metabolic cofactor required for over 100 enzymatic reactions, including the tricarboxylic acid (TCA) cycle, fatty acid β-oxidation, amino acid catabolism, and lipid and sterol synthesis [5–8]. In either mammals or insects, VB5 cannot be *de novo* synthesized and is largely obtained from the diet. It is then absorbed in the intestine and transported into cells via SLC5A6/SMVT, enabling CoA biosynthesis across diverse tissues [7, 8]. CoA is synthesized *de novo* from VB5, cysteine, and ATP via a conserved five-step enzymatic pathway, initiated by the phosphorylation of VB5 by pantothenate kinases (PANKs), of which there are three functional paralogs in vertebrates (PANK1-3) [7]. Notably, we recently found that PANK4, a pseudo-pantothenate kinase [9] with a DUF89-family metabolite phosphatase domain not found in PANK1-3 [10], is a rate-determining enzyme that negatively regulates CoA biosynthesis via its phosphatase activity in mammalian cells [11]. The pantothenate kinase activity of mammalian PANK1-3 (and of PANKs in other species including most bacteria) is modulated by direct feedback inhibition from CoA and acyl-CoAs [12, 13], and responds to nutrient signals such as insulin via post-translational modifications, including PI3K/Akt-mediated phosphorylation of PANK2 and PANK4 [11]. Transcriptional control is another important regulatory layer, but only a limited number of transcription factors have so far been uncovered that regulate genes involved in CoA transport or biosynthesis. For example, PANK1 is a direct transcriptional target of the tumor suppressor and transcription factor p53 [14, 15], while the oncogenic transcription factor MYC has been shown to regulate CoA biosynthesis by driving the expression of the cell surface VB5 transporter SLC5A6/SMVT [16, 17]. Whether MYC also directly regulates CoA biosynthetic enzymes to control CoA production remains unknown. Notably, total CoA levels differ by more than 10-fold across mammalian tissues [7], suggesting that CoA metabolism is highly tissue-specific and coordinated across tissues. However, the mechanisms underlying tissue-specific CoA metabolic regulation remains poorly defined, limiting our understanding of how CoA metabolism is tuned across organs in response to physiological and pathological cues. This is particularly important in high-metabolic-turnover organs such as the kidney, which exhibits high CoA levels [7, 8] but whose regulatory networks governing CoA metabolism remain poorly understudied.

CoA availability is essential for lipid metabolism, particularly fatty acid (FA) and mevalonate (MVA) biosynthesis [18, 19]. FA synthesis begins with acetyl-CoA and malonyl-CoA and yields diverse lipids for membranes and energy storage [18, 20]. The MVA pathway, which initiates with a multi-step conversion of acetyl-CoA to 3-hydroxy-3-methylglutaryl-CoA (HMG)-CoA, is essential in multiple cancer cell types [21–24]. Pharmacological inhibition of this pathway by targeting the rate-limiting enzyme HMG-CoA reductase (HMGCR) with statins reduces tumor growth and promotes apoptosis, underscoring the dependency of tumors on MVA pathway activity [21–24]. A major output of the MVA pathway is the synthesis of isoprenoid backbones, catalyzed by key enzymes such as farnesyl diphosphate synthase (FDPS) and geranylgeranyl diphosphate synthase 1 (GGPS1). These backbones serve as precursors for a range of downstream metabolic products, including ubiquinone (CoQ), dolichol, sterols, and prenylation substrates [19]. While much focus has been placed on cholesterol as the major tumor-promoting metabolite [25], emerging evidence suggests that non-sterol isoprenoids also influence cancer cell behavior. For example, the availability of geranylgeranyl pyrophosphate (GGPP), a key isoprenoid intermediate mediating protein prenylation, has been shown to promote cancer cell proliferation [26, 27]. Nevertheless, although other isoprenoid-derived metabolites are known to participate in key cellular processes such as cell cycle progression and mitochondrial function [19, 28], their specific contributions to tumor growth and survival remain largely undefined.

*Drosophila* is a powerful model organism for investigating tissue homeostasis, metabolism, and inter-organ communication [29, 30]. Importantly, flies share conserved VB5-CoA biosynthetic pathways with mammals and possess tissue analogs to major human organs, making them an ideal system for studying tissue-specific CoA metabolism. In this study, we investigated how VB5-CoA metabolism influences gut tissue homeostasis and tumor growth in flies. We show that dietary VB5 supplementation or renal-specific activation of CoA biosynthesis is sufficient to drive intestinal stem cell (ISC) proliferation via enhanced MVA-isoprenoid pathway in the gut. In addition, we identify Myc as a direct transcriptional regulator of this pathway. In a Yorkie/Yki-driven fly gut tumor model [31], tumor-derived PDGF/VEGF signaling upregulates *Myc* in renal cells, which in turn drives CoA biosynthesis to support tumor growth. Importantly, we demonstrate the clinical relevance of the MYC-driven CoA-isoprenoid axis in human cancer progression, and identify the CoA-isoprenoid pathway as a potential therapeutic target in kidney cancer. Together, our work establishes a functional inter-organ circuit of CoA metabolism, with broad implications for understanding systemic metabolic regulation and developing therapeutic interventions in cancer.

## Results

### VB5 supplementation stimulates intestinal stem cell ISC proliferation

While the B vitamins are essential for systemic health [2, 3], their specific roles in regulating tissue homeostasis remain poorly characterized. To investigate this, we employed the *Drosophila* midgut, a highly plastic organ that serves as an ideal model for dissecting the nutritional and metabolic control of tissue homeostasis [32–34]. We supplemented fly diets with major B vitamins individually, including thiamine (VB1), riboflavin (VB2), niacin (VB3), pantothenate (VB5), pyridoxine (VB6), and biotin (VB7) **(Fig. 1a)**. Notably, VB5 supplementation led to significant expansion of the midgut, reflected by increased width in the R4-R5 regions **(Fig. 1b-c)**, suggesting a role of VB5 in maintaining midgut homeostasis. Since *Drosophila* midgut homeostasis is maintained by proliferating intestinal stem cells (ISCs) [32, 33], we assessed the ISC proliferation under these conditions through staining for phospho-Histone 3 (pH3), a marker of mitotic activity [32, 33]. Consistent with the observed midgut expansion, VB5-fed flies exhibited a significant increase in pH3-positive (pH3^+^) ISCs compared to controls **(Fig. 1b, d)**, suggesting enhanced ISC proliferation. We further validated this phenotype by supplementing fly diets with a range of VB5 concentrations. Interestingly, we observed a modest increase in pH3^+^ signal with 1 mM VB5 supplementation and a more pronounced effect at 2.5 mM, whereas no induction was observed at the higher concentration at 5 mM **(Fig. 1e)**. Because the cornmeal-based fly diet naturally contains VB5, we next examined the dose-dependence using a *Drosophila* holidic medium [35–37] depleted of VB5 and supplemented with 0, 1, 2.5, 5, or 10 mM VB5 **(Fig. S1a)**. Under these conditions, the strongest induction of pH3⁺ signal was observed at 2.5 mM VB5 supplementation, with a statistically significant but smaller increase at 5 mM VB5 supplementation. Notably, 1 mM VB5 did not induce the pH3⁺ signal in the defined food, likely due to the absence of baseline VB5 present in cornmeal-based standard food. Consistently, the induction of pH3⁺ signal was diminished at the higher dose of 10 mM VB5. Together, these results indicate a dose-dependent effect of dietary VB5 supplementation on cell proliferation. Accordingly, we used 2.5 mM VB5 supplementation for subsequent experiments.

**Fig. 1.**
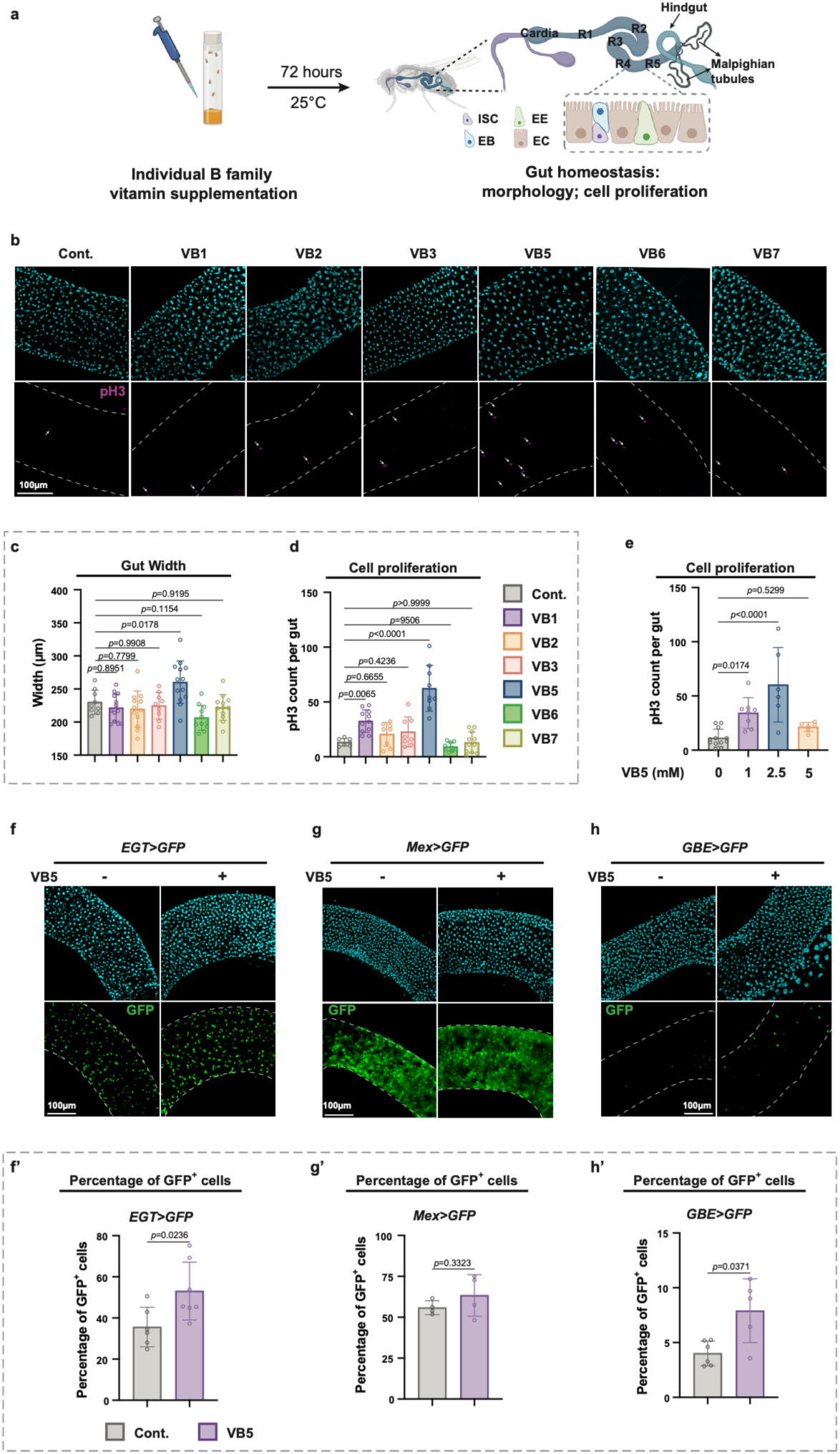
VB5 supplementation stimulates intestinal stem cell ISC proliferation. **a**, Experimental design for individual B vitamin supplementation and assessment of gut homeostasis. The *Drosophila* midgut is composed of five regions (R1-R5) with distinct cell types. The Malpighian tubules (MTs) are connected to the posterior midgut. ISC, intestinal stem cell; EB, enteroblast; EC, enterocyte; EE, enteroendocrine cell. **b,** Representative images of midguts from flies supplemented with VB1, VB2, VB3, VB5, VB6, VB7, and control (Cont.). Vitamin concentrations were determined based on those used in *Drosophila* holidic medium protocols (see **Methods** and **Supplementary Table 1**). In the pH3 panel, the guts are outlined with dashed lines, and pH3⁺ signals are indicated by arrows. **c,** Quantification of midgut width from flies with individual B vitamin dietary supplementation. Each dot represents one midgut. n = 9, 13, 13, 12, 9, 11, and 12 in Cont., VB1, VB2, VB3, VB5, VB6, and VB7, respectively. **d,** Quantification of phospho-Histone 3-positive (pH3^+^) cells per midgut from flies with individual B vitamin dietary supplementation. Each dot represents one midgut. n = 9, 9, 11, 8, 8, 9, and 11 in Cont., VB1, VB2, VB3, VB5, VB6, and VB7, respectively. Both VB1 and VB5 increased pH3^+^ cell counts, with VB5 inducing the most robust response and thus selected for further investigation. **e,** Quantification of pH3^+^ cells per midgut from flies with VB5 dietary supplementation at 0, 1, 2.5, or 5 mM. n = 12, 8, 6, and 6 in 0 mM, 1 mM, 2.5 mM, and 5 mM, respectively. See also **Fig. S1a** for dose-dependence examination using holidic medium. **f, g, h,** Representative images of guts showing GFP expression in ISC/EB (*EGT>*) (**f**), EC (*Mex>*) (**g**), and EB (*GBE>*) (**h**) cell populations. **f’, g’, h’,** Quantification of percentage of GFP^+^ cells from panel **f, g,** and **h**, respectively. n = 6 and 7 in VB5 (-) and VB5 (+) of **f’**, respectively. n = 5 and 4 in VB5 (-) and VB5 (+) of **g’**, respectively. n = 6 and 5 in VB5 (-) and VB5 (+) of **h’**, respectively. Statistical significance is assessed by ordinary one-way ANOVA (**c-e**) and unpaired two-sided Student’s *t*-test (**f’-h’**). Error bars indicate s.d., with means at the center. **a** created in BioRender. Miao, T. (2026) https://BioRender.com/hpytzof. Source data are provided as a Source Data file.

ISCs maintain midgut homeostasis through differentiating into progenitor state enteroblasts (EBs), which then terminally differentiate into enterocytes (ECs) or enteroendocrine cells (EEs), respectively [34] **(Fig. 1a)**. To further investigate how VB5 affects gut tissue homeostasis, we followed the percentage of gut cell subtypes by using cell type-specific Gal4 drivers to express GFP: EGT-Gal4 (ISC/EB), Myo1A-Gal4 (EC), and Su(H)-GBE-Gal4 (EB) **(Fig. 1f-h)**. Notably, VB5 supplementation increased the proportion of ISCs and EBs, suggesting that VB5 enhances stem cell proliferation or self-renewal, thus stimulates differentiation toward early progenitor stages. Together, our data indicates that dietary VB5 induces midgut expansion by stimulating ISC proliferation, uncovering a previously unappreciated role for VB5 in modulating intestinal tissue homeostasis.

### CoA biosynthesis in the Malpighian tubules (MTs) non-autonomously regulates ISC proliferation

VB5 serves as a precursor in the *de novo* biosynthesis of CoA, an essential molecule in numerous fundamental biological processes [5–7]. The CoA biosynthetic pathway is highly conserved across species and known regulatory inputs primarily control the function of PANK1-3 and PANK4 [7, 11] **(Fig. 2a)**. In *Drosophila*, the single pantothenate kinase orthologous to PANK1-3 is encoded by the gene *Fumble* (*Fbl*) [38] while an ortholog for PANK4 has not been previously characterized in flies. Using amino acid sequence alignments of human PANK4 with the fly proteome, we identified CG5828 as a strong candidate ortholog of PANK4. Notably, CG5828 completely lacks the N-terminal pantothenate (pseudo) kinase domain of PANK4 from other species and consists predominantly of a DUF89 domain, sharing ∼45% identity / 61% similarity with the metabolite phosphatase domain of human PANK4. This domain includes two conserved aspartate residues that coordinate the obligate metal cation and are essential for catalytic activity **(Fig. 2b and Fig. S1b)**. To determine whether CG5828 functions as an active metabolite phosphatase, we purified recombinant GST-tagged protein from bacteria **(Fig. S1c)** and subjected it to *in vitro* phosphatase assays. CG5828 exhibited robust phosphatase activity towards both the generic phospho-substrate para-nitrophenyl phosphate (PNPP) and PANK4’s proposed endogenous substrate, the CoA synthesis intermediate 4’-phosphopantetheine (p-PaSH). Importantly, mutation of the predicted catalytic aspartates (D209 or D245) abolished activity **(Fig. 2c)**. Consistent with human PANK4 and other DUF89 family members, CG5828 phosphatase activity requires a divalent metal cation, with activity robustly promoted by cobalt and nickel and to a lesser extent by magnesium and manganese. In contrast, other biologically relevant monovalent or divalent cations failed to support its enzymatic activity **(Fig. S1d)**. Based on the sequence and biochemical features, we designate CG5828 as the *Drosophila* functional ortholog of PANK4 (dPANK4).

**Fig. 2.**
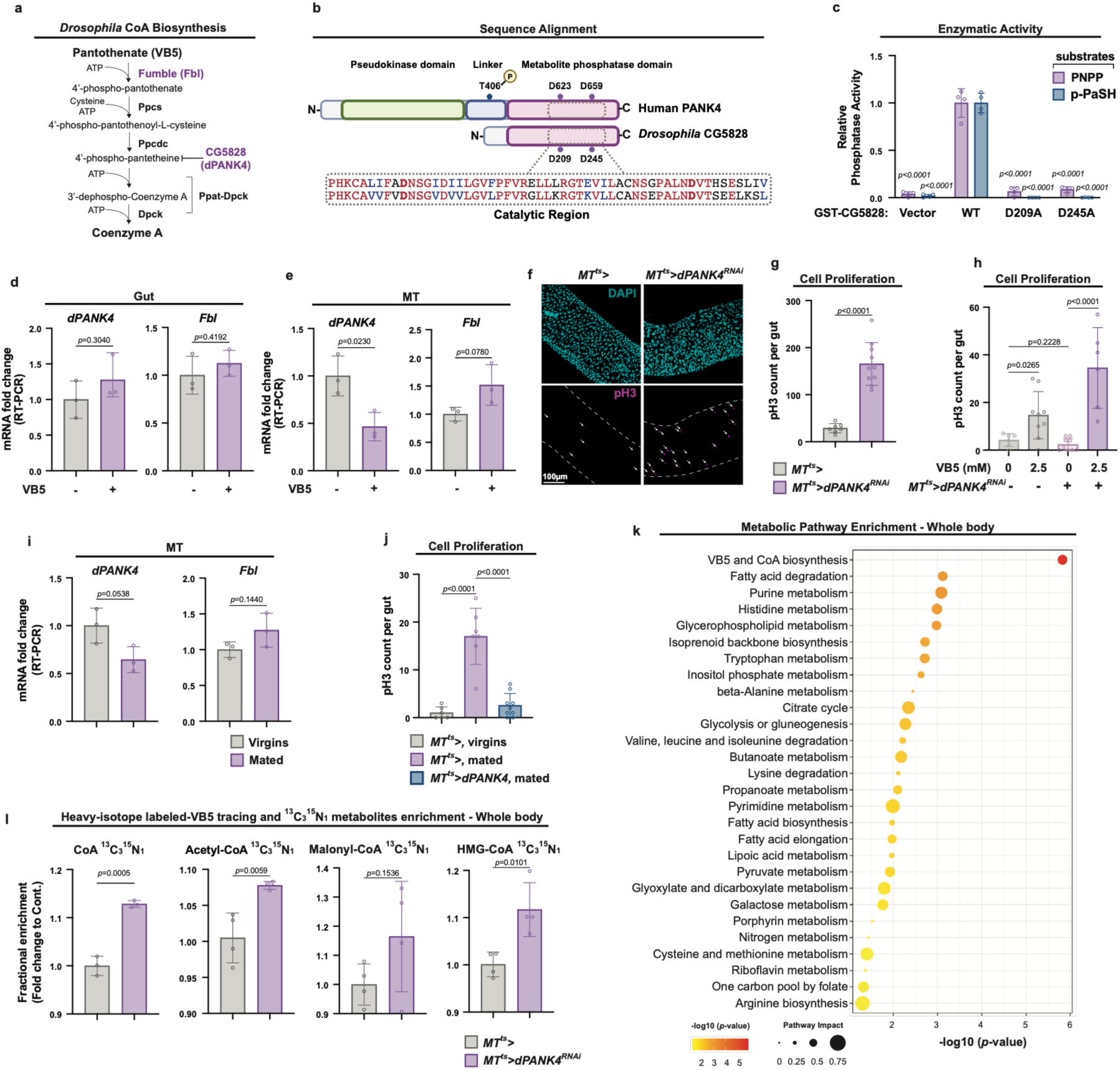
CoA biosynthesis in the Malpighian tubules (MTs) non-autonomously regulates ISC proliferation. **a**, Schematic of the CoA biosynthesis pathway in *Drosophila*. Fbl (Fumble), ortholog of mammalian PANK1–3; Ppcs, Phosphopantothenoylcysteine Synthetase; Ppcdc, Phosphopantothenoylcysteine Decarboxylase; Ppat, Phosphopantetheine Adenylyltransferase; Dpck, Dephospho-CoA Kinase. Ppat and Dpck activities are combined in a single bifunctional enzyme in many eukaryotes, known as Ppck-Dpck or COASY (Coenzyme A Synthase). **b,** Amino acid sequence alignment of the catalytic region between human PANK4 and *Drosophila* CG5828 (dPANK4). See also **Fig. S1b** for full-length alignment. **c**, CG5828/dPANK4 *in vitro* activity measurement showing phosphatase activity of GST-CG5828/dPANK4 (WT, D209A, or D245A) towards para-nitrophenyl phosphate (PNPP; 50mM) or 4′-phosphopantetheine (p-PaSH; 0.5mM). For statistical analysis, the relative activities of the GST vector control and GST-CG5828 mutants toward PNPP or p-PaSH are compared with those of WT GST-CG5828. n = 4. **d, e,** qRT-PCR analysis of *dPANK4* and *Fbl* mRNA levels in guts (**d**) and MTs (**e**) of flies with or without dietary VB5 supplementation. Each dot represents a biological replicate. n = 3. **f,** Representative images of guts in flies with or without MT-specific *dPANK4* knockdown at day 10. In the pH3 panel, the guts are outlined with dashed lines, and pH3⁺ signals are indicated by arrows. **g,** Quantification of pH3^+^ cells per midgut from **f**. n = 7 and 10 in *MT^ts^>* and *MT^ts^>dPANK4-RNAi*, respectively. **h,** pH3^+^ cell counts per midgut in flies with or without MT-specific *dPANK4* knockdown, fed on holidic diets with or without VB5 depletion. n = 7, 9, 10, and 6 in *MT^ts^>* (0 mM), *MT^ts^>* (2.5 mM), *MT^ts^>dPANK4-RNAi* (0 mM), and *MT^ts^>dPANK4-RNAi* (2.5 mM), respectively. **i,** qRT-PCR analysis of *dPANK4* and *Fbl* mRNA levels in MTs from virgin or mated female flies. n = 3. **j,** Quantification pH3^+^ cells per midgut in virgin or mated female flies with or without MT *dPANK4* overexpression at day 7. n = 6, 7, and 10 in *MT^ts^>* (virgins), *MT^ts^>* (mated), and *MT^ts^>dPANK4-RNAi* (mated), respectively. See also **Fig. S1l, m** for representative gut images for panel **j** and validation of this phenotype with independent fly lines. **k,** Pathway enrichment analysis of whole-fly metabolomics profiling of flies with MT-specific *dPANK4* knockdown versus controls at day 9. The top enriched KEGG pathways are shown, ranked by statistical significance (-log10(*p*-value)) and colored accordingly. Dot size represents pathway value. n = 5. **l,** Relative levels of [^13^C^15^N]-labeled CoA, acetyl-CoA, malonyl-CoA, and HMG-CoA in MT-specific *dPANK4* knockdown versus controls at day 9. n = 4. Statistical significance is assessed by unpaired two-sided Student’s *t*-test (**d-e, g, i, l**), one-way ANOVA (**c, h, j**), and two-tailed Fisher’s exact test (**k**). Error bars indicate s.d., with means at the center. **b** created in BioRender. Miao, T. (2026) https://BioRender.com/416o6v6. Source data are provided as a Source Data file.

To determine whether the gut CoA biosynthetic pathway contributes to the VB5-induced ISC proliferation, we first assessed the expression of *Fbl* and *dPANK4* in the gut following dietary VB5 supplementation and found that their transcript levels remained unchanged **(Fig. 2d)**. Next, we knocked down *Fbl* in the gut using the temperature-sensitive EGT-Gal4 (*EGT^ts^>*) or Myo1A-Gal4 (*Mex^ts^>*) driver to test whether gut-intrinsic CoA biosynthesis is required for VB5-induced ISC proliferation. Intriguingly, *Fbl* knockdown in neither ISC/EBs nor ECs suppressed VB5-induced ISC proliferation **(Fig. S1e)**. In contrast, these flies exhibited elevated baseline level of pH3^+^ signal, and VB5 supplementation further increased the ISC proliferation. This elevated baseline proliferation likely reflects metabolic imbalance and gut damage caused by loss of Fbl function in the gut. In addition, our previous whole-body single-nucleus RNA-seq (snRNA-seq) analysis revealed that VB5 transporter *Smvt* (the *Drosophila* homolog of SLC5A6/SMVT), which is known to stimulate CoA biosynthesis [17], was minimally expressed in the ISCs and ECs **(Fig. S2f)** [39]. Collectively, these results suggest that gut-intrinsic CoA biosynthesis may not be required for VB5-induced ISC proliferation.

To determine whether CoA biosynthesis in another tissue mediates VB5-induced ISC proliferation in a non-autonomous manner, we examined the expression pattern of CoA biosynthetic genes across *Drosophila* tissues by analyzing the reference single-cell transcriptomic database [40]. Although *Fbl* expression is enriched broadly across various tissues, including fat body, oenocytes, heart, gut, and Malpighian tubules (MTs; functionally analogous to human kidney), *dPANK4* expression is predominantly enriched in MTs, particularly in principal cells (**Fig. S1g-h)** [41].Since PANK4 acts as a negative regulator of CoA biosynthesis, the relatively high *dPANK4* expression in the MTs suggests that this tissue may harbor a high intrinsic capacity for CoA biosynthesis that is restrained under certain physiological conditions. Supporting this hypothesis, VB5 supplementation significantly downregulated *dPANK4* and upregulated *Fbl* expression in the MTs **(Fig. 2e)**, suggesting increased CoA biosynthesis in this tissue. To test whether MT-specific CoA biosynthesis influences ISC proliferation, we knocked down *dPANK4* using the temperature-sensitive, MT principal cell-specific driver (hereafter referred to as *MT^ts^>*) [42]. Indeed, *dPANK4* knockdown in the MTs led to more than fivefold increase in ISC proliferation, as indicated by pH3^+^ cell counts **(Fig. 2f-g and Fig. S1i)**. This effect was consistently observed in both female and male flies **(Fig. S1j)** and validated using two additional independent RNAi lines **(Fig. S1k)**. Importantly, the ISC proliferation phenotype was dependent on VB5 availability, as *MT^ts^>dPANK4^RNAi^* flies fed on a VB5-deprived diet did not exhibit increased ISC proliferation **(Fig. 2h)**. Together, our data suggests that dietary VB5 supplementation activates CoA biosynthesis in the MTs, which in turn non-autonomously drives ISC proliferation in the gut.

Building on these findings, we hypothesized that CoA biosynthesis in the MT may be activated under specific physiological conditions to support gut homeostasis and renewal. To test this, we examined the effects of mating, a well-established physiological stimulus known to induce ISC proliferation in female flies [43, 44]. Intriguingly, mating significantly downregulated the expression of *dPANK4* in the MTs **(Fig. 2i)**, suggesting that mating promotes CoA production by relieving dPANK4-mediated inhibition. Supporting this, overexpression of *dPANK4* in the MTs, which suppresses CoA production, completely abolished mating-induced ISC proliferation **(Fig. 2j and Fig. S1l-m)**. Altogether, these data indicate that CoA biosynthesis in MTs is required to maintain gut homeostasis and renewal in response to both dietary VB5 supplementation and physiological stimulation.

To further access the systemic impact of MT CoA biosynthesis, we performed targeted liquid chromatography-coupled tandem mass spectrometry (LC-MS/MS) based metabolomics analyses on whole-body samples from flies with and without MT-specific *dPANK4* knockdown. Importantly, pathway enrichment analysis identified VB5 and CoA biosynthesis as the most significantly affected metabolic pathway **(Fig. 2k)**. Consistently, VB5 levels were significantly reduced in *dPANK4* knockdown flies **(Fig. S1n)**, suggesting altered CoA synthesis. To directly measure CoA biosynthetic flux, we fed flies with heavy isotope(^13^C_3_^15^N_1_)-labeled VB5 and performed LC-MS/MS-based isotopic tracing metabolomics to monitor the incorporation of labeled VB5 into CoA and its derivatives. Notably, we observed a significant increase in the fractional abundance of labeled CoASH (unesterified CoA) and several key acyl-CoAs including acetyl-CoA, malonyl-CoA, and HMG-CoA in whole *MT^ts^>dPANK4^RNAi^*flies **(Fig. 2l)**, indicating a conserved role of dPANK4 in suppressing *de novo* CoA biosynthesis. These findings reveal that the renal system as a central site for CoA production in flies. Collectively, our results demonstrate that VB5 supplementation or physiological stimuli can activate CoA biosynthesis specifically in the MTs, which in turn promotes ISC proliferation in the gut. Our findings identify dPANK4 as a key regulator of CoA biosynthesis *in vivo* and highlight the MT as a surprisingly influential site of CoA biosynthesis, linking renal CoA production to whole-body metabolic coordination and gut homeostasis.

### MT CoA production promotes gut MVA and isoprenoid backbone biosynthesis

CoA is a central cofactor involved in numerous cellular processes, including *de novo* lipogenesis and protein acetylation, through the function of its key derivative, acetyl-CoA [18, 20, 45]. We therefore investigated whether activation of CoA production in the MTs influences gut homeostasis by altering these downstream pathways. We first measured the expression of essential lipogenic genes in the gut of *MT^ts^>dPANK4^RNAi^* flies. However, transcription levels of *Acly* (ATP citrate lyase), *AcCoAS* (acetyl-CoA synthetase), *Acc* (acetyl-CoA carboxylase), and *FASN1* (fatty acid synthase 1) were unchanged **(Fig. S2a)**. Consistent with this, BODIPY lipid staining showed no increase in lipid droplet accumulation in the gut of *MT^ts^>dPANK4^RNAi^* flies **(Fig. S2b-c)**. These observations suggest that fatty acid synthesis may not be activated in the gut upon increased CoA biosynthesis in the MTs. We next examined protein acetylation, a post-translational modification derived from acetyl-CoA that modulates a wide range of cellular functions [45]. Protein acetylation is catalyzed by lysine acetyltransferases (KATs), which transfer acetyl groups from acetyl-CoA to lysine residues on target proteins [46]. Although we observed increased expression of several KAT homologs in the guts of *MT^ts^>dPANK4^RNAi^* flies, including *Mof* (*KAT8*), *Hat1*, and *Gcn5* **(Fig. S2d)**, knocking down these genes in the gut did not block VB5-induced ISC proliferation **(Fig. S2e)**, suggesting that protein acetylation is unlikely to be the primary driver of this response.

To further investigate how CoA biosynthesis in the MTs influences gut homeostasis, we performed targeted metabolomics on gut samples from MT-specific *dPANK4*-knockdown flies. Pathway enrichment analysis identified several impacted metabolic pathways, including thiamine metabolism, arginine and proline metabolism, and isoprenoid backbone biosynthesis **(Fig. 3a)**. Intriguingly, our heavy isotope-labeled VB5 tracing and metabolomics analysis revealed that *dPANK4* knockdown enhanced labeled VB5 incorporation into HMG-CoA in whole animals **(Fig. 2l)**, indicating that HMG-CoA biosynthesis was promoted. Given that HMG-CoA is primarily used for MVA production [24], the key entry point for isoprenoid backbone synthesis **(Fig. 3b)**, we hypothesized that these pathways may be activated in the gut in response to enhanced CoA production in the MTs. To test this hypothesis, we examined the expression of genes encoding key enzymes in the MVA and isoprenoid backbone biosynthesis pathways, including *Hmgcr*, *Fpps* (fly homolog of *FDPS*), and *qm* (*quemao*, fly homolog of *GGPS1*) **(Fig. 3b)**. Indeed, all three genes were upregulated in the guts of *MT^ts^>dPANK4^RNAi^* flies **(Fig. 3c)**. Additionally, the expression of *β-GGT-I* (β subunit of type I geranylgeranyl transferase, fly homolog of *PGGT1B*), an enzyme mediating protein prenylation utilizing GGPP derived from isoprenoid backbone biosynthesis pathway, was also elevated **(Fig. 3b-c)**. These results suggest that the MVA and isoprenoid backbone biosynthesis is activated in the gut in response to increased CoA biosynthesis in the MTs. To test whether MVA pathway acts downstream of MT CoA biosynthesis to drive ISC proliferation, we fed flies with or without MT-specific *dPANK4* depletion a diet supplemented with simvastatin, a well-established MVA pathway inhibitor **(Fig. 3b)** [47]. Supporting our hypothesis, simvastatin treatment significantly suppressed the increased pH3⁺ signal in the gut of *MT^ts^>dPANK4^RNAi^* flies **(Fig. 3d and Fig. S2f)**, suggesting that MVA pathway activity is required for ISC proliferation induced by CoA biosynthesis in the MTs. To further validate this, we genetically disrupted the MVA and isoprenoid biosynthesis pathways in the gut by knocking down *Hmgcr* or *qm* and assessed ISC proliferation in response to VB5 supplementation. Intriguingly, the results showed that VB5-induced ISC proliferation was completely blocked by knockdown of either gene **(Fig. 3e)**, demonstrating that both MVA and isoprenoid backbone biosynthesis in the gut are essential for this effect.

**Fig. 3.**
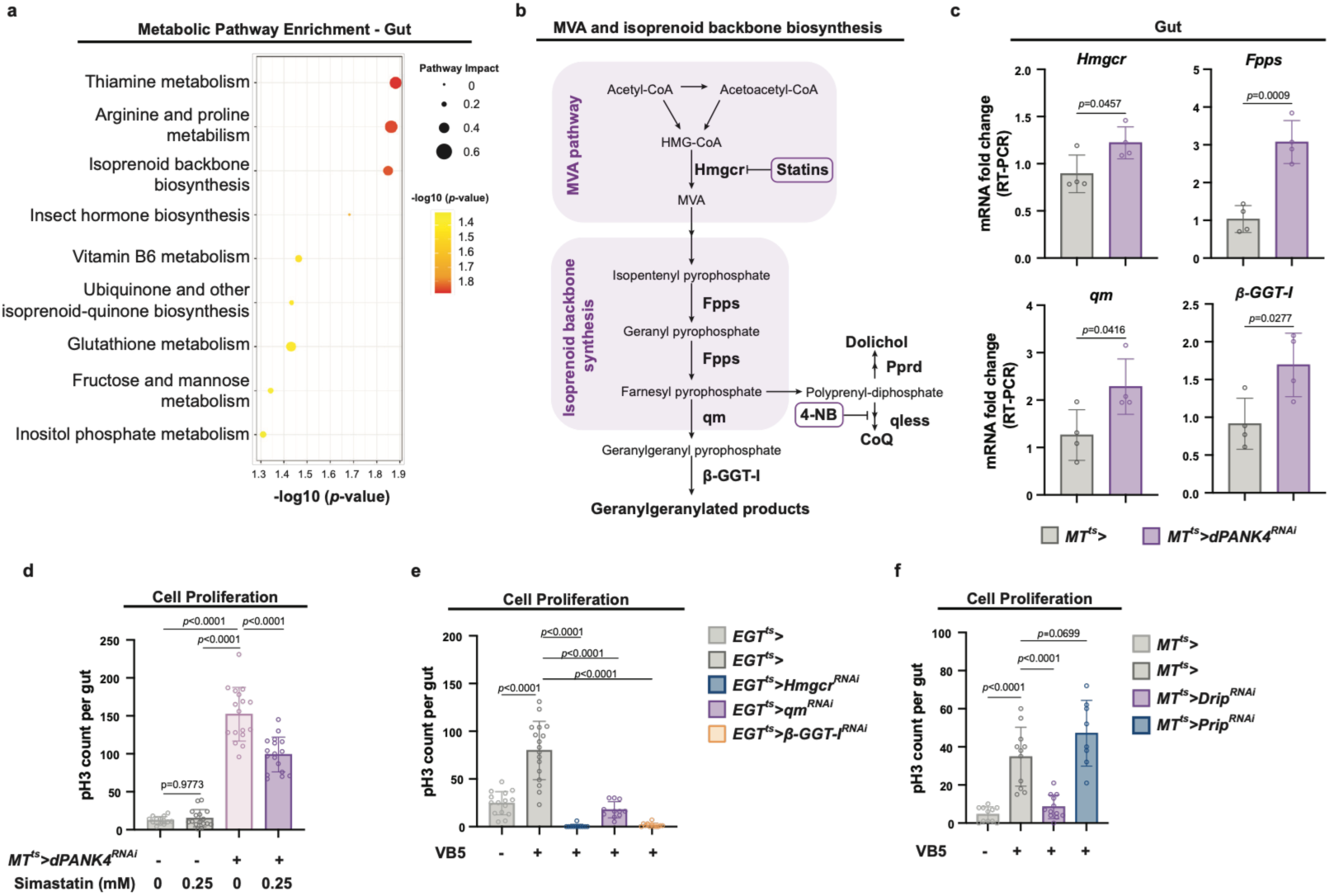
MT CoA production promotes gut mevalonate (MVA) and isoprenoid backbone biosynthesis. **a**, Pathway enrichment analysis of gut metabolomics profiling of flies with MT-specific *dPANK4* knockdown versus controls at day 9. The top enriched KEGG pathways are shown, ranked by statistical significance (-log10(*p*-value)) and colored accordingly. Dot size represents pathway value. n = 4. **b,** Schematic of mevalonate (MVA) and isoprenoid biosynthesis pathway in *Drosophila*. Hmgcr, HMG-CoA reductase; Fpps, Farnesyl pyrophosphate synthase; qm, Quasimodo, *Drosophila* homolog of geranylgeranyl diphosphate synthase (GGPS1); β-GGT-I, β subunit of Geranylgeranyltransferase type I (GGTase-I). **c,** qRT-PCR analysis of *Hmgcr*, *Fpps*, *qm*, and *β GGT-I* mRNA levels in guts from MT-specific *dPANK4* knockdown and control flies at day 9. n = 4. **d,** Quantification of pH3^+^ cells per midgut in flies with or without MT *dPANK4* depletion, fed on DMSO or simvastatin-containing (0.25 mM) diet. n = 16, 16, 18, and 17 in *MT^ts^>* (0 mM), *MT^ts^>* (0.25 mM), *MT^ts^>dPANK4-RNAi* (0 mM), and *MT^ts^>dPANK4-RNAi* (0.25mM), respectively. See also **Fig. S2f** for effect of 2.5mM simvastatin treatment. **e,** pH3^+^ cell quantification in control flies or flies with gut-specific knockdown of *Hmgcr*, *qm*, or *β-GGT-I* under VB5 supplementation. n = 15, 17, 12, 11, and 11 in *EGT^ts^>* [VB5 (-)], *EGT^ts^>* [VB5 (+)], *EGT^ts^>Hmgcr-RNAi* [VB5 (+)], *EGT^ts^>qm-RNAi* [VB5 (+)], and *EGT^ts^> β-GGT-I-RNAi* [VB5 (+)], respectively. **f,** pH3^+^ cell quantification per midgut in control flies and flies with MT principal cell-specific *Drip* or *Prip* knockdown under VB5 supplementation. n = 13, 11, 12, and 8 in *MT^ts^>* [VB5 (-)], *MT^ts^>* [VB5 (+)], *MT^ts^>Drip-RNAi* [VB5 (+)], and *MT^ts^>Prip-RNAi* [VB5 (+)], respectively. Statistical significance is assessed by two-tailed Fisher’s exact test (**a**), unpaired two-sided Student’s *t*-test (**c**), and one-way ANOVA (**d-f**). Error bars indicate s.d., with means at the center. Source data are provided as a Source Data file.

We next explored the downstream mechanisms of MVA-isoprenoid pathway activity. Although cholesterol biosynthesis is a well-characterized output of the MVA-isoprenoid pathway implicated in cell proliferation and tumor progression in mammals, this branch is absent in insects [25, 48]. We therefore tested for effects on protein prenylation, which relies on isoprenoid intermediates and is mediated by geranylgeranyl transferase*s* including *β-GGT-I*. Consistently, gut-specific knockdown of *β-GGT-I* completely abolished VB5-induced ISC proliferation **(Fig. 3e)**. Previous studies have shown that β-GGT-I-mediated prenylation activates Rho GTPases, which are required for the activation of the Hippo pathway mediators YAP and TAZ (YAP/TAZ) and thereby promote cell proliferation and tissue overgrowth in *Drosophila* [49]. To test whether YAP/TAZ signaling is activated upon enhanced renal CoA production, we examined the expression of two well-established target genes of Yorkie (Yki; the *Drosophila* YAP/TAZ ortholog), *Diap1* and *expanded* (*ex*) [49, 50], in the gut of *MT^ts^>dPANK4^RNAi^* flies. Consistent with prior studies showing downregulation of these genes upon inhibition of β-GGT-I-mediated prenylation [49], we observed significant upregulation of both *Diap1* and *ex* **(Fig. S2g)**. These findings suggest that prenylated signaling proteins, such as Rho GTPases, may be activated in the gut in response to increased isoprenoids output driven by renal CoA biosynthesis, leading to enhanced YAP/TAZ activity and ISC proliferation. We also tested the potential involvement of CoQ and dolichol biosynthesis, two additional major downstream outputs of MVA-isoprenoid pathway **(Fig. 3b)**. Dietary supplementation with 4-nitrobenzoic acid (4-NB), a competitive inhibitor of CoQ biosynthesis **(Fig. 3b)** [51] significantly suppressed the elevated pH3⁺ signal in the gut of *MT^ts^>dPANK4^RNAi^* flies **(Fig. S2h-i)**, suggesting that CoQ biosynthesis is required for ISC proliferation induced by enhanced renal CoA production. Consistently, gut-specific knockdown of *qless* (fly ortholog of *PDSS1*), a CoQ biosynthesis enzyme, completely abolished VB5-induced ISC proliferation **(Fig. S2j)**. Because no specific pharmacological inhibitor of dolichol biosynthesis is available, we instead genetically perturbed this pathway by knocking down *Pprd* (*Polyprenal reductase*, fly homology of *SRD5A3*). Inhibition of dolichol biosynthesis in the gut modestly attenuated VB5-induced ISC proliferation **(Fig. S2j)**. Collectively, these results demonstrate that MT-derived CoA pool influences gut homeostasis through the MVA-isoprenoid biosynthesis pathway. Multiple downstream outputs of this pathway, including Rho prenylation-mediated YAP/TAZ activation, as well as CoQ and dolichol biosynthesis, are likely engaged upon increased isoprenoid production and contribute to the stimulation of ISC proliferation.

A remaining question is how activation of the MVA-isoprenoid pathways in the gut is mechanistically coupled to CoA biosynthesis in the MT. Since CoA itself is not transported across the plasma membrane [5], we considered the possibility that CoA biosynthesis intermediates or metabolites produced in a CoA-dependent manner are transported from the MT to the gut, where they fuel MVA-isoprenoid biosynthesis and promote ISC proliferation. While inter-organ metabolite exchange often occurs via the circulatory system (hemolymph in *Drosophila*), the MTs and gut are anatomically connected **(Fig. 1a)**, raising the possibility of direct metabolite transfer via the shared lumen. Supporting this, we observed a spatially patterned increase in pH3^+^ cells in the midgut of flies with MT-specific knockdown of *dPANK4*. These mitotic signals first appeared in the posterior R4-R5 region, which lies adjacent to the MT-gut junction, and progressively spread anteriorly upon prolonged *dPANK4* knockdown **(Fig. 1a and Fig. S2k-l)**. This spatially restricted onset of proliferation and non-uniform distribution of mitotic cells suggest that the pro-proliferative signal may be locally transferred from the MTs to the gut, rather than disseminated through the hemolymph. A recent study reported that direct fluid exchange between the MTs and gut in *Drosophila* is mediated by aquaporins such as Drip [52]. The two well-established renal aquaporins, *Drip* and *Prip*, are expressed in both stellate cells and principal cells of the MTs **(Fig. S2m)** where they promote water influx through the renal tubules [41, 53]. To determine whether aquaporin-mediated water influx is involved in the inter-organ metabolite exchange, we knocked down *Drip* or *Prip* in stellate cells (*tsh^ts^>*) or principal cells (*MT^ts^>*). Knockdown of either aquaporin in these MT cell types did not affect baseline ISC proliferation (**Fig. S2n-o).** In line with the role of CoA biosynthesis activation in principal cells in driving ISC proliferation **(Fig. 2f-h and Fig. S1i-k)**, knockdown of *Drip* in principal cells completely abolished VB5-induced ISC proliferation **(Fig. 3f)**. In contrast, knockdown of *Drip* in stellate cells had no effect **(Fig. S2o)**. However, knockdown of *Prip* in neither stellate cells nor principal cells blocked VB5-induced ISC proliferation **(Fig. 3f; Fig. S2p)**. These observations are consistent with previous reports showing that Drip, rather than Prip, mediates renal-gut countercurrent flow that delivers renal-derived cytokines to the gut [52]. Altogether, these results suggest that CoA-dependent metabolites could reach the gut from the MTs through Drip-mediated water influx, which facilitates bulk fluid flow and enhances delivery, thereby activates intestinal MVA and isoprenoid backbone biosynthesis pathways to promote ISC proliferation.

### MT CoA biosynthesis promotes gut tumor growth

Local CoA biosynthesis has been implicated in oncogenic metabolism and tumor progression [11, 17], yet the role of host CoA metabolism to tumor growth via inter-organ metabolite exchange remains unexplored. To investigate this, we employed a *Drosophila* gut tumor model driven by overexpression of a constitutively active form of the transcriptional co-activator and Hippo pathway effector *Yki* (*yki^[S3A]^*) in the ISCs (hereafter referred to as Yki flies) [31, 39]. We observed significant transcriptional changes in CoA biosynthesis genes, *Fbl* and *dPANK4*, specifically in the MT principal cells, but not in the gut ISCs/tumors or ECs **(Fig. 4a and Fig. S3a)**, suggesting a potential contribution of MT-derived CoA synthesis to gut tumor growth. Supporting this, our previous snRNA-seq analysis of Yki flies revealed elevated expression of the VB5 transporter *Smvt* in the MT principal cells but not in ISCs or ECs **(Fig. S3a)** [39]. This observation suggests that dietary VB5 is preferentially transported to the MTs for CoA production in the Yki model. To decipher the role of MT CoA biosynthesis in gut tumor growth, we used a dual binary expression system (UAS-Gal4 and LexAop-LexA) to simultaneously induce tumor in the gut and knock down *Smvt* in the MTs (*EGT^ts^>yki^[S3A]^, MT^ts^>Smvt^RNAi^*). Interestingly, *Smvt* knockdown in the MTs significantly suppressed tumor growth in Yki flies **(Fig. 4b)**. Further, we inhibited MT CoA biosynthesis through either knocking down *Fbl* or overexpressing *dPANK4* in Yki flies (*EGT^ts^>yki^[S3A]^, MT^ts^>Fbl^RNAi^* or *EGT^ts^>yki^[S3A]^, MT^ts^>dPANK4*). Both interventions significantly suppressed tumor growth, as evidenced by reduced gut width, decreased Yki-GFP signal, and lower mitotic (pH3^+^) cell counts **(Fig. 4c-f and Fig. S3b-i)**. Consistent with reduced tumor burden, inhibition of CoA biosynthesis in the MTs also ameliorated the bloating phenotype of Yki flies, a symptom of systematic fluid imbalance **(Fig. 4c, g-h and Fig. S3f, j)** [31], and extended their overall survival **(Fig. 4i)**. These findings indicate that MT CoA biosynthesis promotes tumor growth and contributes to systemic disease progression in this model.

**Fig. 4.**
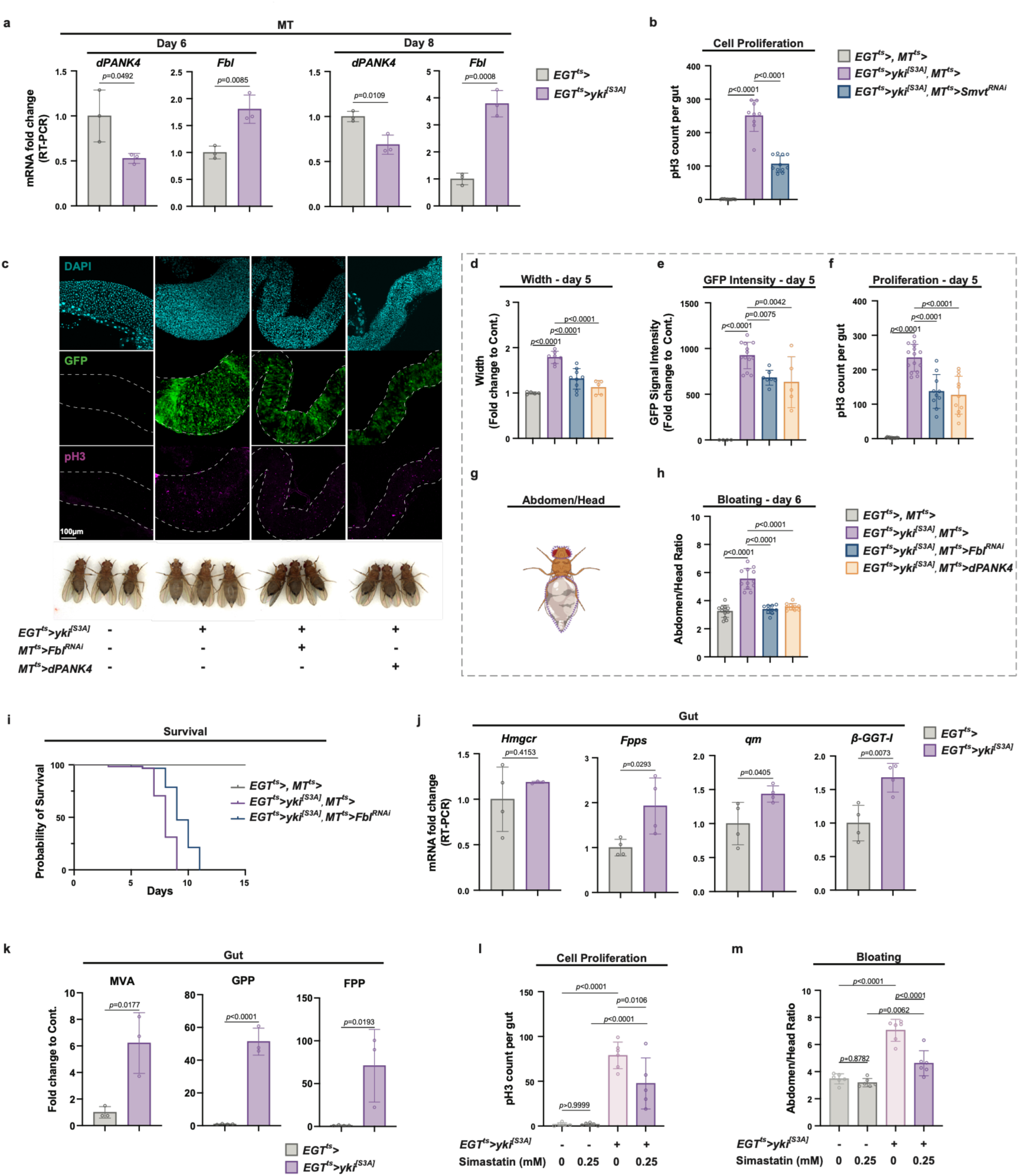
MT CoA biosynthesis promotes gut tumor growth. **a**, qRT-PCR analysis of *dPANK4* and *Fbl* mRNA levels in MTs from Yki and control flies at day 6 and 8. n = 3. **b,** pH3^+^ cell quantification in control flies and Yki flies with or without MT-specific *Smvt* knockdown at day 5. n = 13, 9, and 11 in *EGT^ts^>, MT^ts^>*, *EGT^ts^>yki^[S3A]^*, *MT^ts^>*, and *EGT^ts^>yki^[S3A]^, MT^ts^>Smvt-RNAi*, respectively. **c,** Representative images of guts (day 5) and representative images of bloating phenotypes (day 6) from control flies and Yki flies with or without MT-specific *Fbl* knockdown or *dPANK4* overexpression. **d-f,** Quantification of gut width (**d**), GFP intensity (**e**), and pH3^+^ cell counts (**f**) in (**c**). n = 6, 8, 9, and 5 in *EGT^ts^>, MT^ts^>*, *EGT^ts^>yki^[S3A]^*, *MT^ts^>*, *EGT^ts^>yki^[S3A]^, MT^ts^>Fbl-RNAi*, and *EGT^ts^>yki^[S3A]^, MT^ts^>dPANK4* in **d**, respectively. n = 6, 13, 7, and 5 in *EGT^ts^>, MT^ts^>*, *EGT^ts^>yki^[S3A]^*, *MT^ts^>*, *EGT^ts^>yki^[S3A]^, MT^ts^>Fbl-RNAi*, and *EGT^ts^>yki^[S3A]^, MT^ts^>dPANK4* in **e**, respectively. n = 15, 15, 10, and 11 in *EGT^ts^>, MT^ts^>*, *EGT^ts^>yki^[S3A]^*, *MT^ts^>*, *EGT^ts^>yki^[S3A]^, MT^ts^>Fbl-RNAi*, and *EGT^ts^>yki^[S3A]^, MT^ts^>dPANK4* in **f**, respectively. In the pH3 and GFP panels, the guts are outlined with dashed lines. See also **Fig. S3b-i** for phenotypes at day 3 and validations using independent fly lines. **g-h,** Quantification method (**g**) and results (**h**) for bloating phenotypes shown in (**c**), defined as abdomen-to-head area ratio. n = 12, 12, 11, and 12 in *EGT^ts^>, MT^ts^>*, *EGT^ts^>yki^[S3A]^*, *MT^ts^>*, *EGT^ts^>yki^[S3A]^, MT^ts^>Fbl-RNAi*, and *EGT^ts^>yki^[S3A]^, MT^ts^>dPANK4* in **h**, respectively. See also **Fig. S3b, j** for validation with independent lines. **i,** Survival curves of control, Yki, and Yki flies with MT *Fbl* depletion. N = 40-45. **j,** qRT-PCR analysis of *Hmgcr*, *Fpps*, *qm*, and *β GGT-I* mRNA levels in guts from Yki or control flies at day 6. n = 4. **k,** Relative metabolite levels of mevalonate (MVA), geranyl pyrophosphate (GPP), and farnesyl pyrophosphate (FPP) retrieved from gut metabolomics analysis of Yki and control flies at day 6 (n = 4). See also **Fig. S3k** for pathway enrichment analysis of gut metabolomics profiling. **l,** pH3^+^ cell quantification in control flies and Yki flies with or without simvastatin treatment (0.25 mM) at day 5. n = 7, 7, 12, 6, and 5 in *EGT^ts^>* (0 mM), *EGT^ts^>* (0.25 mM), *EGT^ts^> yki^[S3A]^* (0 mM), and *EGT^ts^> yki^[S3A]^* (0.25 mM), respectively. **m,** Quantification of bloating in control flies and Yki flies with or without simvastatin treatment at day 6. n = 6. Also see **Fig. S3l** for representative images of bloating phenotypes. Statistical significance is assessed by unpaired two-sided Student’s *t*-test (**a, j-k**), one-way ANOVA (**b, d-f, h, l-m)**, and log-rank (Mantel–Cox) test (**i**). Error bars indicate s.d., with means at the center. **g** created in BioRender. Miao, T. (2026) https://BioRender.com/6sataog. Source data are provided as a Source Data file.

We have demonstrated that MVA-isoprenoid axis functions downstream of MT CoA biosynthesis to promote ISC proliferation in the wild-type flies **(Fig. 3)**. We next asked whether tumor cell proliferation is similarly driven by the activation of the MVA-isoprenoid biosynthetic pathway. Supporting this hypothesis, we observed upregulation of key genes involved in these pathways, including *Fpps*, *qm*, and *β-GGT-I*, in the gut tumors **(Fig. 4j)**. Consistent with this transcriptional activation, targeted metabolomics analysis of the gut samples from Yki flies revealed significant enrichment of the isoprenoid backbone biosynthesis pathway **(Fig. S3k)**, along with notable accumulation of intermediate metabolites, including MVA, geranyl pyrophosphate (GPP), and farnesyl pyrophosphate (FPP) **(Fig. 4k)**, suggesting robust activation of the MVA-isoprenoid axis in the gut tumor. To further test whether MVA pathway activity is functionally required for tumor growth, we treated Yki flies with simvastatin, a specific inhibitor of MVA pathway. Consistent with our hypothesis, simvastatin treatment significantly reduced tumor cell proliferation **(Fig. 4l)**, ameliorated the associated bloating phenotype **(Fig. 4m and Fig. S3l)**, and extended their overall survival **(Fig. S3m)**. In summary, the results demonstrate that CoA biosynthesis is upregulated in the MTs of tumor-bearing flies and is required for intestinal tumor growth. Enhanced CoA production in the MTs correlates with activation of the MVA-isoprenoid axis in the gut which supports tumor cell proliferation, suggesting an inter-organ communication mechanism similar to that observed in wild-type flies. More broadly, these findings provide evidence for an MT/kidney-gut metabolite exchange as a shared regulatory mechanism underlying both gut tissue homeostasis and tumor growth.

### *Myc* transcriptionally regulates CoA biosynthesis genes in the MTs

We previously reported that the PI3K-AKT signaling axis regulates CoA synthesis through phosphorylation of PANK4 in mammalian cells [11]. However, the *Drosophila* PANK4 ortholog, dPANK4, lacks the conserved phosphorylation sites found in mammalian PANK4 **(Fig. 2b)**. Nevertheless, our data indicate that dPANK4 transcript levels are altered by several physiological and pathological conditions in *Drosophila* **(Fig. 2e, 2i, 4a)**, suggesting that *dPANK4* may be subject to transcriptional rather than post-translational regulation. To identify transcriptional regulators of CoA biosynthesis, we employed the Transcription Factor-to-Target Gene (TF2TG) prediction tool [54] to analyze the promoter and intron regions (up to 1.5 kb upstream) of six *Drosophila* CoA biosynthetic genes **(Fig. 2a)**. This analysis identified 250 candidate transcription factors (TFs). Among them, six TFs, including the well-characterized oncogenic factor Myc, were predicted to regulate all six genes **(Supplementary Table 2)**. Notably, MYC has previously been linked to CoA metabolism in mammals through its regulation of the VB5 transporter SLC5A6/SMVT [17]. We have shown that CoA biosynthesis pathway is transcriptionally activated in the MTs of Yki flies **(Fig. 4a and Fig. S3a)**. Consistently, analysis of MT snRNA-seq data revealed that *Myc* expression was elevated in the principal cells and stellate cells in the MTs of Yki flies **(Fig. S4a)** [42]. In addition, TF regulon analysis identified Myc as the most significantly activated TF in the MTs of Yki flies **(Fig. S4b)** [39]. Notably, *Myc* transcription was not induced in ISCs or ECs in the guts of Yki flies **(Fig. S4c)** [39], in agreement with the unchanged expression of CoA biosynthesis genes in the gut **(Fig. S3a)**. These findings suggest that Myc may modulate transcription of CoA biosynthesis genes in the MTs to support increased CoA demand in tumor expansion. Consistently, our data showed that *Myc* overexpression in the MTs robustly upregulated *Fbl* and repressed *dPANK4* expression **(Fig. 5a)**. Supporting a direct regulatory role, a published chromatin immunoprecipitation followed high-throughput sequencing (ChIP-Seq) dataset of fly larvae (ENCSR191VCQ) revealed potential Myc binding sites near *dPANK4* and *Fbl* loci **(Fig. 5b)**. To confirm the direct binding of *dPANK4* and *Fbl* by Myc in our condition, we expressed HA-tagged Myc (*MT^ts^>Myc-HA*) and performed ChIP-qPCR, which validated Myc occupancy at the promoter regions of both genes in the MTs **(Fig. 5c)**. These findings demonstrate that Myc directly regulates the expression of *dPANK4* and *Fbl* in the MTs, thereby modulating CoA biosynthesis.

**Fig. 5.**
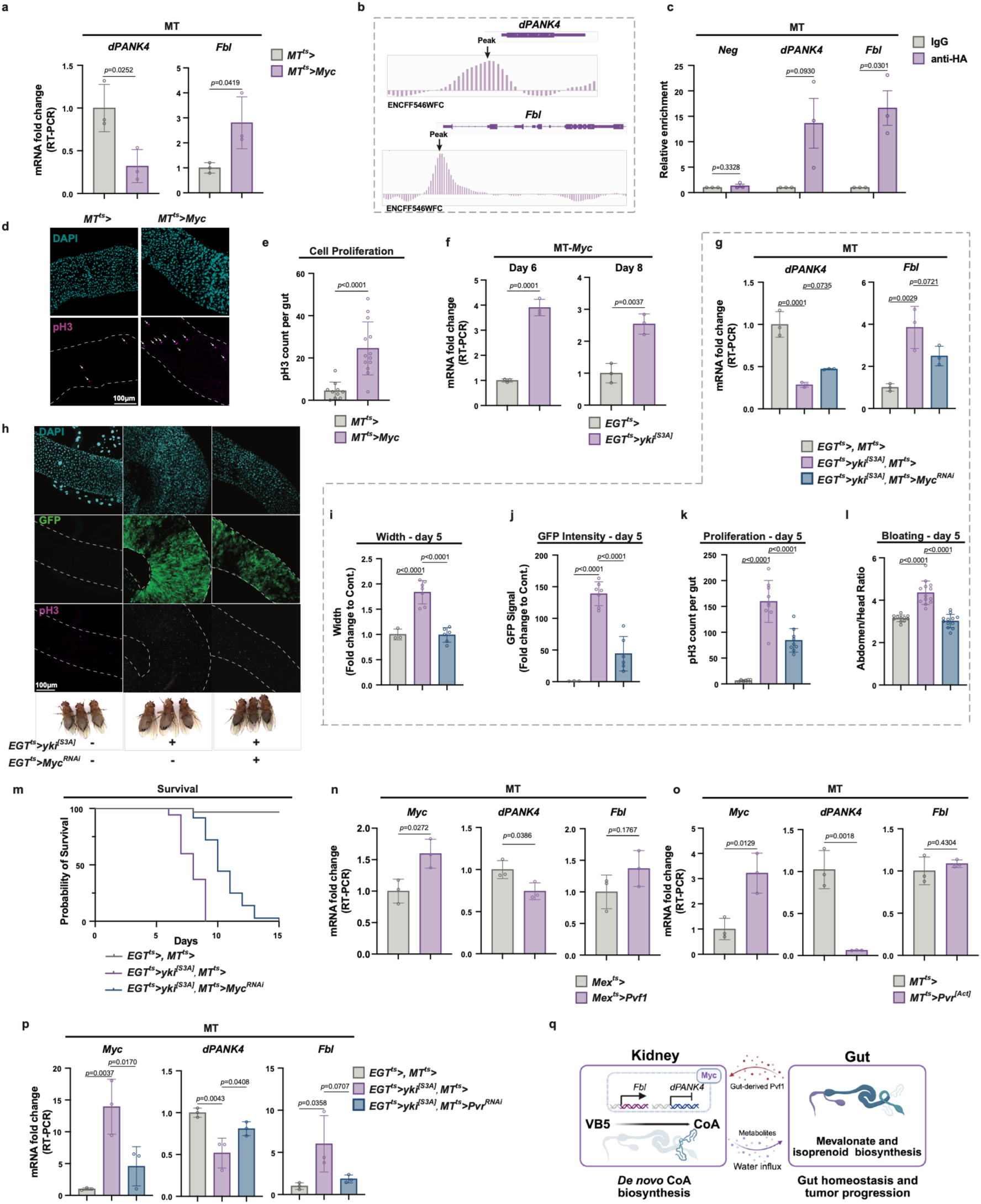
*Myc* transcriptionally regulates CoA biosynthesis genes in the MTs. **a**, qRT-PCR analysis of *dPANK4* and *Fbl* mRNA levels in MTs from *Myc*-overexpressing versus control flies at day 7. n = 3. **b,** Data retrieved from the modENCODE database indicating enrichment of Myc binding at the promoter regions of *dPANK4* and *Fbl*. **c,** ChIP-qPCR showing enrichment of HA-tagged Myc binding at the promoter regions of *dPANK4* and *Fbl* gene, shown by fold changes relative to control IgG (n = 3). Neg, negative control. **d-e,** Representative gut images (**d**) and pH3^+^ cell quantification (**e**) in *Myc*-overexpressing versus control flies at day 9. In the pH3 panel, the guts are outlined with dashed lines, and pH3⁺ signals are indicated by arrows. n = 10 and 13 in *MT^ts^>* and *MT^ts^>Myc*, respectively. **f,** qRT-PCR analysis of *Myc* mRNA level in MTs from Yki or control flies at day 6 and 8. n = 3. **g,** qRT-PCR analysis of *dPANK4* and *Fbl* mRNA levels in MTs from control and Yki flies with or without MT *Myc* depletion at day 6. n = 3. **h,** Representative gut tumor images (day 5) and representative images of bloating phenotypes (day 6) from control and Yki flies with or without MT-specific *Myc* knockdown. In the pH3 and GFP panels, the guts are outlined with dashed lines. **i–k,** Quantification of gut width (**i**), GFP intensity (**j**), and pH3^+^ cell counts (**k**) in (**h**). n = 3, 7, and 6 in *EGT^ts^>, MT^ts^>*, *EGT^ts^>yki^[S3A]^*,*MT^ts^>*, and *EGT^ts^>yki^[S3A]^, MT^ts^>Myc-RNAi* in **i**, respectively. n = 3, 7, and 6 in *EGT^ts^>, MT^ts^>*, *EGT^ts^>yki^[S3A]^*,*MT^ts^>*, and *EGT^ts^>yki^[S3A]^, MT^ts^>Myc-RNAi* in **j**, respectively. n = 7, 9, and 9 in *EGT^ts^>, MT^ts^>*, *EGT^ts^>yki^[S3A]^*,*MT^ts^>*, and *EGT^ts^>yki^[S3A]^, MT^ts^>Myc-RNAi* in **k**, respectively. See also **Fig. S4h-o** for phenotypes at day 3 and validations using independent fly lines. **l,** Bloating quantification via abdomen-to-head ratio. n = 11, 13, and 12 in *EGT^ts^>, MT^ts^>*, *EGT^ts^>yki^[S3A]^*,*MT^ts^>*, and *EGT^ts^>yki^[S3A]^, MT^ts^>Myc-RNAi* in **l**, respectively. See **Fig. S4l, p** for validation with independent lines. **m,** Survival curve of control and Yki flies with or without MT *Myc* knockdown. n = 40-45. **n,** qRT-PCR analysis of *Myc*, *dPANK4*, and *Fbl* mRNA levels in MTs from control and gut *Pvf1-*overexpressing flies. n = 3. **o,** qRT-PCR analysis of *Myc*, *dPANK4*, and *Fbl* mRNA levels in MTs from control and MT *Pvr* activation flies. n = 3. **p,** qRT-PCR analysis of *Myc*, *dPANK4*, and *Fbl* mRNA levels in MTs of control and Yki flies with or without MT *Pvr* depletion. n = 3. **q,** Summary model of the gut-kidney axis in *Drosophila*. Statistical significance is assessed by unpaired two-sided Student’s *t*-test (**a, c, e, f, n, o**), one-way ANOVA (**g, i-l, p)**, and log-rank (Mantel–Cox) test (**m**). Error bars indicate s.d., with means at the center. **q** created in BioRender. Miao, T. (2026) https://BioRender.com/62ab5bx. Source data are provided as a Source Data file.

Given that increased MT CoA production drives ISC proliferation **(Fig. 2)**, we next tested whether *Myc* overexpression could recapitulate this effect. Indeed, *Myc* overexpression in the MTs induced ISC proliferation in the gut **(Fig. 5d-e)**. Because *Myc* overexpression of can trigger apoptosis in certain conditions [55], it is possible that MT apoptosis releases stress signals to the gut and in turn drives ISC proliferation. We therefore examined the expression of canonical apoptosis-inducible genes in the MTs, including *reaper* (*rpr*), *hid*, *grim*, and *sickle* (*skl*), which are transcriptionally activated in response to diverse pro-apoptotic stimuli [56]. We found that the expression of *rpr* and *hid* was unchanged upon *Myc* overexpression, whereas *grim* expression was reduced **(Fig. S4d)**. *skl* was undetectable in the MTs of either control or *Myc*-overexpressing flies. These data suggest apoptosis is not induced by *Myc* overexpression in the MTs in our experiments, supporting the model that Myc promotes ISC proliferation primarily through activation of CoA biosynthesis rather than secondary tissue damage. Furthermore, we found that *Myc* expression is upregulated in the MTs of mated females **(Fig. S4e)**, consistent with the increased CoA biosynthetic activity following mating **(Fig. 2i)**. Furthermore, MT-specific knockdown of *Myc* substantially blocked mating-induced ISC proliferation **(Fig. S4f-g)**, suggesting that mating-induced CoA biosynthesis is mediated by Myc. Altogether, these results establish Myc as a key transcriptional regulator of CoA biosynthesis in the MTs in wild-type flies.

We next examined whether Myc also mediates CoA biosynthetic pathway activation in the context of tumor growth. Notably, *Myc* expression was increased in the MTs of Yki flies **(Fig. 5f)**. Moreover, MT-specific knockdown of *Myc* in Yki flies (*EGT^ts^>yki^[S3A]^, MT^ts^>Myc^RNAi^*) partially restored *dPANK4* expression and suppressed *Fbl* levels **(Fig. 5g)**, suggesting that Myc drives CoA biosynthesis in the MTs of the tumor-bearing flies. Importantly, depletion of *Myc* in the MTs of Yki flies significantly reduced gut tumor cell proliferation, as evidenced by reduced gut width, decreased Yki-GFP intensity, and lower pH3^+^ cell counts **(Fig. 5h-k and Fig. S4h-o)**. Consistent with reduced tumor burden, *Myc* knockdown alleviated the bloating phenotype (**Fig. 5h, l and Fig. S4l, p)** and significantly extended the survival of tumor-bearing flies **(Fig. 5m)**. Together, these results demonstrate that Myc expression in the MTs contributes to gut tumor growth by promoting CoA biosynthesis.

We then sought to identify the upstream signaling pathways responsible for activating *Myc* expression in the MTs. Our previous work identified several tumor-secreted cytokines, including ImpL2, Upd3, and Pvf1, that disrupt insulin, JAK-STAT, and PDGF/VEGF signaling, respectively, in various host tissues in Yki flies [31, 39, 57, 58]. Notably, *Myc* transcription is known to be inducible by multiple pathways, including PDGF signaling [59]. In line with this, we recently reported that gut-derived Pvf1 activates PDGF/VEGF receptor (Pvr) mediated JNK-Jra signaling in the MTs of Yki flies [42]. To test if *Myc* expression is modulated by this axis in the MTs, we overexpressed *Pvf1* in the gut of wild-type flies. Intestinal *Pvf1* overexpression significantly increased *Myc* transcription while concomitantly repressing *dPANK4* expression in the MTs **(Fig. 5n)**. Consistently, overexpression of a constitutively active form of Pvr in the MTs (*MT^ts^>Pvr^[Act]^*) robustly increased *Myc* transcription and Myc transcription activity **(Fig. 5o and Fig. S5q)**, while downregulating *dPANK4* **(Fig. 5o)**. These results indicate gut-derived Pvf1 activates Pvr signaling in the MTs, with Myc acting downstream of Pvr to regulate CoA biosynthesis. To assess the relevance of this regulatory axis in the tumor context, we knocked down *Pvr* in the MTs of Yki flies (*EGT^ts^>yki^[S3A]^, MT^ts^>Pvr^RNAi^*). This manipulation reduced *Myc* expression and restored *dPANK4* and *Fbl* levels to near-baseline values **(Fig. 5p)**. Collectively, our findings demonstrate that Myc directly binds the promoter regions of key CoA biosynthesis genes in the MTs and orchestrates pathway activation under both physiological and tumor-associated conditions. In the Yki tumor setting, *Myc* is upregulated in the MTs in response to Pvr signaling, thereby reprogramming host metabolism to support tumor growth **(Fig. 5q)**.

### Conserved pathogenic mechanism in kidney cancer

We previously reported that PANK4 suppresses tumorigenesis by inhibiting CoA biosynthesis in a mammalian model **(Fig. 6a)** [11]. Our *Drosophila* studies further demonstrated that Myc enhances tumor growth by increasing CoA production specifically in the MT/kidney **(Fig. 5)**. To determine whether this MYC-CoA axis also operates in human cancer, we analyzed published ChIP-seq data from a mammalian cancer model [60] and found strong MYC occupancy at the promoters of all four PANK genes **(Fig. S5a)**. Notably, MYC binding to *PANK1-3* promoters is prominent in the pre-tumoral and tumor samples but absent in the matched normal tissue **(Fig. S5a)**, suggesting a tumorigenesis-related CoA biosynthesis induction by MYC. These observations are complemented by prior work that MYC enhances VB5 uptake in breast cancer to elevate CoA levels [17]. While these findings underscore a general oncogenic demand for CoA, whether the MYC-CoA axis is functionally relevant in kidney malignancy, an organ with inherently high CoA levels and strong PANK1 expression [7, 8, 61, 62], remains underexplored.

**Fig. 6.**
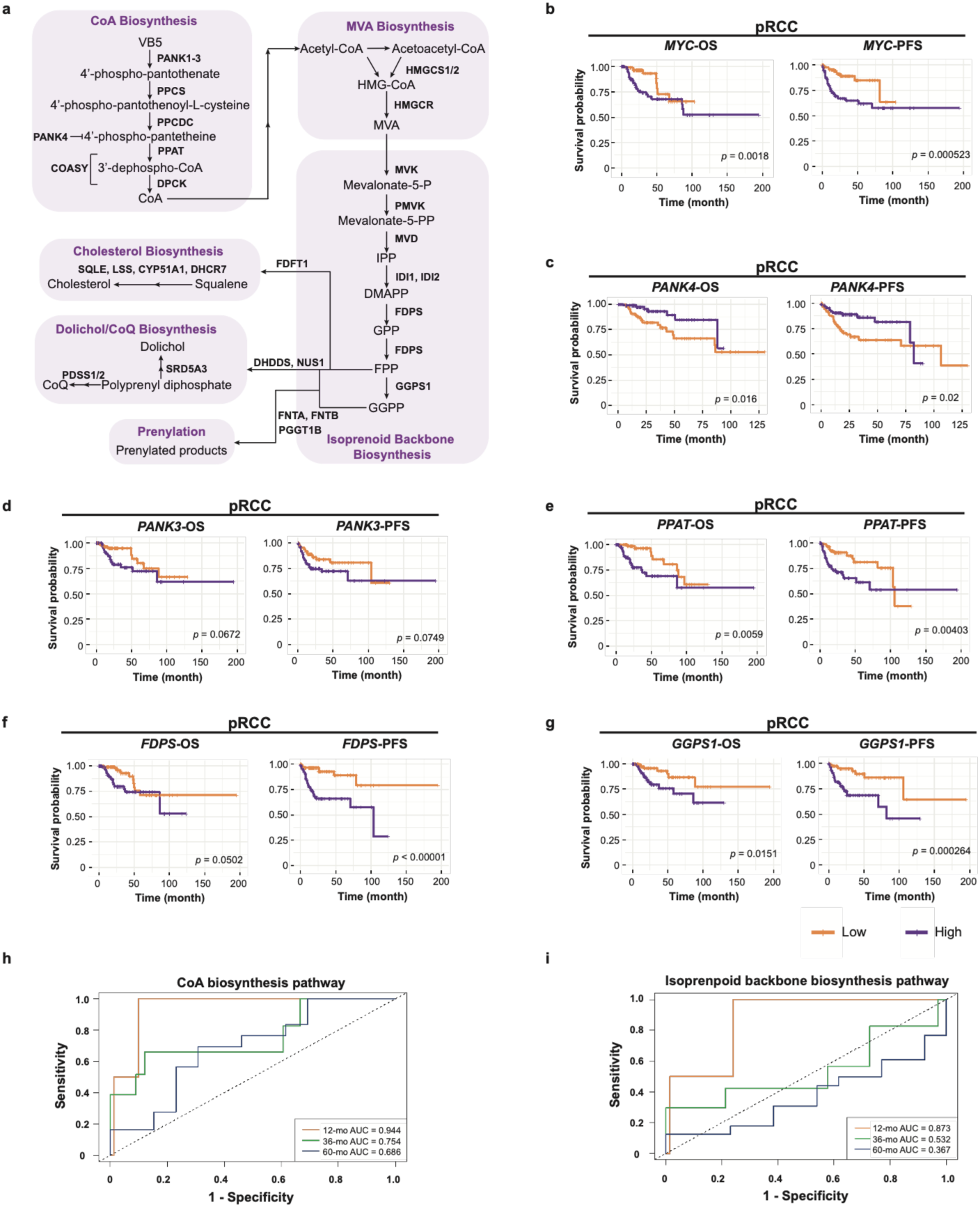
Conserved pathogenic mechanism in kidney cancer. **a**, Schematic of CoA, MVA, and isoprenoid backbone pathways and their downstream branches in humans, including cholesterol biosynthesis, dolichol and ubiquinone (CoQ) biosynthesis, and protein prenylation. HMGCS1/2, HMG-CoA synthase 1/2; HMGCR, HMG-CoA reductase; MVK, mevalonate kinase; PMVK, phosphomevalonate kinase; MVD, mevalonate diphosphate decarboxylase; IDI1/2, isopentenyl-diphosphate delta isomerase 1/2; FDPS, farnesyl diphosphate synthase; GGPS1, geranylgeranyl diphosphate synthase 1; FDFT1, farnesyl-diphosphate farnesyltransferase 1 (also known as squalene synthase); SQLE, squalene epoxidase; LSS, lanosterol synthase; CYP51A1, cytochrome P450 family 51 subfamily A member 1; DHCR7, 7-dehydrocholesterol reductase; DHDDS, dehydrodolichyl diphosphate synthase subunit; NUS1, Nogo-B receptor; PDSS1/2, decaprenyl-diphosphate synthase subunit 1/2; SRD5A3, steroid 5 alpha-reductase 3; FNTA, farnesyltransferase, alpha subunit; FNTB, farnesyltransferase, beta subunit; PGGT1B, protein geranylgeranyltransferase type I, beta subunit. IPP, isopentenyl pyrophosphate; DMAPP, dimethylally pyrophosphate; GPP, geranyl pyrophosphate; FPP, farnesyl pyrophosphate; GGPP, geranylgeranyl pyrophosphate. **b-g,** Kaplan-Meier (KM) survival analyses of overall survival (OS) and progression-free survival (PFS) for 280 Papillary Renal Cell Carcinoma (pRCC/KIRP) patients with low (bottom third) versus high (top third) expression of *MYC* (**b**), *PANK4* (**c**), *PANK3* (**d**), *PPAT* (**e**), *FDPS* (**f**), and *GGPS1* (**g**); the graph is based on data from the PanCancer Atlas of the TCGA. **h-i,** Time-dependent receiver operating characteristic (ROC) curve analysis showing the prognostic performance (area under the curve, AUC) of gene expression signatures from the CoA biosynthesis pathway (**h**) (n = 91) and the isoprenoid backbone synthesis pathway (**i**) (n = 91) in pRCC patients. AUC values were calculated at 12, 24, and 36 months of metastasis stage M0. Pathway gene sets are CoA biosynthesis: *PANK1-4, PPCS, PPCDC, PPAT* and *COASY*; isoprenoid backbone synthesis: *MVK, PMVK, MVD, IDI1, IDI2, FDPS* and *GGPS1*. Statistical significance is assessed by log-rank (Mantel–Cox) test (**b–g**) and AUC analysis (**h–i**). See also **Fig. S5-S6**. Source data are provided as a Source Data file.

Clear Cell Renal Cell Carcinoma (ccRCC) and Papillary Renal Cell Carcinoma (pRCC) are the two major subtypes of renal cell carcinomas (RCC) [63]. Compared to ccRCC, pRCC retains tubular architecture and is thus more structurally and functionally similar to the renal tubules [63], the mammalian counterpart of *Drosophila* MTs. To assess the clinical relevance of the kidney-intrinsic MYC-CoA axis, we interrogated gene expression and survival data from both ccRCC and pRCC patients in the TCGA PanCancer Atlas cohort [64, 65]. Intriguingly, Kaplan-Meier (KM) analyses showed that high *MYC* expression was significantly associated with poorer overall survival (OS) (*p* = 0.018) and progression-free survival (PFS) (*p* = 0.00052) in pRCC patients **(Fig. 6b)**, but not in ccRCC patients **(Fig. S5b)**, suggesting a pronounced pathogenic role for MYC in specifically in pRCC. Additionally, *MYC* expression showed a strong inverse correlation with PANK*4* expression in pRCC patients **(Fig. S5c)**, consistent with our *Drosophila* findings that Myc suppresses *dPANK4* to promote CoA biosynthesis. We next examined the association between CoA biosynthetic gene **(Fig. 6a)** expression and clinical outcomes. In pRCC, high expression of *PANK4*, a negative regulator of CoA biosynthesis, was significantly associated with improved OS (*p* = 0.016) and PFS (*p* = 0.02) **(Fig. 6c)**. In contrast, elevated expression of genes encoding enzymes promoting CoA biosynthesis, *PANK3* and *PPAT*, predicted poorer prognosis **(Fig. 6d-e)**. Notably, these trends were not observed in ccRCC, where *PANK4* and *PPAT* expression did not correlate with survival **(Fig. S5d-e)**, and higher *PANK3* expression was unexpectedly associated with better outcomes **(Fig. S5f)**. These findings reveal a fundamental divergence in CoA metabolic programs between RCC subtypes and underscore a unique pathogenic role for MYC-driven activation of CoA biosynthesis specifically in pRCC.

Building on our fly findings showing that CoA biosynthesis activates the MVA-isoprenoid axis to promote cell proliferation **(Fig. 3)**, we next evaluated the prognostic significance of key genes in these downstream pathways **(Fig. 6a)** in pRCC. Consistent with our fly data, high expression of *FDPS* and *GGPS1*, genes encoding essential enzymes in isoprenoid backbone synthesis that generate GPP, FPP and GGPP, correlated with poorer OS (*p* = 0.05 and *p* = 0.015) and PFS (*p* < 0.0001 and *p* = 0.00026) **(Fig. 6f-g)**. These findings support a model in which enhanced isoprenoid production contributes to pRCC progression. Since FPP and GGPP are required for protein prenylation, a process known to promote proliferative signaling [22], we further examined *FNTA* (GGTase-I-alpha), an enzyme mediating this process, and found that its elevated expression was also significantly linked to reduced survival (OS: *p* < 0.0001, PFS: *p* = 0.00015) **(Fig. S5g)**. Additionally, genes involved in the other major branches of isoprenoid metabolism, including dolichol and CoQ biosynthesis (*NUS1*, *PDSS1*, and *SRD5A3*) and cholesterol synthesis (*FDFT1* and *SQLE*), also showed significant associations with worse prognosis **(Fig. S5h-l)**. Collectively, these results implicate multiple isoprenoid-driven processes in pRCC pathogenesis and highlight their potential prognostic and therapeutic relevance.

To further evaluate the prognostic utility of these metabolic pathways, we performed time-dependent Receiver Operating Characteristic (ROC) analyses. Multi-gene signatures for CoA biosynthesis (*PANK1-4*, *PPCS*, *PPCDC, PPAT*, and *COASY*) and isoprenoid backbone biosynthesis (*MVK*, *PMVK*, *MVD*, *IDI1*/*2*, *FDPS*, and *GGPS1*) showed strong predictive power at the 12-month mark in low-metastasis cases (AUC = 0.944 and 0.873, respectively) **(Fig. 6h-i)**, highlighting these pathways as promising early prognostic markers in pRCC. We next stratified by gender and found that the predictive performance of the CoA-biosynthesis and isoprenoid-biosynthesis signatures was more pronounced in male patients **(Fig. S6a-d)**. Intriguingly, gene sets associated with prenylation (*FNTA*, *FNTB*, and *PGGT1B*), dolichol/CoQ biosynthesis (*DHDDS*, *NUS1, PDSS1*/*2*, and *SRD5A3*) and cholesterol synthesis (*FDFT1, SQLE, LSS, CYP51A1,* and *DHCG7*) also displayed high predictive accuracy, particularly in males **(Fig. S6e-m)**, suggesting coordinated regulation along the CoA-isoprenoid axis and potential sex-specific clinical relevance. Collectively, analyses on clinical datasets are largely consistent with our *Drosophila* findings, in which renal CoA biosynthesis promotes intestinal isoprenoid production and activates downstream mechanisms that drive gut tumor progression. Different from the inter-organ metabolic coordination observed in flies, renal CoA production in humans is likely to activate isoprenoid biosynthesis and utilization locally during pRCC progression. Nevertheless, our findings support a conserved role for the CoA-isoprenoid metabolic axis in pRCC and nominate metabolic branches within this axis as candidate targets for therapeutic intervention.

## Discussion

In this study, we uncover a previously unrecognized role for Myc/MYC-driven CoA biosynthesis in orchestrating a systemic inter-organ signaling axis that governs stem cell proliferation and tumor growth. Specifically, (1) we discover that dietary VB5 supplementation is sufficient to induce ISC proliferation, revealing a previously unappreciated link between vitamin intake and tissue homeostasis; (2) we identify fly MT/kidney as a primary site for CoA biosynthesis and demonstrate that renal CoA production non-autonomously promotes MVA and isoprenoid biosynthesis in the gut to stimulate ISC proliferation; (3) we show that inhibition of renal CoA biosynthesis suppresses tumor cell proliferation in a fly gut tumor model, implicating host metabolic reprogramming in supporting tumor growth; (4) we demonstrate that the oncogenic transcription factor Myc directly regulates *Fbl* and *dPANK4* in the kidney, uncovering a tissue-specific transcriptional control of CoA metabolism; (5) we establish the clinical relevance of the MYC-CoA-isoprenoid axis in pRCC, highlighting its potential as both a biomarker and therapeutic target. Collectively, our findings reveal a conserved mechanism that links nutrient availability to inter-organ metabolic coordination, stem cell proliferation, and tumor progression, with broader implications for systemic metabolic regulation and therapeutic intervention in cancer.

### CoA biosynthesis in inter-organ metabolic coordination

Although CoA is synthesized in nearly all tissues, its abundance varies markedly across mammalian organs, with the liver, heart, and kidney exhibiting particularly large CoA pools [7, 8]. These differences suggest that individual tissues maintain distinct CoA reservoirs to support their unique metabolic demands. Beyond their local functions, tissue-specific CoA pools may also contribute to systemic metabolic regulation in response to physiological stimuli. For instance, the hepatic CoA pool plays a central role in orchestrating the metabolic transition from the fed to the fasted state [66]. However, the physiological relevance of CoA pools in other CoA-rich organs, such as the kidney, remains poorly understood. In this study, we show that CoA is preferentially synthesized in the kidney to support anabolic demands in other tissues under specific physiological (e.g., mating) and pathological conditions (e.g., tumorigenesis) in *Drosophila*. Notably, the rate-determining enzyme dPANK4, a negative regulator of CoA biosynthesis, is highly enriched in this tissue, suggesting that renal CoA production is normally restrained. However, under conditions of elevated metabolic demand, such as mating or tumor growth, the CoA biosynthetic machinery in the kidney becomes activated and contributes substantially to the global CoA availability, thereby supporting tissue homeostasis and stem cell/ tumor cell proliferation. CoA itself cannot cross biological membranes [7, 8], historically leading to the assumption that its function is restricted to the cell or tissue where it is synthesized. However, our findings demonstrate that CoA production in the kidney contributes to gut homeostasis in a non-autonomous manner, implying inter-organ communication, perhaps via CoA-related metabolites. To reconcile this, several plausible mechanisms have been proposed in recent years. For example, 4’-phosphopantetheine, a CoA precursor downstream of VB5, can circulate systemically and may be taken up by other tissues to jump-start CoA biosynthesis [67, 68]. Notably, this intermediate is thought to be the specific substrate of PANK4. Another possibility is that metabolites derived from the acyl groups carried by CoA and produced in a CoA-dependent manner in the MT may be released and distributed to other tissues, thus bypassing the need for CoA-dependent production of those carbon groups in the recipient tissue. These transportable metabolites could include free fatty acids, acyl groups transferred to carnitine, smaller carboxylate anions such as acetate and succinate, or the ketone body acetoacetate [69–71]. It is therefore possible that CoA precursor intermediates or metabolites produced in a CoA-dependent manner are transported from the kidney to the gut. In *Drosophila*, the anatomical connection between the MTs and gut enables direct metabolite exchange. In mammals, however, whether renal CoA production contributes to systemic metabolic homeostasis via the circulation or other mechanisms remains an important avenue for future investigation. Nevertheless, our study highlights an unappreciated role for the renal CoA pool in supporting CoA demands in other organs through inter-organ metabolic exchange.

### CoA-fueled isoprenoid metabolism drives proliferative signaling

While CoA metabolism is known to support fundamental processes such as TCA cycle activity and FA biosynthesis [11, 17, 72], its involvement in MVA pathway, a CoA-dependent lipid biosynthetic route, has received comparatively less attention. Our *in vivo* VB5 tracing revealed that renal activation of CoA biosynthesis increases flux to HMG-CoA, the key entry point of the MVA pathway. We further demonstrate that enhanced CoA production in the MTs boosts MVA pathway activity in the gut and promotes the biosynthesis of downstream isoprenoid metabolites, providing previously unreported *in vivo* evidence that CoA availability directly regulates MVA/isoprenoid output. Functionally, activation of this axis drives both stem cell and tumor cell proliferation. Notably, while cholesterol is a well-known MVA-derived metabolite linked to tumor progression in mammals [24, 25], *Drosophila* lacks the cholesterol biosynthesis branch [48], allowing for focused dissection of non-sterol MVA outputs. Despite the absence of this branch, inhibition of the MVA pathway markedly suppresses cell proliferation in both VB5-supplemented and tumor-bearing flies, underscoring the functional importance of alternative downstream branches such as prenylation and CoQ/dolichol synthesis. Supporting this, inhibition of prenylation or CoQ/dolichol synthesis also abolished VB5 supplementation-induced ISC proliferation. Consistently, our analysis of kidney cancer pRCC reveals a pathogenic role for these isoprenoid downstream activities. While CoA biosynthesis has been implicated in supporting tumor growth by fueling the TCA cycle [17], our findings reveal an additional mechanism through which CoA metabolism contributes to tumorigenesis by promoting isoprenoid biosynthesis.

### From historically intractable MYC to targetable CoA biosynthesis

MYC is among the most frequently dysregulated oncogenes in human cancers yet remains challenging to target directly in clinical settings [73, 74]. This limitation has fueled efforts to identify essential downstream pathways that can be therapeutically targeted in MYC-driven tumors. In this study, we identify the CoA biosynthesis pathway as a potential metabolic vulnerability downstream of MYC activity. In *Drosophila*, the chromatin profiling and gene expression analyses indicate that Myc binds the promoter regions of both *Fbl* (orthologous to mammalian *PANK1–3*) and *dPANK4*, two key rate-determining enzymes in CoA biosynthesis. Functionally, Myc activates *Fbl* while repressing *dPANK4*, thereby enhancing CoA biosynthetic flux. This differential regulation at Myc-bound loci likely reflects the fact that MYC acts as both a transcriptional activator and repressor, depending on its interaction with cofactors such as MAX or MIZ1[75]. Consistent with our fly data, MYC binds to promoter regions of all four *PANK* genes in mammals, and its expression inversely correlates with that of *PANK4* in human kidney cancer pRCC. These observations support a conserved role of MYC-dependent tuning of CoA production. Importantly, inhibiting CoA biosynthesis downstream of Myc suppresses tumor growth and extends survival of tumor-bearing flies. In line with these findings, we uncover a pathogenic role for MYC-driven activation of CoA biosynthesis in pRCC. Together, these findings position CoA synthesis as a therapeutically tractable metabolic node in MYC-high cancers.

PDGF/VEGF signaling, classically associated with angiogenesis, is known to regulate MYC [59, 76, 77]. Our prior work demonstrated that PDGF/VEGF signaling is activated in the fly kidney by tumor-secreted Pvf1 [42]. Here, we further show that this PDGF/VEGF-MYC axis amplifies renal CoA biosynthesis, which in turn supports gut tumor growth. This tumor-host circuit highlights a previously unrecognized role for angiogenic signaling in reprogramming host organ metabolism across tissues. Therapeutically, inhibitors targeting the VEGF/PDGF axis are widely used in anti-angiogenic cancer treatment; however, resistance frequently develops, often linked to tumor metabolic adaptation under anaerobic conditions [78, 79]. Our findings suggest that targeting CoA metabolism downstream of PDGF/VEGF signaling may offer an alternative strategy to overcome these therapeutic limitations. We therefore nominate the CoA biosynthetic pathway as a metabolic vulnerability in PDGF/VEGF- and MYC-driven tumors, proposing that inhibition of CoA biosynthesis, alone or in combination with anti-angiogenic regimens, may help restrict metabolic adaptation and impede tumor progression.

### CoA-isoprenoid metabolic axis as pRCC-specific vulnerability

RCC is the most common form of kidney cancer in adults, comprising two major subtypes, ccRCC ad pRCC [63]. Despite distinct molecular and clinicopathologic features between these two subtypes, current diagnostic and therapeutic approaches for pRCC are largely extrapolated from ccRCC-focused studies, resulting in only modest improvements in clinical outcomes [80–82]. Notably, pRCC is structurally and functionally closer to renal tubules, the mammalian counterpart of fly MTs, which allows our fly model to provide relevant mechanistic insight to this tumor subtype. In *Drosophila*, we define a functional CoA-isoprenoid axis in which CoA production enhances MVA pathway activity and boosts the synthesis of downstream isoprenoid metabolites, driving stem cell proliferation and tumor growth. Extending these findings to humans, integrated transcriptomic and clinical analyses in pRCC show that high expression of CoA biosynthetic and isoprenoid pathway genes strongly correlates with poor survival, revealing potential therapeutic vulnerabilities within the CoA-isoprenoid axis. Furthermore, multi-gene signatures for CoA synthesis and terpenoid backbone synthesis display strong early predictive performance, supporting their potential as biomarkers for early-stage patient stratification. Importantly, these associations are specific to pRCC and are not observed in ccRCC. Reconciling our findings, prior cell line studies indicate that the involvement of isoprenoid metabolites in supporting proliferation varies by cell type, as supplementation with individual intermediates can sometimes rescue, but often fail to prevent, the antiproliferative effects of statins [22, 83, 84]. This heterogeneity likely reflects differences in cellular reserves of isoprenoids and the specific proliferative demands for these metabolites across cell types or tissues. Together, our cross-species analysis identifies the CoA-isoprenoid metabolic axis as a pRCC-specific vulnerability, highlighting a promising target for biomarker development and therapeutic intervention in this under-characterized cancer subtype.

### Limitation of this study

We propose an aquaporin-mediated kidney-gut metabolite exchange model in which CoA-derived metabolites are transported from the MTs to the gut via renal-gut countercurrent flow. However, owing to current technical limitations, we were unable to directly identify or quantify the specific metabolites delivered through this pathway, and we cannot exclude the possibility that signaling molecules are also transported. Thus, the molecular identities and transport dynamics of the CoA-derived metabolites remain unresolved. Nevertheless, our genetic and physiological data support the existence of kidney-gut exchange, and future studies will be needed to definitively characterize the metabolites involved and establish their functional contributions to gut homeostasis.

## Methods

All experiments were performed in compliance with all relevant ethical regulations. Analyses of human data were conducted exclusively using publicly available, de-identified datasets (e.g., TCGA), and thus did not require IRB approval. No live vertebrate animals were used in this study. Both males and females were used in this study. Females were used in most experiments as they show more significant phenotypes.

### *Drosophila* husbandry, diet, and strains

Flies were maintained on standard cornmeal-yeast-agar medium at 25 °C under a 12-hour light/dark cycle unless otherwise noted. Fly crosses were grown at 18 °C to inactivate Gal4 and LexA. Adult offspring were collected within 48 hours after eclosion, kept at 18 °C for 24-48 hours, and then shifted to 29 °C for the indicated duration to induce transgene expression (e.g., “day 3” indicates 3 days post induction). Unless stated otherwise, twelve female flies and three male flies were put in one vial and were flipped to fresh food every other day unless noted. To compare virgin and mated female flies, samples were collected 72 hours after mating. For vitamin supplementation experiments, B vitamins were individually added to cooled standard fly food. The amount of vitamins was determined based on those used in *Drosophila* holidic medium protocols (see **Supplementary Table 1**) [35–37]. For VB5 supplementation experiments, 2.5 mM VB5 was added to the cooled fly food unless otherwise noted. For VB5 depletion experiments, flies were fed the *Drosophila* holidic medium specifically lacking VB5 [35–37]. For simvastatin treatment experiments, 0.25 mM or 2.5 mM simvastatin were added to cooled standard fly food. For 4-NB treatment experiments, 5 mM or 25 mM 4-NB were added to cooled standard fly food. For both experiments, an equal volume of DMSO was added to the food as a vehicle control. Flies were fed on special diet for 72 hours before sample collection unless otherwise noted. Fly stocks used include: *esg-Gal4 (P[GawB]NP5130), tub-Gal80ts, tub-Gal80ts; Su(H)GBE-Gal4* (Perrimon lab stock), *Mex-Gal4/FM7;tub-Gal80ts* (BDSC 91367; combined with *tub-Gal80ts* by Perrimon lab), *UAS-GFP* [31], *esg-LexA::GAD* (BDSC 66632), *tub-Gal80ts, CG31272-Gal4* (MT driver) [42], *CG31272-Gal4* (BDSC 76171), *UAS-Yki3SA* [85], *LexAop-Yki3SA-GFP* [86], *UAS-CG5828-RNAi* (VDRC 27523, BDSC 57227, NIG 5828R-3), *UAS-CG5828* (this study), *UAS-Fbl-RNAi* (BDSC 64596, BDSC 35259), *UAS-Hmgcr-RNAi* (BDSC 50652), *UAS-Qm-RNAi* (BDSC 65179), *UAS*-*β-GGT-I-RNAi* (BDSC 34687), *UAS-qless-RNAi* (BDSC 60026), *UAS-Pprd-RNAi* (BDSC 32379), *UAS-Mof-RNAi* (BDSC 58281), *UAS-Hat1-RNAi* (BDSC 42488), *UAS-Gcn5-RNAi* (BDSC 35601), *UAS-Smvt-RNAi* (VDRC 40650), *UAS-Myc-RNAi* (BDSC 43962, BDSC 25783), *UAS-Myc-HA* (BDSC 64759), *UAS-Drip-RNAi* (BDSC 44661), *UAS-Prip-RNAi* (BDSC 50695), *UAS-Pvf1* (BDSC 58426), *UAS-Pvr-act* (BDSC 58496), *UAS-Pvr-RNAi* (VDRC 43459). *w1118* was used as a control. All experiments used female flies unless otherwise specified. Genotypes are listed in **Supplementary Table 3**.

### Molecular cloning and generation of transgenic flies

The UAS-CG5828 (dPANK4) was generated by insertion of the CG5828 into pENTR^TM^ TOPO vector, and then into a fly pWALIUM10-roe vector through Gateway recombination. Primers used are listed in **Supplementary Table 4**. The transgenic flies were generated by injecting embryos with the specified landing site (attP2 and attP40) using PhiC31-mediated site-specific recombination by UniHuaii Corporation.

### Immunofluorescence and imaging

Adult guts were dissected in cold 1× PBS and fixed for 30 min in 4% formaldehyde. Tissues were washed in 0.1% 1 × PBS with 0.1% Triton X-100 (PBST) and mounted in Vectashield with DAPI (Vector Laboratories, H-1200). For pH3 staining, samples were blocked with 5% NDS in PBST, incubated with rabbit anti-pH3 (1:500, CST 9701L) overnight, followed by donkey anti-rabbit Alexa Fluor 594 (1:1,000, Invitrogen A12381) for 1 hour. After three washes in PBST, guts were mounted in Vectashield with DAPI (Vector Laboratories, H-1200). For BODIPY lipid staining, guts were dissected and fixed under the same condition mentioned above. After three times wash in 1 × PBS, the samples were incubated in BODIPY (493/503) reagent (Invitrogen D3922, 1 mg/ml in 1 × PBS) for 30 minutes. The guts were then washed three times in PBS and were mounted in Vectashield with DAPI. Confocal images were acquired on an Olympus IX83 system with FV3000 scan unit (FV31S-SW) using a UPLSAPO 20×/0.75 NA, dry objective. Acquisition parameters were kept constant across genotypes within each experiment. Images were collected as z-stacks with z-step (voxel depth = 2.50 µm); in-plane sampling was 0.795 × 0.795 µm/pixel (as recorded in Fiji/ImageJ). The excitation/detection windows were: DAPI: 405 nm excitation, detection 430–470 nm; GFP and BODIPY: 488 nm excitation, detection 500–547 nm; pH3 (Alexa 594): 561 nm excitation; detection 570–670 nm. All matched genotypes were imaged in the same session with identical laser power, detector gain/offset, pinhole, scan speed, and pixel size. Unless noted, guts were imaged in the R4-R5 region. For display and all quantifications, we used maximum-intensity projections spanning the epithelium. Scale bars were rendered from embedded metadata in Fiji. For each image, ROIs were placed using anatomical landmarks blinded to genotype/condition and applied identically across channels. Within the saved R4-R5 ROI, gut width was measured once per gut at a random region perpendicular to the gut axis using the Straight Line tool followed by Analyze➔Measure. pH3^+^ cells of the whole gut were counted manually; counting was performed blinded. Percentage of GFP^+^ cells was calculated as (number of GFP^+^ cells/ total cell) x 100, where total cells were defined by nuclear (DAPI) counts within an identical ROI using Fiji-imageJ. Adult fly bloating phenotype were captured using a ZEISS Axiozoom V16 fluorescence microscope. Bloating was quantified by the abdomen-to-head area ratio using Fiji-imageJ.

### Metabolite extraction, profiling, and VB5 isotope tracing in flies

Metabolites were extracted in 80% methanol. 12 adult flies or 30 dissected guts were homogenized in 1.5 ml Eppendorf tubes using 400 µl of ice-cold 80% HPLC-grade methanol and 0.5 mm zirconia beads. Following homogenization, samples were centrifuged at 20,000 × g for 5 min at 4 °C. The supernatant was transferred to a new 1.5 ml Eppendorf tube. For maximal extraction efficiency, the pellet was re-extracted with an additional 400 µl of 80% methanol using the same homogenization and centrifugation procedure, and the resulting supernatant was combined with the first. The combined extracts were vacuum dried and stored at -80 °C until analysis. Prior to analysis, dried extracts were resuspended in 20 µl of HPLC-grade water and analyzed using a hybrid 6500 QTRAP triple quadrupole mass spectrometer (AB/SCIEX) coupled to a Prominence UFLC HPLC system (Shimadzu). Selected reaction monitoring (SRM) was used to quantify approximately 300 endogenous water-soluble metabolites. A subset of metabolites was detected in both positive and negative ion modes, yielding 311 SRM transitions through polarity switching. The HPLC gradient began at 85% buffer B (HPLC-grade acetonitrile) and ramped as follows: 85% to 42% B (0–5 min), 42% to 0% B (5–16 min), held at 0% B (16–24 min), returned to 85% B (24–25 min), and re-equilibrated at 85% B for 7 min. Buffer A consisted of 20 mM ammonium hydroxide and 20 mM ammonium acetate (pH 9.0) in 95:5 water:acetonitrile. Peak areas were integrated using MultiQuant v3.2 (AB/SCIEX), and data were analyzed using MetaboAnalyst 6.0 (https://www.metaboanalyst.ca). For VB5 isotope tracing, flies were fed on holidic medium for *Drosophila* [35–37] supplemented with 5 mM ^13^C^15^N-labeled pantothenic acid for 48 h. 12 adult flies were collected in 1.5 ml Eppendorf tubes and the extraction was performed following the steps above. For heavy-isotope-labelled metabolite tracing, SRMs were adapted from previous studies as detailed in our previous publication [11]. For fractional enrichment calculation, isotopologue abundances for each metabolite were first normalized to the total metabolite pool (labeled + unlabeled) and then normalized to labeled VB5 within each sample to account for variability in tracer intake. The resulting values were expressed as fold change relative to the matched control group.

### RT-qPCR of fly samples

The Nucleospin RNA kit (Macherey-Nagel) and TRIzol reagent (Thermo Fisher) was used to extract RNA from MTs and guts, respectively. cDNA was synthesized using the iScript cDNA Synthesis Kit (Bio-Rad, 1708890) and qPCR was performed with a Bio-Rad CFX96 with iQ SYBR Green Supermix (Bio-Rad). Expression was normalized to RP49 or CG13220. Primers are listed in **Supplementary Table 4**.

### Chromatin Immunoprecipitation (ChIP-qPCR)

ChIP was performed using the SimpleChIP Enzymatic Kit (CST 9005). Dissected MTs from adult flies with HA-tagged Myc MT-specific expression (n = 100 for each replicate) were cross-linked in 1.5% formaldehyde for 20 min at room temperature. Cross-linking was stopped by the addition of glycine solution for 5 min at room temperature. Samples were then washed twice with 1 ml 1 × PBS containing 1× Protease Inhibitor Cocktail and disaggregated using a Dounce homogenizer. Nuclei were prepared according to the manufacturer’s protocol and were lysed using Diagenode Bioruptor sonicator to release the cross-linked chromatin. Chromatin was diluted in 1 × ChIP buffer and incubated with 10 µl HA-tagged rabbit monoclonal antibodies (CST 3724) or normal rabbit IgG (CST 2729) overnight at 4 °C with rotation. 30 µl of ChIP-Grade Protein G Magnetic Beads (CST 9006) was incubated with each immunoprecipitation for 2 hours at 4 °C with rotation. Beads were washed and incubated in 150 µl 1 × ChIP Elution Buffer at 65 °C for 30 min with vertexing to elute the chromatin. Cross-links were reversed by adding 6 µl 5 M NaCl and 2 µl Proteinase K to the eluted chromatin supernatant and incubating 2 h at 65 °C, followed by DNA purification step. 1 µl DNA sample was used as template for qPCR to detect enrichments of certain DNA regions. qPCR of a fragment in the *Tll* gene region was used as the negative control. Primer sequences are detailed in the Supplementary Data.

### Survival curve of flies

Survival of flies was analyzed by calculating the percentage of flies alive in each vial incubated at 29 °C. Survival was scored daily starting on day 0 at the same time each day.

### Recombinant GST-CG5828 protein purification

The full-length *Drosophila CG5828* DNA coding sequence (1086 base pairs) was PCR amplified from plasmid DmCD00771312 (DNASU Plasmid Repository) and inserted into a pGEX-4T-2 bacterial expression vector using seamless InFusion cloning (In-Fusion Snap Assembly Master Mix, Takara, 638948), creating N-terminally GST-tagged CG5828.

To create catalytically dead CG5828, aspartate residues predicted to be required for metal coordination and phosphatase activity were mutated to alanine using the Q5 site-directed mutagenesis kit (New England Biolabs, E0554S) and PCR primers that anneal back-to-back including one mutagenic primer and one non-mutagenic primer. All primers were synthesized by Integrated DNA Technologies and resuspended in IDTE buffer [10mM Tris, 0.1mM EDTA]. This construct was expressed in BL21 bacteria (New England Biolabs C2530H) and grown with ampicillin selection at 37 °C. GST-CG5828 protein expression was induced with 250 µM isopropyl β-D-1-thiogalactopyranoside (IPTG; MedChemExpress, HY-15921) for 3 hrs. Bacteria were pelleted by centrifugation, washed and resuspended with PBS at 4 °C, pelleted again and snap frozen in liquid nitrogen. Frozen aliquots were thawed on ice, resuspended in ice-cold resuspension buffer [20 mM Tris, pH 7.5, 140 mM NaCl, 1 mM EDTA, 1mM DTT, 1:100 protease inhibitor cocktail (Millipore-Sigma, P8340)], mixed with 200 µg/ml lysozyme (Thermo Scientific, 89833) for 30 min at 4 °C, then mixed with 0.15% sodium deoxycholate for 15 min at room temp. and finally mixed with 10mM MgCl_2_ and 30 U/mL DNase (New England Biolabs, M0303) for 15 min at room temp. These bacterial lysates were cleared by centrifugation at 15,000 g prior to incubation. with glutathione-conjugated magnetic agarose beads (ThermoFisher Scientific/Pierce, 78602) for 2-3 hrs at 4 °C and elution with 20 mM reduced glutathione (ThermoFisher Scientific, 78259) in 100mM Tris, pH 7.5. Relative protein abundance in GST purifications from bacterial lysate was assessed using Coomassie R-250 dye staining of SDS-PAGE gels. Elutates were snap frozen in liquid nitrogen and stored as one use aliquots at -80°C.

### Phosphatase activity assays

Based on previous work [10, 11] with recombinant PANK4, cobalt was anticipated to be a preferred cation co-factor for CG5828 and was therefore used for initial assessment of enzymatic activity. For initial comparison of vector, WT, and D-A mutants of CG5828, reaction buffers at a final volume of 50 µL included 50 mM Tris-HCl pH7.5, 0.5 mM Co^2+^, equal amounts of WT and mutant enzyme protein, and either 50 mM para-nitrophenyl phosphate (PNPP; New England Biolabs P0757S) or 0.5 mM 4’-phosphopantetheine (synthesized as previously described [11]) as substrates. Reactions in PCR tubes were initiated by addition of substrate then incubated for 30 min at 30 °C and 350rpm in a ThermoMixer (Eppendorf, 05412503P). For reactions using PNPP as a substrate, completed 50 µL reactions were mixed with 100 µL of 1N NaOH, transferred to flat clear bottom 96-well plates, and para-nitrophenol release was quantified by measuring absorbance at 405nm. For reactions using 4’-phosphopantetheine as the substrate, inorganic phosphate release was quantified using a malachite green phosphate detection kit (R&D Systems, DY996) and measurement of absorbance at 620nm. Absorbance for both assays was measured using an Agilent Biotek plate reader. For phosphatase assays testing multiple metal cations, reaction conditions were as described above, 50mM PNPP was used as the substrate, and indicated metal cations were used at 0.5mM. Metal cation chloride salts were purchased from Sigma-Aldrich: magnesium chloride hexahydrate (M2670-500G), potassium chloride (P9541-1KG), cobalt(II) chloride hexahydrate (202185-100G), nickel(II) chloride hexahydrate (654507-5G), sodium chloride (S7653-250G), calcium chloride hexahydrate (21108-500G), lithium chloride (62476-100G-F), iron(II) chloride tetrahydrate (380024-5G), copper(II) chloride dihydrate (C3279-100G), zinc chloride (Z0152-50G) and from Fischer Scientific, manganese chloride (M87-100).

### Statistics and Reproducibility

Sample sizes were determined through preliminary experiments and previous studies to achieve the required statistical power. Experiments were repeated at least twice and/or use different methods (e.g., different primers, RNAi lines, different time points). Minimum three biological replicates were used in each experiment. The experiments were not randomized. Blinding in sample processing/collection is impossible because animal with different genotypes showed visible distinct phenotypes. Samples were quantified blinded with genotype/condition masked during image quantification and outcome assessment where feasible. GraphPad Prism was used for all statistical analyses. The ROUT outlier test (Q = 10%) was applied to each experimental group before data normalization and removed flagged outliers. Statistical analyses were then conducted on the retained replicates. *p*-values were calculated via unpaired two-tailed *t*-test or one-way ANOVA unless stated. *p*-values < 0.05 were statistically significant. Gene expression was normalized to controls. *n* represents biological replicates. Error bars indicate s.d. Quantitative and statistical parameters, including statistical methods, error bars, n numbers, and *p*-values, are indicated in each figure. Kaplan-Meier (KM) survival curves were generated using the R packages survival and survminer. Time-dependent ROC analysis used the timeROC package, stratified by time point, metastasis status, and gender. hdWGCNA (0.4.05) was used to predict regulon activity using snRNA-sequencing datasets. Briefly, each dataset was subset to the clusters labelled as Malpighian tubule and ran through the hdWGCNA transcription factory regulatory network analysis pipeline using default settings (https://smorabit.github.io/hdWGCNA/articles/tf_network.html). The FindDifferentialRegulons() function was used to perform differential expression analysis on regulon scores/activity. Biorender was used for illustrations. ChatGPT assisted in proofreading; all content was verified and revised by the authors.

## Data Availability

The data generated in this study are provided in the Source Data file. Cell-type-specific expression data were sourced from GSE229526 (https://www.ncbi.nlm.nih.gov/geo/query/acc.cgi?acc=GSE229526), GSE202575 (https://www.ncbi.nlm.nih.gov/geo/query/acc.cgi?acc=GSE202575), GSE251715 (https://www.ncbi.nlm.nih.gov/geo/query/acc.cgi?acc=GSE251715). ChIP-seq against Myc in *D. melanogaster* whole organisms was sourced from GSE49774 (https://www.ncbi.nlm.nih.gov/geo/query/acc.cgi?acc=GSE49774). ChIP-seq against Myc in Mus musculus was sourced from GSE51011 (https://www.ncbi.nlm.nih.gov/geo/query/acc.cgi?acc=GSE51011). Patient transcriptome and survival data were retrieved from TCGA PanCancer Atlas (KIRP and KIRC) via cBioPortal (KIRP: https://master.cbioportal.org/study/summary?id=kirp_tcga_pan_can_atlas_2018; KIRC: https://www.cbioportal.org/study/summary?id=kirc_tcga_pan_can_atlas_2018).

## Acknowledgements

We thank R. Binari, J. Zirin, L. Liu, C. Villalta, S. Mohr, K. Huang, A. Carte, B. Xia, A.R.Kim and all members of the Perrimon lab for their critical suggestions and help on this research. We thank P. M. Llopis, P. V. Anekal, A. Wells, and Microscopy Resources on the North Quad (MicRoN) core facility at Harvard Medical School for advice and help with confocal imaging. We thank the *Drosophila* RNAi Screening Center (DRSC), Bloomington Drosophila Stock Center (BDSC), Vienna *Drosophila* Resource Center (VDRC), and Fly Stocks of National Institute of Genetics (NIG) for providing fly stocks used in this study. We thank FlyBase for the genetics knowledgebase and database resource. This work is funded in part by NIH/NIDDK (R56DK140239 and R01DK140239) (C.C.D) and by the CANCAN team supported by the Cancer Grand Challenges partnership funded by Cancer Research UK (CGCATF-2021/100022) and the National Cancer Institute (1 OT2 CA278685-01). T. M. is supported by the Cystinosis Research Foundation and the Glenn Foundation for Medical Research Postdoctoral Fellowship in Aging Research. Y.L. is supported by the Charles A. King Trust Fellowship. N.P. is an investigator at the Howard Hughes Medical Institute. This article is subject to HHMI’s Open Access to Publications policy. HHMI lab heads have previously granted a non-exclusive CC BY 4.0 license to the public and a sublicensable license to HHMI in their research articles. Pursuant to those licenses, the author-accepted manuscript of this article can be made freely available under a CC BY 4.0 license immediately upon publication.

## Author contributions

T.M., C.D., and N.P. conceived the study and designed the experiments. T.M. conducted the majority of the experiments and the data analysis. T.M., C.D., J.A., and Y.L. designed and conducted the metabolomics studies. T.M., and X.S conducted the molecular cloning experiments for transgenic fly lines. D. A. conducted the phosphatase activity assays. L. d. C. P-A. assisted with the phosphatase assays and Myc experiments. Y.L., M.Q., and Y.H. were responsible for the bioinformatics analysis. T.M., Y.L., Y.H., C.D., and N.P. interpreted the results. T.M. and N.P. wrote the manuscript with contributions and feedback from all authors. All authors reviewed and approved the final manuscript.

