## Supplementary Information for "Renal Coenzyme A (CoA) Production from VB5 Fuels Stem Cell Proliferation and Tumor Growth"

Figs. S1 to S6

Supplementary Table 1-4

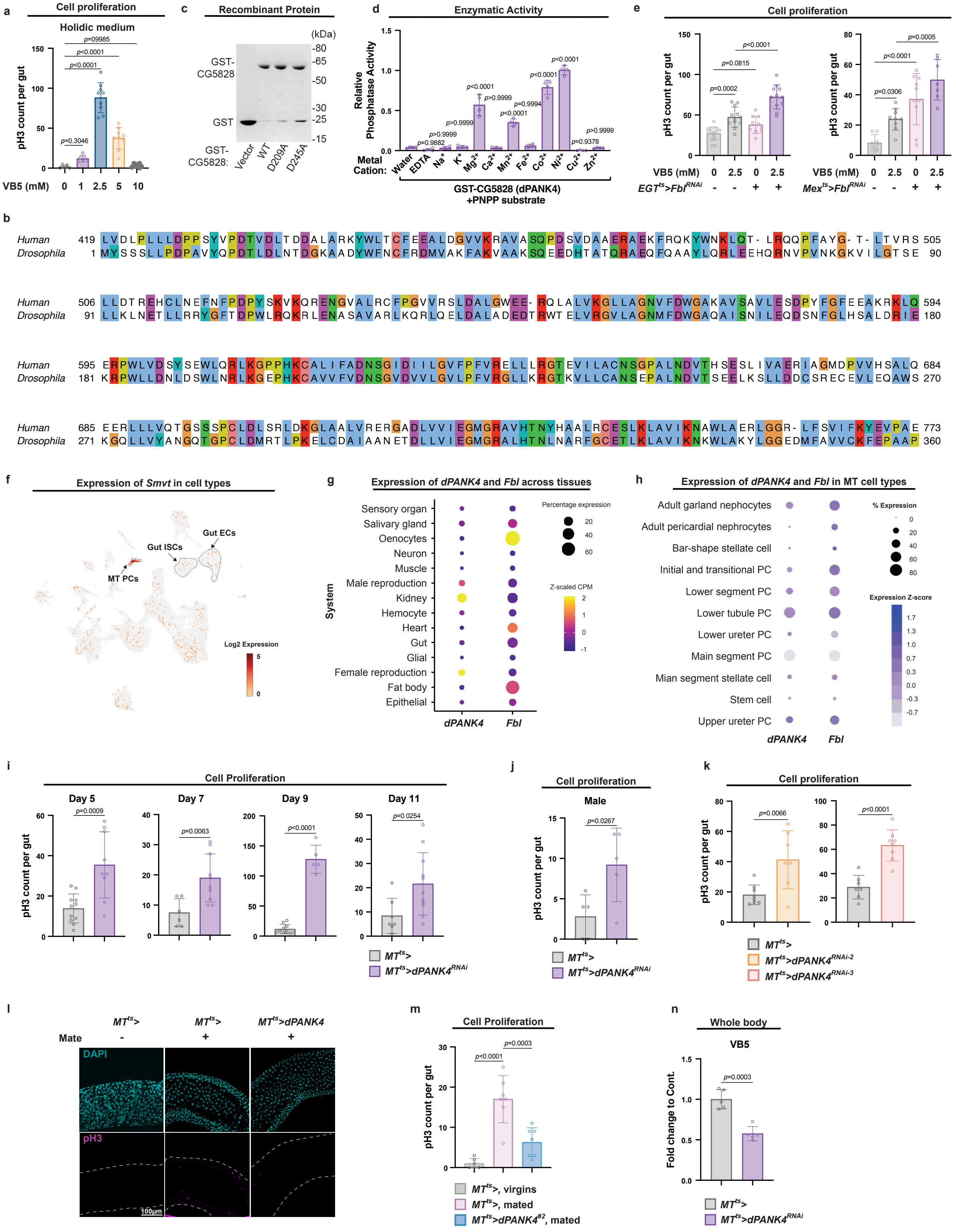

a
Cell proliferation
Holidic medium
pH3 count per gut
VB5 (mM) 0 1 2.5 5 10
c
Recombinant Protein
(kDa)
GST-CG5828
GST
GST-CG5828: Vector WT D209A D245A
d
Enzymatic Activity
Relative Phosphatase Activity
Metal Cation: Water EDTA Na+ K+ Mg2+ Ca2+ Mn2+ Fe2+ Co2+ Ni2+ Cu2+ Zn2+
GST-CG5828 (dPANK4) +PNPP substrate
e
Cell proliferation
pH3 count per gut
VB5 (mM) 0 2.5 0 2.5
EGTts>FblRNAi - - + +
Mexts>FblRNAi - - + +
b
Human
Drosophila
f
Expression of Smvt in cell types
Gut ECs
Gut ISCs
MT PCs
Log2 Expression
g
Expression of dPANK4 and Fbl across tissues
Sensory organ
Salivary gland
Oenocytes
Neuron
Muscle
Male reproduction
Kidney
Hemocyte
Heart
Gut
Glial
Female reproduction
Fat body
Epithelial
System
dPANK4 Fbl
Percentage expression
Z-scaled CPM
h
Expression of dPANK4 and Fbl in MT cell types
Adult garland nephrocytes
Adult pericardial nephrocytes
Bar-shape stellate cell
Initial and transitional PC
Lower segment PC
Lower tubule PC
Lower ureter PC
Main segment PC
Mian segment stellate cell
Stem cell
Upper ureter PC
dPANK4 Fbl
% Expression
Expression Z-score
i
Cell Proliferation
Day 5
Day 7
Day 9
Day 11
pH3 count per gut
MTts>
MTts>dPANK4RNAi
j
Cell proliferation
Male
pH3 count per gut
MTts>
MTts>dPANK4RNAi
k
Cell proliferation
pH3 count per gut
MTts>
MTts>dPANK4RNAi-2
MTts>dPANK4RNAi-3
l
MTts> MTts> MTts>dPANK4
Mate - + +
DAPI
pH3
100μm
m
Cell Proliferation
pH3 count per gut
MTts>, virgins
MTts>, mated
MTts>dPANK4#2, mated
n
Whole body
VB5
Fold change to Cont.
MTts>
MTts>dPANK4RNAi

**Fig. S1 | Related to Figure 1 and 2.** **a,** Quantification of pH3$^+$ cells per midgut from flies with VB5 dietary supplementation at 0, 1, 2.5, 5, or 10 mM on holidic medium. n = 7, 6, 9, 12, 9, 9, and 10 in 0, 1, 2.5, 5, and 10 mM, respectively. **b,** Full length amino acid alignment between human PANK4 (NCBI Reference Sequence: NP_060686.3) and *Drosophila* CG5828 (dPANK4; NCBI Reference Sequence: NP_608907.1). **c,** Coomassie stained SDS-PAGE of vector and GST-CG5828 purification showing relative quantities used in reactions shown in **Fig. 2c**. **d,** Metal cation dependence of CG5828/dPANK4 phosphatase activity. Metal cation preference was determined as in **Fig. 2c** with PNPP substrate and chloride salts of indicated metal cations (0.5mM). For statistical analysis, the relative activities of the GST-CG5828 in chloride salts or EDTA are compared with that in water. n = 4. **e,** Quantification of pH3$^+$ cells per midgut in control versus ISC/EB-specific (*EGT*$^{ts}$>; left) or EC-specific (*Mex*$^{ts}$>; right) *Fbl* knockdown flies under VB5 supplementation. n = 14, 12, 10, and 11 in *EGT*$^{ts}$> (0 mM), *EGT*$^{ts}$> (2.5 mM), *EGT*$^{ts}$> *Fbl-RNAi* (0 mM), and *EGT*$^{ts}$> *Fbl-RNAi* (2.5 mM), respectively. n = 8, 9, 10, and 7 in *Mex*$^{ts}$> (0 mM), *Mex*$^{ts}$> (2.5 mM), *Mex*$^{ts}$> *Fbl-RNAi* (0 mM), and *Mex*$^{ts}$> *Fbl-RNAi* (2.5 mM), respectively. **f.** UMAP visualization of expression levels of *Smvt* across all cell clusters; data retrieved from published snRNA-seq. Malpighian tubule (MT) principal cells (PCs), gut intestinal stem cells (ISCs), and gut enterocytes (ECs) are outlined with dashed lines. **g,** Expression patterns of *dPANK4* and *Fbl* across tissues based on percentage of expressing cells and Z-scaled counts per million (CPM); data from FlyAtlas single-nucleus RNA-seq (snRNA-seq) database. **h,** Expression of *dPANK4* and *Fbl* in cell types of the MTs, represented by percent-expressing cells and Z-scored expression; data from published MT snRNA-seq dataset. PC, principal cell. **i,** Quantification of pH3$^+$ cells per midgut at day 5, 7, 9, and 11 in control and MT-specific *dPANK4* knockdown flies. n = 11 and 9 in *MT*$^{ts}$> and *MT*$^{ts}$>*dPANK4-RNAi* at Day 5, respectively. n = 6 and 11 in *MT*$^{ts}$> and *MT*$^{ts}$>*dPANK4-RNAi* at Day 7, respectively. n = 10 and 5 in *MT*$^{ts}$> and *MT*$^{ts}$>*dPANK4-RNAi* at Day 9, respectively. n = 7 and 11 in *MT*$^{ts}$> and *MT*$^{ts}$>*dPANK4-RNAi* at Day 11, respectively. **j,** pH3$^+$ cell counts in midguts from male flies with or without MT-specific *dPANK4* knockdown at day 10. n = 5. **k,** pH3$^+$ cell counts in midguts of flies with or without MT-specific dPANK4 *knockdown* at day 10 using two additional independent RNAi lines. n = 9 and 7 in *MT*$^{ts}$> and *MT*$^{ts}$>*dPANK4-RNAi-#2*, respectively. n = 7 and 8 in *MT*$^{ts}$> and *MT*$^{ts}$>*dPANK4-RNAi-#3*, respectively. **l,** Representative gut images from virgin and mated female flies with or without MT-specific *dPANK4* overexpression at day 7. In the pH3 panel, the guts are outlined with dashed lines. **m,** Quantification of pH3$^+$ cells per midgut from virgin and mated female flies with or without MT-specific *dPANK4* overexpression at day 7 using an independent *dPANK4* overexpression line. n = 6, 7 and 7 in *MT*$^{ts}$> (virgins), *MT*$^{ts}$> (mated), and *MT*$^{ts}$>*dPANK4-#2 MT*$^{ts}$> (mated), respectively. **n,** Relative metabolite levels of VB5 in control and MT-specific *dPANK4* knockdown flies at day 9, retrieved from whole-body metabolomics analysis. n = 5. Statistical significance assessed by one-way ANOVA (**a, d, e, m)** and unpaired two-sided Student's *t*-test (**i, j, k, n**) and. Error bars indicate s.d., with means at the center. Source data are provided as a Source Data file.

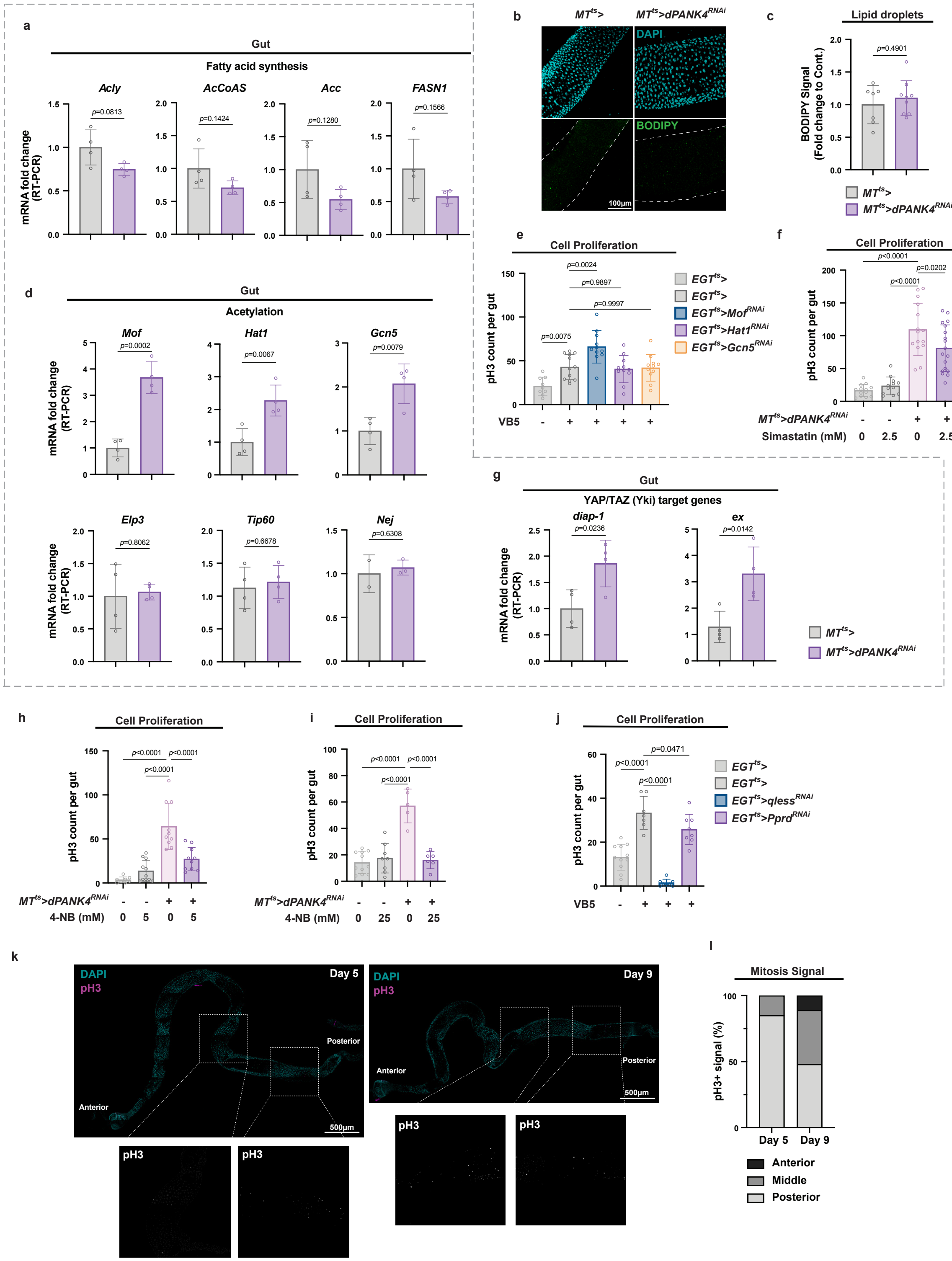
a
Gut
Fatty acid synthesis
Acly
AcCoAS
Acc
FASN1
mRNA fold change (RT-PCR)
p=0.0813
p=0.1424
p=0.1280
p=0.1566
b
MTts>
MTts>dPANK4RNAi
DAPI
BODIPY
100μm
c
Lipid droplets
BODIPY Signal (Fold change to Cont.)
p=0.4901
MTts>
MTts>dPANK4RNAi
d
Gut
Acetylation
Mof
Hat1
Gcn5
p=0.0002
p=0.0067
p=0.0079
Elp3
Tip60
Nej
p=0.8062
p=0.6678
p=0.6308
mRNA fold change (RT-PCR)
e
Cell Proliferation
pH3 count per gut
p=0.0024
p=0.9897
p=0.9997
p=0.0075
VB5
EGTts>
EGTts>
EGTts>MofRNAi
EGTts>Hat1RNAi
EGTts>Gcn5RNAi
f
Cell Proliferation
pH3 count per gut
p<0.0001
p=0.0202
p<0.0001
MTts>dPANK4RNAi
Simastatin (mM)
g
Gut
YAP/TAZ (Yki) target genes
diap-1
ex
p=0.0236
p=0.0142
mRNA fold change (RT-PCR)
MTts>
MTts>dPANK4RNAi
h
Cell Proliferation
pH3 count per gut
p<0.0001
p<0.0001
p<0.0001
MTts>dPANK4RNAi
4-NB (mM)
i
Cell Proliferation
pH3 count per gut
p<0.0001
p<0.0001
p<0.0001
MTts>dPANK4RNAi
4-NB (mM)
j
Cell Proliferation
pH3 count per gut
p=0.0471
p<0.0001
p<0.0001
VB5
EGTts>
EGTts>
EGTts>qlessRNAi
EGTts>PprdRNAi
k
DAPI
pH3
Day 5
Day 9
Anterior
Posterior
500μm
pH3
l
Mitosis Signal
pH3+ signal (%)
Day 5
Day 9
Anterior
Middle
Posterior

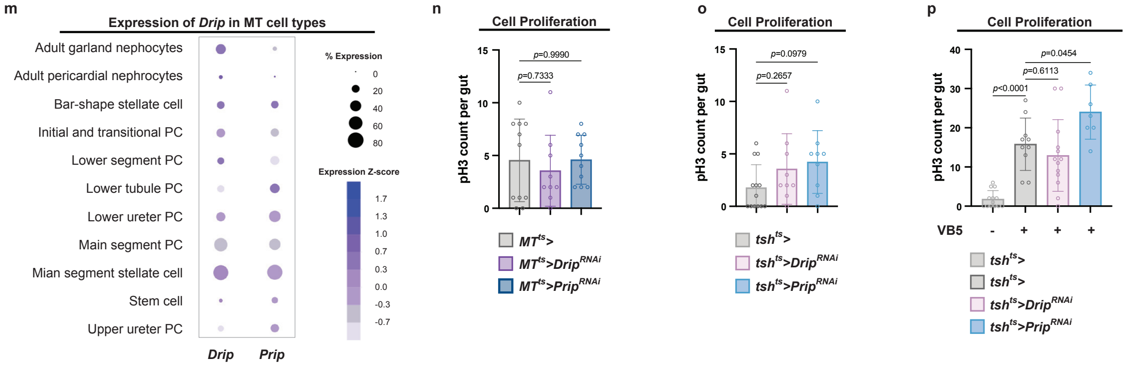
m
Expression of Drip in MT cell types
Adult garland nephocytes
Adult pericardial nephrocytes
Bar-shape stellate cell
Initial and transitional PC
Lower segment PC
Lower tubule PC
Lower ureter PC
Main segment PC
Mian segment stellate cell
Stem cell
Upper ureter PC
Drip
Prip
% Expression
0
20
40
60
80
Expression Z-score
1.7
1.3
1.0
0.7
0.3
0.0
-0.3
-0.7
n
Cell Proliferation
pH3 count per gut
p=0.9990
p=0.7333
MTts>
MTts>DripRNAi
MTts>PripRNAi
o
Cell Proliferation
pH3 count per gut
p=0.0979
p=0.2657
tshts>
tshts>DripRNAi
tshts>PripRNAi
p
Cell Proliferation
pH3 count per gut
p=0.0454
p=0.6113
p<0.0001
VB5
-
+
+
+
tshts>
tshts>
tshts>DripRNAi
tshts>PripRNAi

**Fig. S2 | Related to Figure 3. a,** qRT-PCR analysis of lipogenic genes (*Acly, AcCoAS, Acc,* and *FASN1*) mRNA levels in guts from control and MT-specific *dPANK4* knockdown flies at day 9. n = 4. Acly, ATP citrate lyase, ortholog of ATPCL; AcCoAS, acetyl-CoA synthetase, ortholog of ACSS1/2; ACC, acetyl-CoA carboxylase; FASN1, fatty acid synthase 1. **b, c,** Representative images (**b**) and quantification (**c**) of BODIPY lipid staining in guts from control and MT *dPANK4* knockdown flies at day 9. In the BODIPY panel, the guts are outlined with dashed lines. n = 7 and 9 in $MT^{ts}$> and $MT^{ts}$>*dPANK4-RNAi*, respectively. **d,** qRT-PCR analysis of histone acetyltransferase genes (*Mof, HAT1, Gcn5, Elp3, Tip60,* and *Nej*) mRNA levels in guts from control and MT *dPANK4* knockdown flies at day 9. n = 4. Nej, ortholog of CBP/p300. **e,** Quantification of $pH3^+$ cells per midgut in flies with gut-specific knockdown of *HAT1, Mof,* or *Gcn5* under VB5 supplementation. n = 9, 12, 11, 12, and 11 in $EGT^{ts}$> [VB5 (-)], $EGT^{ts}$> [VB5 (+)], $EGT^{ts}$> *Mof-RNAi* [VB5 (+)], $EGT^{ts}$> *Hat1-RNAi* [VB5 (+)], and $EGT^{ts}$> *Gcn5-RNAi* [VB5 (+)], respectively. **f,** Quantification of $pH3^+$ cells per midgut in flies with or without MT-specific *dPANK4* knockdown at day 10. Flies were fed on DMSO or simvastatin-containing (2.5 mM) diet for 5 days. n = 13, 12, 15, and 18 in $MT^{ts}$> (0 mM), $MT^{ts}$> (2.5 mM), $MT^{ts}$>*dPANK4-RNAi* (0 mM), and $MT^{ts}$>*dPANK4-RNAi* (2.5mM), respectively. **g,** qRT-PCR analysis of YAP/TAZ (Yki) target genes (*diap-1* and *expanded*) mRNA levels in guts from control and MT-specific *dPANK4* knockdown flies at day 9. n = 4. **h-i,** Quantification of $pH3^+$ cells per midgut in flies with or without MT *dPANK4* depletion, fed on diet supplemented with 5 mM (**h**) or 25 mM (**i**) 4-nitrobenzoic acid (4-NB). n = 10, 11, 10, and 10 in $MT^{ts}$> (0 mM), $MT^{ts}$> (5 mM), $MT^{ts}$>*dPANK4-RNAi* (0 mM), and $MT^{ts}$>*dPANK4-RNAi* (5mM) in **h**, respectively. n = 11, 8, 5, and 6 in $MT^{ts}$> (0 mM), $MT^{ts}$> (25 mM), $MT^{ts}$>*dPANK4-RNAi* (0 mM), and $MT^{ts}$>*dPANK4-RNAi* (25mM) in **i**, respectively. **j,** $pH3^+$ cell quantification in control flies or flies with gut-specific knockdown of *qless* or *Pprd* under VB5 supplementation. n = 11, 7, 8, and 8 in $EGT^{ts}$> [VB5 (-)], $EGT^{ts}$> [VB5 (+)], $EGT^{ts}$> *qless-RNAi* [VB5 (+)], and $EGT^{ts}$> *Pprd-RNAi* [VB5 (+)], respectively. **k,** Representative whole midgut images from control and MT *dPANK4* knockdown flies at day 5 and day 9 showing posterior-biased $pH3^+$ signal at day 5 and anterior progression by day 9. **l,** Distribution of mitosis signal in anterior (R1-2), middle (R3), and posterior (R4-5) guts of control and MT *dPANK4* knockdown flies at day 5 and day 9. n = 3. **m,** Expression profile of *Drip* and *Prip* in MT cell types, shown as percent of expressing cells and Z-scored expression; data from published MT snRNA-seq dataset. **n,** Quantification of $pH3^+$ cells per midgut in control flies and flies with *Drip* or *Prip* knockdown in principal cells ($MT^{ts}$>). n = 13, 9, and 10 in $MT^{ts}$>, $MT^{ts}$>*Drip-RNAi*, and $MT^{ts}$>*Prip-RNAi*, respectively. **o,** Quantification of $pH3^+$ cells per midgut in control flies and flies with *Drip* or *Prip* knockdown in stellate cells ($tsh^{ts}$>). n = 13, 9, and 11 in $tsh^{ts}$>, $tsh^{ts}$>*Drip-RNAi*, and $tsh^{ts}$>*Prip-RNAi*, respectively. **p,** Quantification of $pH3^+$ cells per midgut in control flies and flies with *Drip* or *Prip* knockdown in stellate cells ($tsh^{ts}$>) under VB5 supplementation. n = 13, 10, 14 and 7 in $tsh^{ts}$> [VB5 (-)], $tsh^{ts}$> [VB5 (+)], $tsh^{ts}$>*Drip-RNAi* [VB5 (+)], and $tsh^{ts}$>*Prip-RNAi* [VB5 (+)], respectively. Statistical significance assessed by unpaired two-sided Student's *t*-test (**a, c, d, g**) and one-way ANOVA (**e, f, h, l, j, n, o, p)**. Error bars indicate s.d., with means at the center. Source data are provided as a Source Data file.

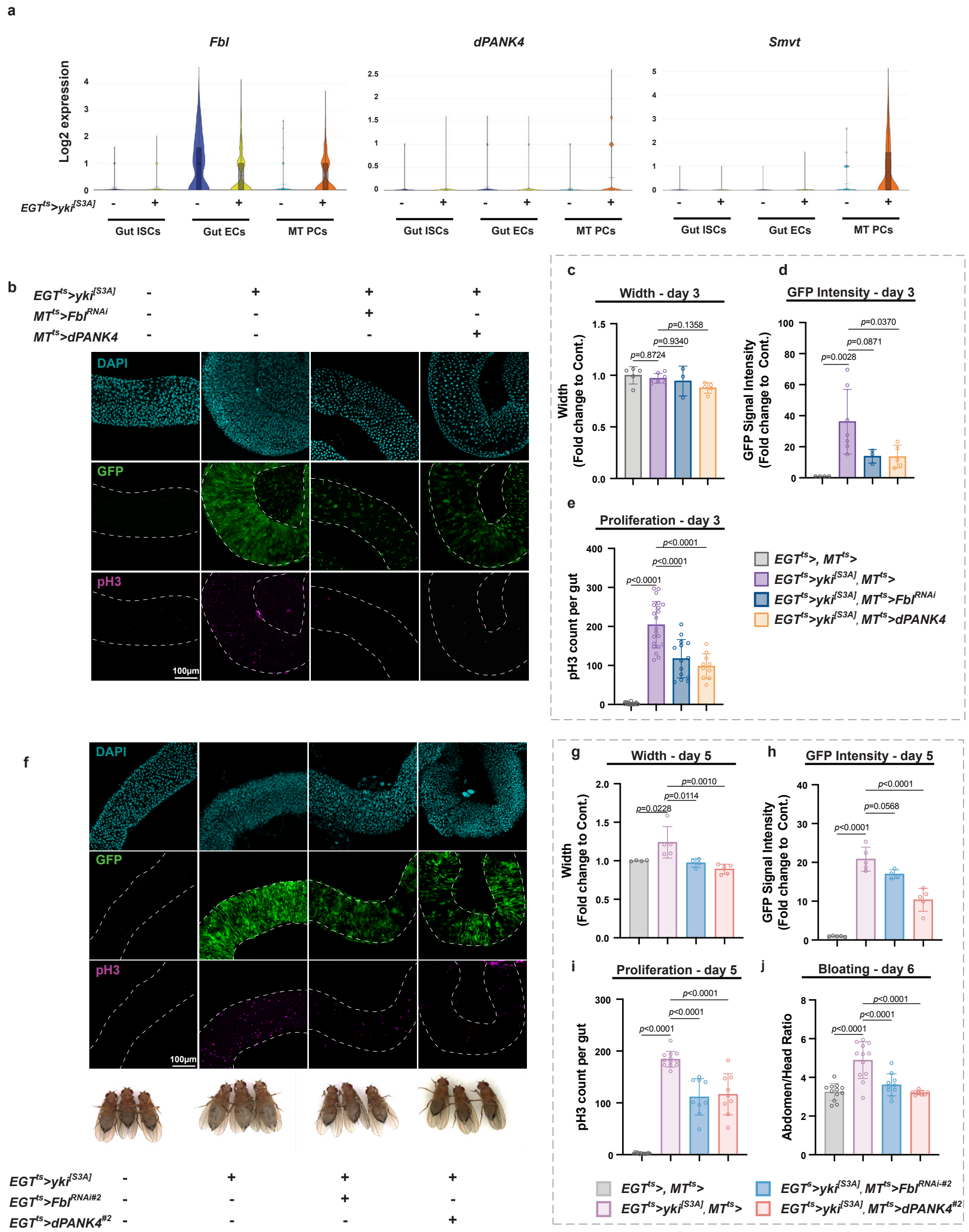
a
Fbl
dPANK4
Smvt
Log2 expression
EGT^ts>yki^[S3A]
Gut ISCs
Gut ECs
MT PCs
b
EGT^ts>yki^[S3A]
MT^ts>Fbl^RNAi
MT^ts>dPANK4
DAPI
GFP
pH3
100μm
c
Width - day 3
Width
(Fold change to Cont.)
p=0.8724
p=0.9340
p=0.1358
d
GFP Intensity - day 3
GFP Signal Intensity
(Fold change to Cont.)
p=0.0028
p=0.0871
p=0.0370
e
Proliferation - day 3
pH3 count per gut
p<0.0001
p<0.0001
p<0.0001
EGT^ts>, MT^ts>
EGT^ts>yki^[S3A], MT^ts>
EGT^ts>yki^[S3A], MT^ts>Fbl^RNAi
EGT^ts>yki^[S3A], MT^ts>dPANK4
f
DAPI
GFP
pH3
100μm
EGT^ts>yki^[S3A]
EGT^ts>Fbl^RNAi#2
EGT^ts>dPANK4^#2
g
Width - day 5
Width
(Fold change to Cont.)
p=0.0228
p=0.0114
p=0.0010
h
GFP Intensity - day 5
GFP Signal Intensity
(Fold change to Cont.)
p<0.0001
p=0.0568
p<0.0001
i
Proliferation - day 5
pH3 count per gut
p<0.0001
p<0.0001
p<0.0001
j
Bloating - day 6
Abdomen/Head Ratio
p<0.0001
p<0.0001
p<0.0001
EGT^ts>, MT^ts>
EGT^ts>yki^[S3A], MT^ts>
EGT^ts>yki^[S3A], MT^ts>Fbl^RNAi-#2
EGT^ts>yki^[S3A], MT^ts>dPANK4^#2

k

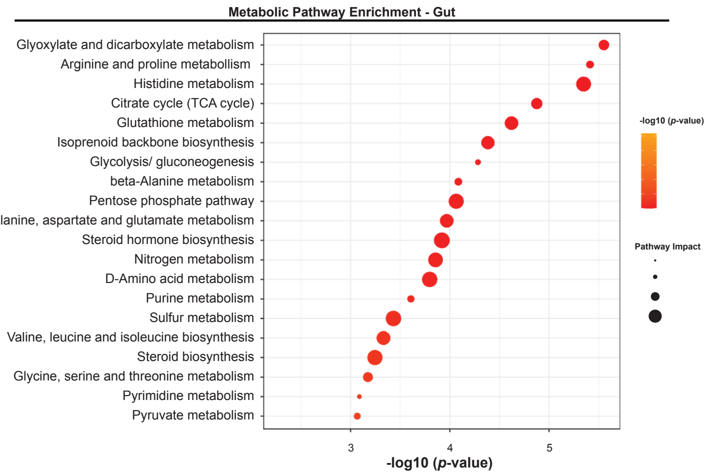
Metabolic Pathway Enrichment - Gut
Glyoxylate and dicarboxylate metabolism
Arginine and proline metabollism
Histidine metabolism
Citrate cycle (TCA cycle)
Glutathione metabolism
Isoprenoid backbone biosynthesis
Glycolysis/ gluconeogenesis
beta-Alanine metabolism
Pentose phosphate pathway
Alanine, aspartate and glutamate metabolism
Steroid hormone biosynthesis
Nitrogen metabolism
D-Amino acid metabolism
Purine metabolism
Sulfur metabolism
Valine, leucine and isoleucine biosynthesis
Steroid biosynthesis
Glycine, serine and threonine metabolism
Pyrimidine metabolism
Pyruvate metabolism
3
4
5
-log10 (p-value)
-log10 (p-value)
Pathway Impact

l

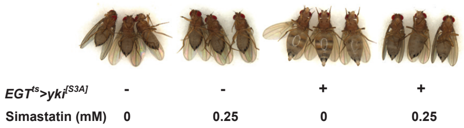
EGTts>yki[S3A]
-
-
+
+
Simastatin (mM)
0
0.25
0
0.25

m

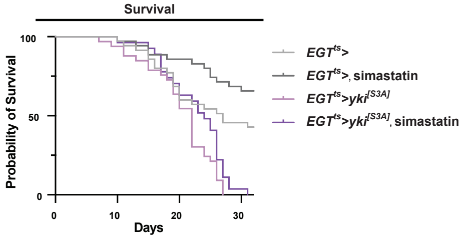
Survival
100
50
0
Probability of Survival
0
10
20
30
Days
EGTts>
EGTts>, simastatin
EGTts>yki[S3A]
EGTts>yki[S3A], simastatin

**Fig. S3 | Related to Figure 4. a,** Log2 expression of *dPANK4*, *Fbl*, and *Smvt* in gut ISCs, gut ECs and MT PCs of control and Yki flies; data from published whole-body Yki snRNA-seq dataset. **b,** Representative images of guts from control flies and Yki flies with or without MT-specific *Fbl* knockdown or *dPANK4* overexpression at day 3. In the GFP and pH3 panels, the guts are outlined with dashed lines. **c–e,** Quantification of gut width (**c**), GFP intensity (**d**), and pH3$^+$ cell counts (**e**). n = 5, 7, 3, and 5 in $EGT^{ts}$>, $MT^{ts}$>, $EGT^{ts}$>$yki^{[S3A]}$, $MT^{ts}$>, $EGT^{ts}$>$yki^{[S3A]}$, $MT^{ts}$>*Fbl-RNAi*, and $EGT^{ts}$>$yki^{[S3A]}$, $MT^{ts}$>*dPANK4* in **c**, respectively.  n = 5, 7, 3, and 5 in $EGT^{ts}$>, $MT^{ts}$>, $EGT^{ts}$>$yki^{[S3A]}$, $MT^{ts}$>, $EGT^{ts}$>$yki^{[S3A]}$, $MT^{ts}$>*Fbl-RNAi*, and $EGT^{ts}$>$yki^{[S3A]}$, $MT^{ts}$>*dPANK4* in **d**, respectively. n = 24, 21, 14, and 10 in $EGT^{ts}$>, $MT^{ts}$>, $EGT^{ts}$>$yki^{[S3A]}$, $MT^{ts}$>, $EGT^{ts}$>$yki^{[S3A]}$, $MT^{ts}$>*Fbl-RNAi*, and $EGT^{ts}$>$yki^{[S3A]}$, $MT^{ts}$>*dPANK4* in **e**, respectively. **f,** Representative images of guts (day 5) and representative images of bloating phenotypes (day 6) from control flies and Yki flies with or without MT-specific *Fbl* knockdown or *dPANK4* overexpression using independent fly lines. **g-i,** Quantification of gut width (**g**), GFP intensity (**h**), and pH3$^+$ cell counts (**i**). n = 4, 5, 4, and 5 in $EGT^{ts}$>, $MT^{ts}$>, $EGT^{ts}$>$yki^{[S3A]}$, $MT^{ts}$>, $EGT^{ts}$>$yki^{[S3A]}$, $MT^{ts}$>*Fbl-RNAi-#2*, and $EGT^{ts}$>$yki^{[S3A]}$, $MT^{ts}$>*dPANK4-#2* in **g**, respectively. n = 5, 5, 4, and 5 in $EGT^{ts}$>, $MT^{ts}$>, $EGT^{ts}$>$yki^{[S3A]}$, $MT^{ts}$>, $EGT^{ts}$>$yki^{[S3A]}$, $MT^{ts}$>*Fbl-RNAi-#2*, and $EGT^{ts}$>$yki^{[S3A]}$, $MT^{ts}$>*dPANK4-#2* in **h**, respectively. n = 15, 14, 9, and 9 in $EGT^{ts}$>, $MT^{ts}$>, $EGT^{ts}$>$yki^{[S3A]}$, $MT^{ts}$>, $EGT^{ts}$>$yki^{[S3A]}$, $MT^{ts}$>*Fbl-RNAi-#2*, and $EGT^{ts}$>$yki^{[S3A]}$, $MT^{ts}$>*dPANK4-#2* in **i**, respectively. **j,** Bloating quantification via abdomen-to-head ratio of panel **f**. n = 12, 12, 10, and 12 in $EGT^{ts}$>, $MT^{ts}$>, $EGT^{ts}$>$yki^{[S3A]}$, $MT^{ts}$>, $EGT^{ts}$>$yki^{[S3A]}$, $MT^{ts}$>*Fbl-RNAi-#2*, and $EGT^{ts}$>$yki^{[S3A]}$, $MT^{ts}$>*dPANK4-#2*, respectively. **k,** Pathway enrichment analysis of gut metabolomics profiling of flies with Yki flies versus controls at day 6. The top enriched KEGG pathways are shown, ranked by statistical significance (-log10(*p*-value)) and colored accordingly. Dot size represents pathway value. n = 4. **l,** Representative images of bloating phenotypes in control and Yki flies with or without simvastatin treatment (0.25 mM) at day 6. Flies were fed on DMSO or simvastatin-containing diet for 6 days. **m**, Survival curve of control and Yki flies with or without simvastatin treatment (0.25 mM). n = 27-35. Statistical significance assessed by one-way ANOVA (**c-e, g-j**), two-tailed Fisher's exact test (**k**), and log-rank (Mantel–Cox) test (**m**). Error bars indicate s.d., with means at the center. Source data are provided as a Source Data file.

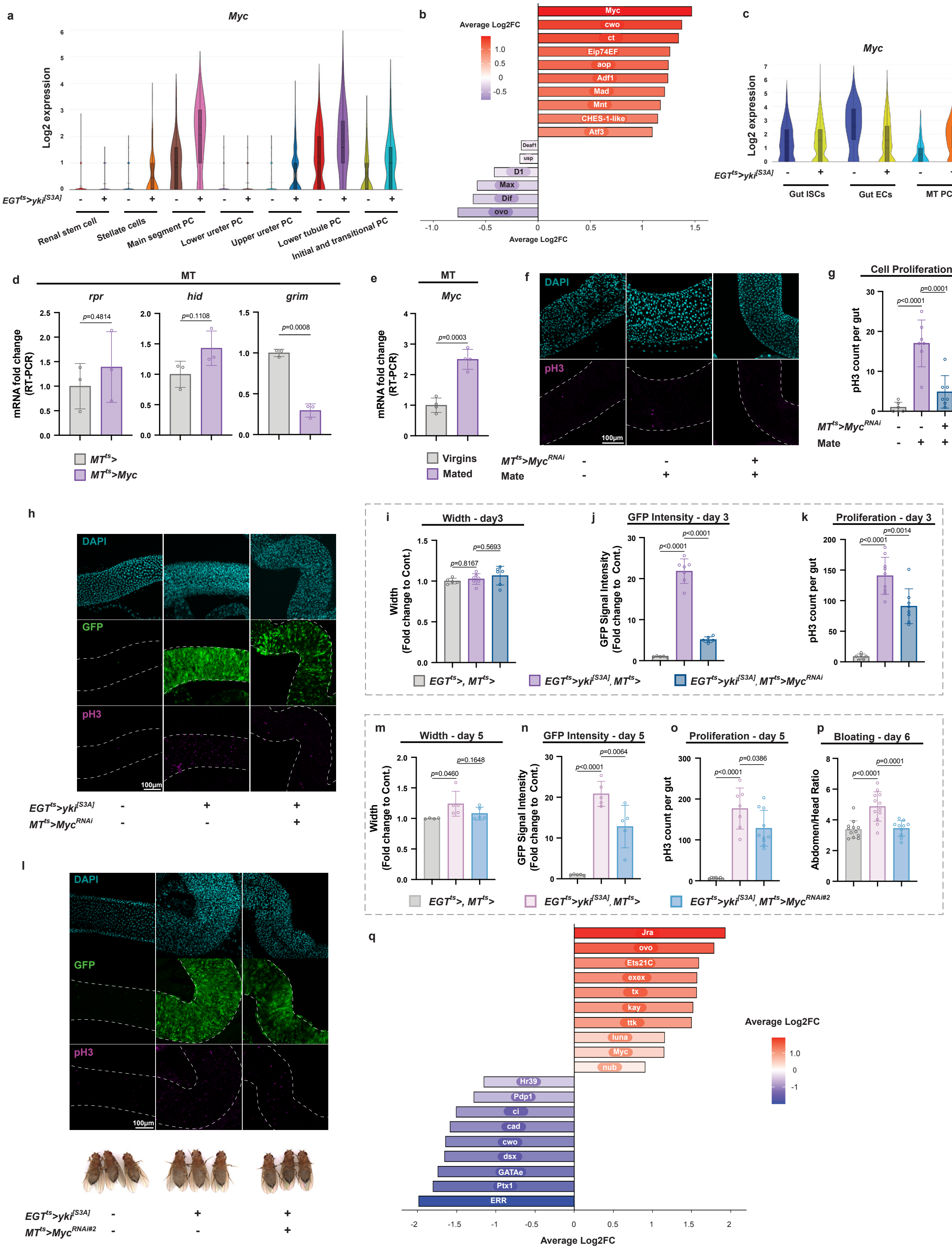

a
Myc
Log2 expression
EGT^ts>yki^[S3A]
Renal stem cell
Stellate cells
Main segment PC
Lower ureter PC
Upper ureter PC
Lower tubule PC
Initial and transitional PC
b
Average Log2FC
Myc
cwo
ct
Eip74EF
aop
Adf1
Mad
Mnt
CHES-1-like
Atf3
Deaf1
usp
D1
Max
Dif
ovo
c
Myc
Log2 expression
EGT^ts>yki^[S3A]
Gut ISCs
Gut ECs
MT PCs
d
MT
rpr
hid
grim
mRNA fold change (RT-PCR)
p=0.4814
p=0.1108
p=0.0008
MT^ts>
MT^ts>Myc
e
MT
Myc
mRNA fold change (RT-PCR)
p=0.0003
Virgins
Mated
f
DAPI
pH3
100μm
MT^ts>Myc^RNAi
Mate
g
Cell Proliferation
pH3 count per gut
p<0.0001
p=0.0001
MT^ts>Myc^RNAi
Mate
h
DAPI
GFP
pH3
100μm
EGT^ts>yki^[S3A]
MT^ts>Myc^RNAi
i
Width - day3
Width (Fold change to Cont.)
p=0.8167
p=0.5693
j
GFP Intensity - day 3
GFP Signal Intensity (Fold change to Cont.)
p<0.0001
p<0.0001
k
Proliferation - day 3
pH3 count per gut
p<0.0001
p=0.0014
EGT^ts>, MT^ts>
EGT^ts>yki^[S3A], MT^ts>
EGT^ts>yki^[S3A], MT^ts>Myc^RNAi
l
DAPI
GFP
pH3
100μm
EGT^ts>yki^[S3A]
MT^ts>Myc^RNAi#2
m
Width - day 5
Width (Fold change to Cont.)
p=0.0460
p=0.1648
n
GFP Intensity - day 5
GFP Signal Intensity (Fold change to Cont.)
p<0.0001
p=0.0064
o
Proliferation - day 5
pH3 count per gut
p<0.0001
p=0.0386
p
Bloating - day 6
Abdomen/Head Ratio
p<0.0001
p=0.0001
EGT^ts>, MT^ts>
EGT^ts>yki^[S3A], MT^ts>
EGT^ts>yki^[S3A], MT^ts>Myc^RNAi#2
q
Jra
ovo
Ets21C
exex
tx
kay
ttk
luna
Myc
nub
Hr39
Pdp1
ci
cad
cwo
dsx
GATAe
Ptx1
ERR
Average Log2FC

**Fig. S4 | Related to Figure 5. a,** Log2 expression of *Myc* in gut ISCs, gut ECs and MT PCs of control and Yki flies; data from published Yki MT snRNA-seq dataset. **b,** Differential regulon activity analysis of transcription factors in the MTs with published Yki fly snRNA-seq data. A high Log2fc corresponds to increased activity/expression of positively regulated target genes of the regulon in Yki versus control. **c,** Log2 expression of *dPANK4*, *Fbl*, and *Smvt* in gut ISCs, gut ECs and MT PCs of control and Yki flies; data from published Yki snRNA-seq dataset. **d,** qRT-PCR analysis of apoptosis genes (*rpr, hid,* and *grim*) mRNA levels in MTs from control and MT-specific *Myc* overexpression flies at day 7. n = 3. **e,** qRT-PCR analysis of *Myc* in the MTs of virgin or mated female flies. **f, g,** Representative images (**f**) and quantification (**g**) of pH3+ cells per midgut in virgin or mated female flies with or without MT *Myc* knockdown at day 10. In the pH3 panel, the guts are outlined with dashed lines. n = 6, 7 and 7 in $MT^{ts}$> (virgins), $MT^{ts}$> (mated), and $MT^{ts}$>*Myc-RNAi* (mated), respectively. **h,** Representative images of guts from control flies and Yki flies with or without MT-specific *Myc* knockdown at day 3. **i-k,** Quantification of gut width (**i**), GFP intensity (**j**), and pH3+ cell counts (**k**). n = 5, 8, and 6 in $EGT^{ts}$>, $MT^{ts}$>, $EGT^{ts}$>$yki^{[S3A]}$,$MT^{ts}$>, and $EGT^{ts}$>$yki^{[S3A]}$, $MT^{ts}$>*Myc-RNAi* in **i**, respectively. n = 5, 8, and 6 in $EGT^{ts}$>, $MT^{ts}$>, $EGT^{ts}$>$yki^{[S3A]}$,$MT^{ts}$>, and $EGT^{ts}$>$yki^{[S3A]}$, $MT^{ts}$>*Myc-RNAi* in **j**, respectively. n = 6, 9, and 9 in $EGT^{ts}$>, $MT^{ts}$>, $EGT^{ts}$>$yki^{[S3A]}$,$MT^{ts}$>, and $EGT^{ts}$>$yki^{[S3A]}$, $MT^{ts}$>*Myc-RNAi* in **k**, respectively. **l,** Representative images of guts and representative images of bloating phenotypes from control flies and Yki flies with or without MT-specific *Myc* knockdown at day 5 using independent fly lines. **m-o,** Quantification of gut width (**m**), GFP intensity (**n**), and pH3+ cell counts (**o**). n = 4, 5, and 5 in $EGT^{ts}$>, $MT^{ts}$>, $EGT^{ts}$>$yki^{[S3A]}$,$MT^{ts}$>, and $EGT^{ts}$>$yki^{[S3A]}$, $MT^{ts}$>*Myc-RNAi-#2* in **m**, respectively. n = 5, 5, and 5 in $EGT^{ts}$>, $MT^{ts}$>, $EGT^{ts}$>$yki^{[S3A]}$,$MT^{ts}$>, and $EGT^{ts}$>$yki^{[S3A]}$, $MT^{ts}$>*Myc-RNAi-#2* in **n**, respectively. n = 7, 10, and 10 in $EGT^{ts}$>, $MT^{ts}$>, $EGT^{ts}$>$yki^{[S3A]}$,$MT^{ts}$>, and $EGT^{ts}$>$yki^{[S3A]}$, $MT^{ts}$>*Myc-RNAi-#2* in **o**, respectively. **p,** Bloating quantification via abdomen-to-head ratio from panel **l**. n = 12, 12, and 10 in $EGT^{ts}$>, $MT^{ts}$>, $EGT^{ts}$>$yki^{[S3A]}$,$MT^{ts}$>, and $EGT^{ts}$>$yki^{[S3A]}$, $MT^{ts}$>*Myc-RNAi-#2*, respectively. **q,** Differential regulon activity analysis of transcription factors in MTs with published snRNA-seq data. A high Log2fc corresponds to increased activity/expression of positively regulated target genes of the regulon in MT-specific Pvr activation versus control. Statistical significance assessed by wilcox test (**b, q**), unpaired two-sided Student's *t*-test (**d, e**) and one-way ANOVA (**g, i-k, m-p**). Error bars indicate s.d., with means at the center. Source data are provided as a Source Data file.

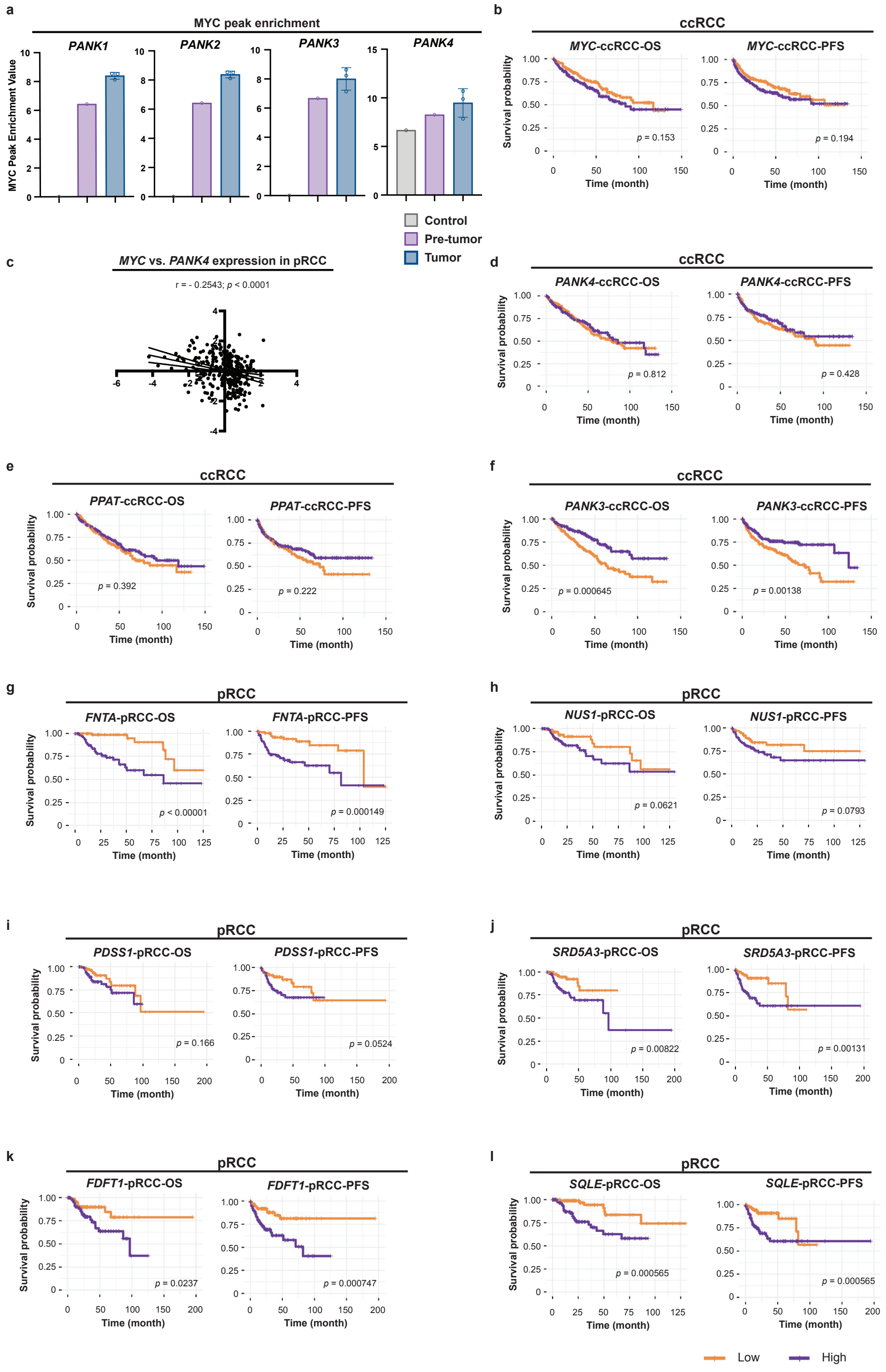
a
MYC peak enrichment
PANK1
PANK2
PANK3
PANK4
MYC Peak Enrichment Value
Control
Pre-tumor
Tumor
b
ccRCC
MYC-ccRCC-OS
MYC-ccRCC-PFS
p = 0.153
p = 0.194
c
MYC vs. PANK4 expression in pRCC
r = - 0.2543; p < 0.0001
d
ccRCC
PANK4-ccRCC-OS
PANK4-ccRCC-PFS
p = 0.812
p = 0.428
e
ccRCC
PPAT-ccRCC-OS
PPAT-ccRCC-PFS
p = 0.392
p = 0.222
f
ccRCC
PANK3-ccRCC-OS
PANK3-ccRCC-PFS
p = 0.000645
p = 0.00138
g
pRCC
FNTA-pRCC-OS
FNTA-pRCC-PFS
p < 0.00001
p = 0.000149
h
pRCC
NUS1-pRCC-OS
NUS1-pRCC-PFS
p = 0.0621
p = 0.0793
i
pRCC
PDSS1-pRCC-OS
PDSS1-pRCC-PFS
p = 0.166
p = 0.0524
j
pRCC
SRD5A3-pRCC-OS
SRD5A3-pRCC-PFS
p = 0.00822
p = 0.00131
k
pRCC
FDFT1-pRCC-OS
FDFT1-pRCC-PFS
p = 0.0237
p = 0.000747
l
pRCC
SQLE-pRCC-OS
SQLE-pRCC-PFS
p = 0.000565
p = 0.000565
Survival probability
Time (month)
Low
High

**Fig. S5 | Related to Figure 6. a,** MYC peak enrichment values in the promoter regions of PANK1-4 from published ChIP-seq datasets. Enrichment is calculated as Log2(ChIP - Input), where ChIP and Input represent the number of reads in the peak region normalized by total library size (in millions). A value of 0 is assigned to promoters with no peak called. **b,** Kaplan–Meier (KM) survival analyses of overall survival (OS) and progression-free survival (PFS) in 336 Clear Cell Renal Cell Carcinoma (ccRCC/KIRC) patients comparing low (bottom third) versus high (top third) expression of *MYC*. **c,** Correlation plot showing the negative relationship between MYC and PANK4 expression in 280 Papillary Renal Cell Carcinoma (pRCC/KIRP) patients from the TCGA PanCancer Atlas. **d-f,** KM survival analyses of OS and PFS in 512 ccRCC/KIRC patients comparing low (bottom third) versus high (top third) expression of *PANK4* (**d**), and *PANK3* (**e**), and *PPAT* (**f**); data from the TCGA PanCancer Atlas. **g-l,** KM survival analyses (OS and PFS) for 188 pRCC patients stratified by expression of *FNTA* (**g**), *NUS1* (**h**), *PDSS1* (**i**), *SRD5A3* (**j**), *FDFT1* (**k**), and *SQLE* (**l**), comparing bottom versus top expression tertiles. Statistical significance assessed by Pearson correlation analysis (**b**), and log-rank (Mantel-Cox) test for survival curves (**c-l**). Source data are provided as a Source Data file.

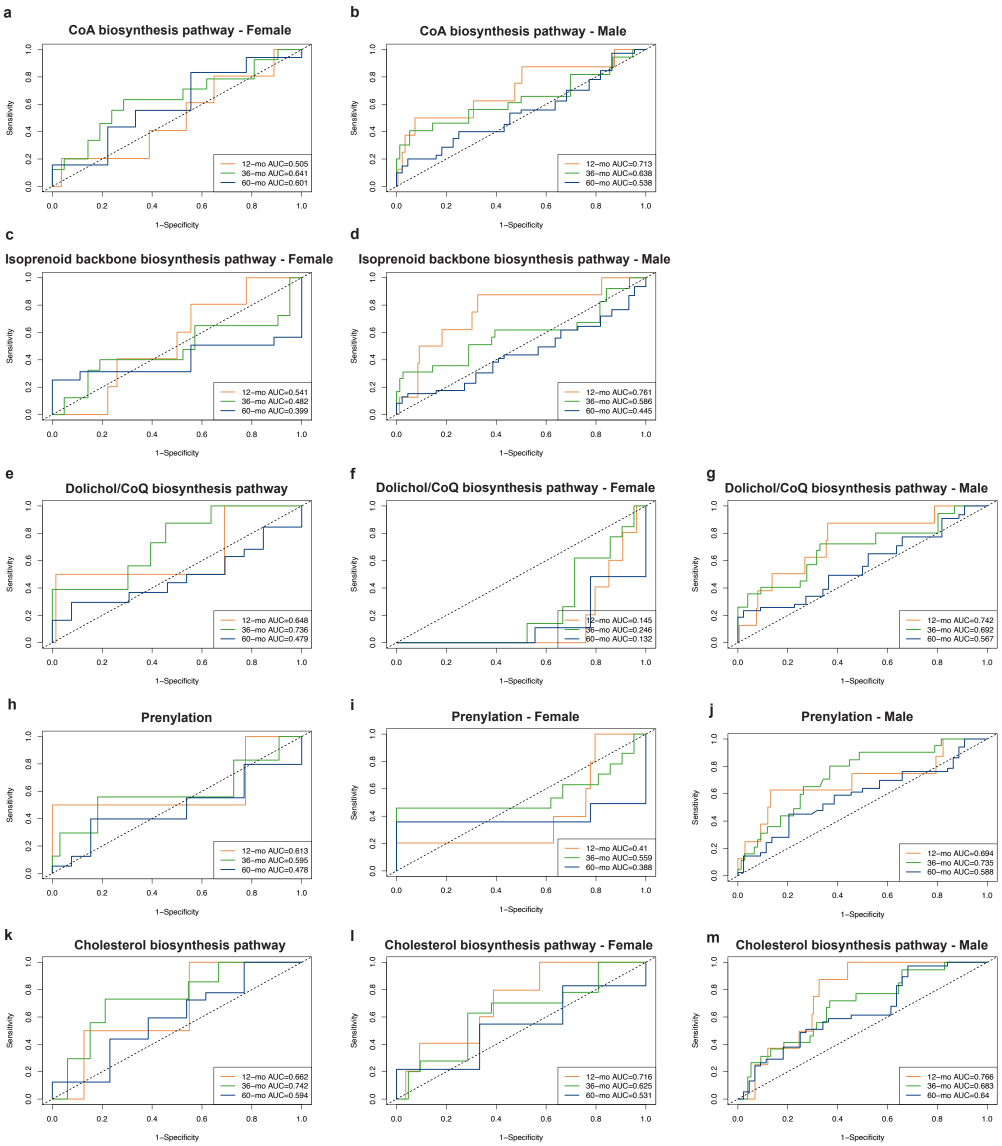
a
CoA biosynthesis pathway - Female
12−mo AUC=0.505
36−mo AUC=0.641
60−mo AUC=0.601
b
CoA biosynthesis pathway - Male
12−mo AUC=0.713
36−mo AUC=0.638
60−mo AUC=0.538
c
Isoprenoid backbone biosynthesis pathway - Female
12−mo AUC=0.541
36−mo AUC=0.482
60−mo AUC=0.399
d
Isoprenoid backbone biosynthesis pathway - Male
12−mo AUC=0.761
36−mo AUC=0.586
60−mo AUC=0.445
e
Dolichol/CoQ biosynthesis pathway
12−mo AUC=0.648
36−mo AUC=0.736
60−mo AUC=0.479
f
Dolichol/CoQ biosynthesis pathway - Female
12−mo AUC=0.145
36−mo AUC=0.246
60−mo AUC=0.132
g
Dolichol/CoQ biosynthesis pathway - Male
12−mo AUC=0.742
36−mo AUC=0.692
60−mo AUC=0.567
h
Prenylation
12−mo AUC=0.613
36−mo AUC=0.595
60−mo AUC=0.478
i
Prenylation - Female
12−mo AUC=0.41
36−mo AUC=0.559
60−mo AUC=0.388
j
Prenylation - Male
12−mo AUC=0.694
36−mo AUC=0.735
60−mo AUC=0.588
k
Cholesterol biosynthesis pathway
12−mo AUC=0.662
36−mo AUC=0.742
60−mo AUC=0.594
l
Cholesterol biosynthesis pathway - Female
12−mo AUC=0.716
36−mo AUC=0.625
60−mo AUC=0.531
m
Cholesterol biosynthesis pathway - Male
12−mo AUC=0.766
36−mo AUC=0.683
60−mo AUC=0.64
Sensitivity
1−Specificity

**Fig. S6 | Related to Figure 6. a-d,** Time-dependent ROC (receiver operating characteristic) curve analyses showing AUC (area under the curve) values at 12, 24, and 36 months of metastasis stage M0. Panels **a-b** show AUC for CoA biosynthesis gene signatures in female (**m**) (n = 72) and male (**n**) (n = 208) pRCC patients; panels **c-d** show AUC for isoprenoid backbone biosynthesis gene signatures in female (**o**) (n = 72) and male (**p**) (n = 208) patients. Pathway gene sets include: CoA biosynthesis: *PANK1-4, PPCS, PPCDC, PPAT* and *COASY*; isoprenoid backbone synthesis: *MVK, PMVK, MVD, IDI1, IDI2, FDPS* and *GGPS1*. **e-m,** Time-dependent ROC analysis of dolichol and ubiquinone (CoQ) biosynthesis gene signatures in pRCC patients showing AUC performance across both genders (**e**) (n = 91), females (**f**) (n = 72), and males (**g**) (n = 208). Pathway gene sets include: *DHDDS, NUS1, SRD5A1* and *PDSS1/2*. **h-j,** Time-dependent ROC analysis of protein prenylation gene signatures in pRCC patients showing AUC performance across both genders (**h**) (n = 91), females (**i**) (n = 72), and males (**j**) (n = 208). Pathway gene sets include: *FNTA, FNTB* and *PGGT1B*. **k-m,** Time-dependent ROC analysis of cholesterol biosynthesis gene signatures in pRCC patients showing AUC performance across both genders (**k**) (n = 91), females (**l**) (n = 72), and males (**m**) (n = 208). Pathway gene sets include: *FDFT1, SQLE, LSS, CYP51A1* and *DHCG7*. Statistical significance assessed by AUC analyses (**a-m**). Source data are provided as a Source Data file.

| Supplementary table1. Information about vitamin supplementation | | |
|---|---|---|
| | Supplier and Catalog number | Final concentration |
| Vitamin B1 (thiamine) | Sigma, T4625 | 0.2 mM |
| Vitamin B2 (riboflavin) | Sigma, R4500 | 0.1 mM |
| Vitamin B3 (nicotinic acid) | Sigma, N4126 | 5 mM |
| Vitamin B5 (pantothenate acid) | Sigma, 21210 | 2.5 mM |
| Vitamin B6 (pyridoxine) | Sigma, P9755 | 0.5 mM |
| Vitamin B7 (Biotin) | Sigma, B4501 | 0.01 mM |

**Supplementary table2. Transcription factor prediction**

| Query gene | TF FBgn | TF Symbol | Peak Count | Motif Count | Location | Protein-Protein Interactors | Genetic Interactors |
|---|---|---|---|---|---|---|---|
| ppcs | FBgn0259750 | ab | 0 | 3 | upstream | | |
| Ppcdc | FBgn0259750 | ab | 0 | 1 | upstream | | |
| fbl | FBgn0259750 | ab | 0 | 1 | upstream | | |
| Ppat-Dpck | FBgn0000022 | ac | 0 | 2 | upstream | da | sc, h (pubmed) |
| Ppat-Dpck | FBgn0000413 | ac | 0 | 6 | upstream | | |
| CG5828 | FBgn0000022 | ac | 0 | 2 | upstream | da | sc, h (pubmed) |
| fbl | FBgn0000022 | ac | 0 | 1 | upstream | | sc, ttk (pubmed) |
| Ppat-Dpck | FBgn0005694 | Aef1 | 0 | 2 | upstream | | |
| Ppcdc | FBgn0003270 | amos | 0 | 1 | upstream | | da (pubmed) |
| ppcs | FBgn0000097 | aop | 0 | 1 | upstream | | pnt, ttk (pubmed) |
| Ppcdc | FBgn0000097 | aop | 0 | 1 | upstream | | H (pubmed) |
| CG5828 | FBgn0000097 | aop | 0 | 1 | upstream | | H, pnt (pubmed) |
| Ppat-Dpck | FBgn0000137 | ase | 0 | 2 | upstream | da | |
| CG5828 | FBgn0000137 | ase | 0 | 2 | upstream | da | |
| fbl | FBgn0000137 | ase | 0 | 1 | upstream | | |
| ppcs | FBgn0004870 | bab1 | 0 | 3 | upstream | | |
| Ppcdc | FBgn0004870 | bab1 | 0 | 7 | upstream | | |
| Ppat-Dpck | FBgn0004870 | bab1 | 0 | 1 | upstream | | |
| Dpck | FBgn0004870 | bab1 | 0 | 2 | upstream | | |
| CG5828 | FBgn0004870 | bab1 | 0 | 2 | upstream | | |
| fbl | FBgn0004870 | bab1 | 0 | 2 | upstream | | |
| Ppcdc | FBgn0015602 | BEAF-32 | 0 | 1 | upstream | Dref | |
| CG5828 | FBgn0015602 | BEAF-32 | 0 | 2 | upstream | Dref | |
| fbl | FBgn0015602 | BEAF-32 | 0 | 1 | upstream | Dref | |
| ppcs | FBgn0045759 | bin | 0 | 2 | upstream | | |

| | | | | | | | |
|---|---|---|---|---|---|---|---|
| ppcs | FBgn0045759 | bin | 0 | 2 | intron | | |
| Dpck | FBgn0045759 | bin | 0 | 1 | upstream | prd, D | |
| CG5828 | FBgn0045759 | bin | 1 | 1 | upstream | D | |
| fbl | FBgn0045759 | bin | 0 | 4 | intron | D | |
| fbl | FBgn0045759 | bin | 0 | 1 | upstream | D | |
| ppcs | FBgn0035625 | Blimp-1 | 0 | 8 | upstream | | |
| ppcs | FBgn0035625 | Blimp-1 | 0 | 4 | intron | | |
| CG5828 | FBgn0035625 | Blimp-1 | 0 | 1 | upstream | | |
| ppcs | FBgn0004893 | bowl | 0 | 1 | upstream | | |
| CG5828 | FBgn0004893 | bowl | 0 | 1 | upstream | | |
| ppcs | FBgn0283451 | br | 2 | 0 | upstream | Rel, rib | Met (pubmed) |
| Ppat-Dpck | FBgn0283451 | br | 1 | 0 | upstream | rib | |
| Dpck | FBgn0283451 | br | 1 | 0 | upstream | rib | |
| Dpck | FBgn0000210 | br | 0 | 1 | upstream | | |
| CG5828 | FBgn0283451 | br | 1 | 0 | upstream | rib | |
| fbl | FBgn0283451 | br | 1 | 0 | upstream | rib | |
| fbl | FBgn0000210 | br | 0 | 4 | intron | | |
| ppcs | FBgn0000210 | br-PE | 0 | 1 | upstream | | |
| ppcs | FBgn0263108 | BtbVII | 0 | 2 | upstream | | |
| ppcs | FBgn0263108 | BtbVII | 0 | 1 | intron | | |
| Dpck | FBgn0263108 | BtbVII | 0 | 1 | upstream | | |
| CG5828 | FBgn0263108 | BtbVII | 0 | 1 | upstream | | |
| fbl | FBgn0263108 | BtbVII | 0 | 1 | upstream | | |
| fbl | FBgn0263108 | BtbVII | 0 | 1 | intron | | |
| Ppat-Dpck | FBgn0025679 | Bteb2 | 0 | 2 | upstream | | |
| fbl | FBgn0025679 | Bteb2 | 0 | 3 | upstream | | |
| Ppat-Dpck | FBgn0000286 | Cf2-PA | 0 | 1 | upstream | | |

| | | | | | | |
|---|---|---|---|---|---|---|
| Dpck | FBgn0000286 | Cf2-PA | 0 | 2 | upstream | bin |
| fbl | FBgn0000286 | Cf2-PA | 0 | 1 | upstream | bin |
| ppcs | FBgn0000286 | Cf2-PB | 0 | 8 | upstream | bin |
| fbl | FBgn0000286 | Cf2-PB | 0 | 1 | intron | bin |
| ppcs | FBgn0037446 | CG10267 | 0 | 2 | upstream | Dll, toy |
| Ppat-Dpck | FBgn0037446 | CG10267 | 0 | 2 | upstream | Dll |
| fbl | FBgn0037446 | CG10267 | 0 | 2 | upstream | |
| Ppcdc | FBgn0034945 | CG10904 | 0 | 1 | upstream | |
| fbl | FBgn0030532 | CG11071 | 0 | 1 | upstream | |
| ppcs | FBgn0035454 | CG12029 | 0 | 3 | intron | |
| Ppcdc | FBgn0035454 | CG12029 | 0 | 1 | upstream | |
| Ppat-Dpck | FBgn0035454 | CG12029 | 0 | 1 | upstream | |
| Dpck | FBgn0035454 | CG12029 | 0 | 3 | upstream | |
| CG5828 | FBgn0035454 | CG12029 | 0 | 3 | upstream | |
| fbl | FBgn0035454 | CG12029 | 0 | 2 | upstream | |
| CG5828 | FBgn0029957 | CG12155 | 0 | 1 | upstream | Trl |
| fbl | FBgn0029822 | CG12236 | 0 | 1 | intron | |
| ppcs | FBgn0037206 | CG12768 | 0 | 1 | intron | Mes2 |
| Dpck | FBgn0037206 | CG12768 | 0 | 1 | upstream | |
| Ppcdc | FBgn0035160 | CG13897 | 0 | 2 | upstream | |
| Ppat-Dpck | FBgn0035160 | CG13897 | 0 | 1 | upstream | |
| fbl | FBgn0035160 | CG13897 | 0 | 1 | upstream | |
| Ppat-Dpck | FBgn0035407 | CG14962 | 0 | 3 | upstream | |
| Dpck | FBgn0003715 | CG16778 | 0 | 1 | upstream | |
| CG5828 | FBgn0003715 | CG16778 | 0 | 1 | upstream | |
| fbl | FBgn0003715 | CG16778 | 0 | 2 | intron | |
| ppcs | FBgn0037735 | CG16899 | 0 | 1 | upstream | |

| | | | | | | | |
|---|---|---|---|---|---|---|---|
| ppcs | FBgn0037735 | CG16899 | 0 | 1 | intron | | |
| Dpck | FBgn0037735 | CG16899 | 0 | 1 | upstream | | |
| CG5828 | FBgn0037735 | CG16899 | 0 | 1 | upstream | | |
| fbl | FBgn0037735 | CG16899 | 0 | 3 | intron | | |
| fbl | FBgn0037735 | CG16899 | 0 | 1 | upstream | | |
| Ppat-Dpck | FBgn0035144 | CG17181 | 0 | 1 | upstream | Aef1 | |
| CG5828 | FBgn0035144 | CG17181 | 0 | 1 | upstream | pnt | |
| ppcs | FBgn0039905 | CG2052 | 0 | 3 | upstream | | |
| ppcs | FBgn0039905 | CG2052 | 0 | 4 | intron | | |
| Ppcdc | FBgn0039905 | CG2052 | 0 | 1 | intron | | |
| Ppat-Dpck | FBgn0039905 | CG2052 | 0 | 3 | upstream | | |
| Dpck | FBgn0039905 | CG2052 | 0 | 3 | upstream | | |
| CG5828 | FBgn0039905 | CG2052 | 0 | 11 | upstream | | |
| fbl | FBgn0039905 | CG2052 | 0 | 2 | upstream | | |
| Ppat-Dpck | FBgn0034946 | CG3065 | 0 | 3 | upstream | | |
| Dpck | FBgn0034946 | CG3065 | 0 | 8 | upstream | | |
| CG5828 | FBgn0034946 | CG3065 | 0 | 4 | upstream | | |
| fbl | FBgn0034946 | CG3065 | 0 | 4 | upstream | | |
| Ppcdc | FBgn0031375 | CG31670 | 0 | 1 | upstream | | klu (pubmed) |
| Dpck | FBgn0031375 | CG31670 | 0 | 4 | upstream | | klu, Doc2 (pubmed) |
| fbl | FBgn0031375 | CG31670 | 0 | 2 | intron | | |
| ppcs | FBgn0052105 | CG32105 | 0 | 1 | upstream | | |
| CG5828 | FBgn0053980 | CG33980 | 0 | 1 | upstream | | |
| Dpck | FBgn0031573 | CG3407 | 0 | 1 | upstream | | |
| ppcs | FBgn0036423 | CG3919 | 0 | 1 | intron | | |
| Ppat-Dpck | FBgn0036423 | CG3919 | 0 | 1 | upstream | | |
| fbl | FBgn0036423 | CG3919 | 0 | 2 | upstream | | |

| | | | | | | |
|---|---|---|---|---|---|---|
| fbl | FBgn0036423 | CG3919 | 0 | 1 | intron | |
| CG5828 | FBgn0038787 | CG4360 | 0 | 1 | upstream | |
| Ppat-Dpck | FBgn0038787 | CG4360-F1-3 | 0 | 4 | upstream | |
| ppcs | FBgn0030432 | CG4404 | 0 | 1 | intron | |
| Ppcdc | FBgn0030432 | CG4404 | 0 | 2 | upstream | |
| fbl | FBgn0030432 | CG4404 | 0 | 1 | upstream | |
| ppcs | FBgn0038766 | CG4854 | 0 | 2 | intron | |
| Ppcdc | FBgn0038766 | CG4854 | 0 | 1 | upstream | |
| Ppat-Dpck | FBgn0038766 | CG4854 | 0 | 2 | upstream | |
| CG5828 | FBgn0043457 | CG5180 | 0 | 1 | upstream | |
| ppcs | FBgn0039169 | CG5669 | 0 | 1 | upstream | |
| Dpck | FBgn0039169 | CG5669 | 0 | 6 | upstream | |
| CG5828 | FBgn0039169 | CG5669 | 0 | 3 | upstream | |
| fbl | FBgn0039169 | CG5669 | 0 | 3 | upstream | |
| ppcs | FBgn0032587 | CG5953 | 0 | 1 | upstream | knrl, sens |
| Ppcdc | FBgn0032587 | CG5953 | 0 | 1 | upstream | sens |
| Dpck | FBgn0032587 | CG5953 | 0 | 2 | upstream | knrl |
| CG5828 | FBgn0032587 | CG5953 | 0 | 1 | upstream | |
| fbl | FBgn0032587 | CG5953 | 0 | 2 | intron | |
| ppcs | FBgn0038316 | CG6276 | 0 | 2 | upstream | |
| ppcs | FBgn0038316 | CG6276 | 0 | 1 | intron | |
| Ppcdc | FBgn0038316 | CG6276 | 0 | 1 | upstream | |
| Dpck | FBgn0038316 | CG6276 | 0 | 2 | upstream | |
| CG5828 | FBgn0038316 | CG6276 | 0 | 2 | upstream | |
| fbl | FBgn0038316 | CG6276 | 0 | 1 | upstream | |
| Ppat-Dpck | FBgn0036179 | CG7368 | 0 | 1 | upstream | |
| ppcs | FBgn0033616 | CG7745 | 0 | 2 | upstream | |

| | | | | | | | |
|---|---|---|---|---|---|---|---|
| ppcs | FBgn0033616 | CG7745 | 0 | 1 | intron | | |
| Dpck | FBgn0033616 | CG7745 | 0 | 1 | upstream | | |
| CG5828 | FBgn0033616 | CG7745 | 0 | 1 | upstream | | |
| Ppat-Dpck | FBgn0039740 | CG7928 | 0 | 1 | upstream | | |
| ppcs | FBgn0035824 | CG8281 | 0 | 1 | upstream | | |
| ppcs | FBgn0035824 | CG8281 | 0 | 1 | intron | | |
| fbl | FBgn0037722 | CG8319 | 0 | 2 | upstream | | |
| Ppcdc | FBgn0036900 | CG8765 | 0 | 1 | upstream | | |
| CG5828 | FBgn0036900 | CG8765 | 0 | 1 | upstream | | |
| fbl | FBgn0036900 | CG8765 | 0 | 2 | intron | | |
| ppcs | FBgn0034810 | CG9895 | 0 | 2 | intron | | |
| Ppat-Dpck | FBgn0034810 | CG9895 | 0 | 1 | upstream | | |
| Dpck | FBgn0034810 | CG9895 | 0 | 6 | upstream | | |
| CG5828 | FBgn0034810 | CG9895 | 0 | 3 | upstream | | |
| fbl | FBgn0034810 | CG9895 | 0 | 2 | upstream | | |
| ppcs | FBgn0086758 | chinmo | 0 | 3 | intron | | |
| Ppcdc | FBgn0086758 | chinmo | 0 | 2 | upstream | | |
| CG5828 | FBgn0086758 | chinmo | 0 | 1 | upstream | | |
| fbl | FBgn0086758 | chinmo | 0 | 1 | intron | | |
| ppcs | FBgn0000370 | Crc | 0 | 3 | upstream | EcR | EcR (pubmed) |
| ppcs | FBgn0036126 | Crc | 0 | 2 | upstream | Xrp1, crc | |
| Ppat-Dpck | FBgn0000370 | Crc | 0 | 1 | upstream | EcR | EcR (pubmed) |
| ppcs | FBgn0014143 | croc | 0 | 1 | upstream | opa | |
| ppcs | FBgn0014143 | croc | 0 | 1 | intron | opa | |
| Ppcdc | FBgn0014143 | croc | 0 | 2 | upstream | | |
| Ppat-Dpck | FBgn0014143 | croc | 0 | 1 | upstream | | |
| Dpck | FBgn0014143 | croc | 0 | 1 | upstream | prd | |

| | | | | | | | |
|---|---|---|---|---|---|---|---|
| fbl | FBgn0014143 | croc | 0 | 1 | upstream | | |
| ppcs | FBgn0020309 | crol-F7-16 | 0 | 1 | upstream | | |
| CG5828 | FBgn0020309 | crol-F7-16 | 0 | 1 | upstream | | |
| CG5828 | FBgn0001994 | crp | 0 | 1 | upstream | | |
| ppcs | FBgn0023094 | cyc | 1 | 0 | intron | | |
| Ppcdc | FBgn0023094 | cyc | 1 | 0 | upstream | | |
| Ppat-Dpck | FBgn0023094 | cyc | 1 | 0 | upstream | | |
| Dpck | FBgn0023094 | cyc | 3 | 0 | upstream | | |
| CG5828 | FBgn0023094 | cyc | 2 | 0 | upstream | | |
| fbl | FBgn0023094 | cyc | 4 | 0 | intron | | |
| Ppcdc | FBgn0000411 | D | 3 | 0 | upstream | vnd | |
| Ppat-Dpck | FBgn0000411 | D | 3 | 0 | upstream | | |
| Dpck | FBgn0000411 | D | 2 | 0 | upstream | Doc1, bin | |
| CG5828 | FBgn0000411 | D | 2 | 0 | upstream | bin | |
| fbl | FBgn0000411 | D | 1 | 0 | upstream | vnd, ftz-f1, ttk, bin | |
| ppcs | FBgn0022935 | D19A | 0 | 2 | upstream | | |
| Dpck | FBgn0022935 | D19A | 0 | 1 | upstream | | |
| CG5828 | FBgn0022935 | D19A | 0 | 1 | upstream | | |
| fbl | FBgn0022935 | D19A | 0 | 1 | upstream | | |
| fbl | FBgn0022935 | D19A | 0 | 1 | intron | | |
| ppcs | FBgn0022699 | D19B-F10-12 | 0 | 1 | intron | | |
| ppcs | FBgn0267821 | da | 1 | 0 | upstream | ey, Fer3 | |
| ppcs | FBgn0000413 | da | 0 | 1 | upstream | | |
| ppcs | FBgn0000413 | da | 0 | 4 | intron | | |
| Ppcdc | FBgn0267821 | da | 1 | 0 | upstream | dimm, ey, Fer3 | amos (pubmed) |
| Ppat-Dpck | FBgn0267821 | da | 3 | 0 | upstream | ac, sc, l(1)sc, ase, HLH54F, ey, Fer3 | sc, l(1)sc (pubmed) |
| CG5828 | FBgn0000413 | da | 0 | 4 | upstream | | |

| | | | | | | | |
|---|---|---|---|---|---|---|---|
| CG5828 | FBgn0267821 | da | 2 | 0 | upstream | ac, sc, l(1)sc, ase, Fer3 | sc, l(1)sc (pubmed) |
| Ppcdc | FBgn0000413 | dei | 0 | 5 | upstream | | |
| Ppcdc | FBgn0008649 | dei | 0 | 2 | upstream | | |
| Ppcdc | FBgn0023091 | dimm | 0 | 1 | upstream | da, sqz | |
| Dpck | FBgn0000413 | dimm | 0 | 1 | upstream | | |
| Dpck | FBgn0023091 | dimm | 0 | 2 | upstream | sqz | |
| ppcs | FBgn0040465 | Dip3 | 0 | 1 | upstream | CG4854 | |
| Ppcdc | FBgn0040465 | Dip3 | 0 | 1 | intron | dl, CG4854 | |
| Ppat-Dpck | FBgn0040465 | Dip3 | 0 | 2 | upstream | CG4854 | |
| Dpck | FBgn0040465 | Dip3 | 0 | 2 | upstream | | |
| CG5828 | FBgn0040465 | Dip3 | 0 | 2 | upstream | | |
| Ppcdc | FBgn0000462 | dl | 0 | 2 | upstream | | |
| ppcs | FBgn0000157 | Dll | 1 | 0 | upstream | Dref | |
| Ppat-Dpck | FBgn0000157 | Dll | 2 | 0 | upstream | Dref | |
| CG5828 | FBgn0000157 | Dll | 1 | 0 | upstream | Dref | |
| Dpck | FBgn0028789 | Doc1 | 0 | 1 | intron | Doc2, D | Doc2 (pubmed) |
| Dpck | FBgn0035956 | Doc2 | 0 | 1 | intron | Doc1 | Doc1 (pubmed) |
| Dpck | FBgn0035954 | Doc3 | 0 | 1 | intron | | |
| Ppat-Dpck | FBgn0010109 | dpn | 0 | 2 | upstream | h, Hey | |
| ppcs | FBgn0000492 | Dr | 0 | 1 | upstream | | |
| ppcs | FBgn0015664 | Dref | 0 | 2 | upstream | Dll | |
| ppcs | FBgn0015664 | Dref | 0 | 1 | intron | Dll | |
| Ppcdc | FBgn0015664 | Dref | 0 | 1 | upstream | BEAF-32 | |
| Ppat-Dpck | FBgn0015664 | Dref | 0 | 3 | upstream | Dll | |
| CG5828 | FBgn0015664 | Dref | 0 | 2 | upstream | BEAF-32, Dll | |
| fbl | FBgn0015664 | Dref | 0 | 1 | upstream | BEAF-32 | |
| ppcs | FBgn0015381 | dsf | 0 | 1 | upstream | | |

| | | | | | | | |
|---|---|---|---|---|---|---|---|
| Ppat-Dpck | FBgn0000504 | dsx-F | 0 | 1 | upstream | ey | |
| CG5828 | FBgn0000504 | dsx-F | 0 | 1 | upstream | | |
| ppcs | FBgn0039411 | dys | 0 | 4 | upstream | tgo | |
| Dpck | FBgn0039411 | dys | 0 | 1 | upstream | tgo | |
| CG5828 | FBgn0015014 | dys | 0 | 1 | upstream | | |
| CG5828 | FBgn0039411 | dys | 0 | 2 | upstream | | |
| Ppcdc | FBgn0000591 | E(spl) | 0 | 1 | upstream | da, h | Dl, H (pubmed) |
| ppcs | FBgn0000546 | EcR | 0 | 1 | upstream | usp, Met, tai, crc | Hr39, crc (pubmed) |
| ppcs | FBgn0000546 | EcR | 0 | 1 | intron | usp, Met, tai, crc | Hr39, crc (pubmed) |
| Ppat-Dpck | FBgn0000546 | EcR | 2 | 0 | upstream | usp, crc | crc (pubmed) |
| ppcs | FBgn0000560 | eg | 0 | 2 | upstream | | |
| ppcs | FBgn0000568 | Eip75B | 0 | 2 | intron | Hr51 | Kr (pubmed) |
| Dpck | FBgn0000568 | Eip75B | 0 | 1 | upstream | | |
| fbl | FBgn0000568 | Eip75B | 0 | 2 | upstream | Hr51 | |
| ppcs | FBgn0004865 | Eip78C | 0 | 1 | upstream | | |
| ppcs | FBgn0004865 | Eip78C | 0 | 1 | intron | | |
| ppcs | FBgn0013948 | Eip93F | 0 | 3 | upstream | | |
| ppcs | FBgn0013948 | Eip93F | 0 | 1 | intron | | |
| Dpck | FBgn0013948 | Eip93F | 0 | 1 | upstream | | |
| CG5828 | FBgn0013948 | Eip93F | 0 | 2 | upstream | | |
| ppcs | FBgn0035849 | ERR | 0 | 1 | intron | | |
| Ppat-Dpck | FBgn0001981 | esg-F3-5 | 0 | 2 | upstream | | ase (pubmed) |
| CG5828 | FBgn0001981 | esg-F3-5 | 0 | 3 | upstream | Sp1 | ase (pubmed) |
| fbl | FBgn0001981 | esg-F3-5 | 0 | 2 | upstream | Sp1 | ase (pubmed) |
| ppcs | FBgn0000591 | Espl | 0 | 2 | intron | da | |
| Ppat-Dpck | FBgn0000591 | Espl | 0 | 3 | upstream | da, dpn, h | l(1)sc, H (pubmed) |
| Dpck | FBgn0000591 | Espl | 0 | 1 | upstream | h | H (pubmed) |

| | | | | | | | |
|---|---|---|---|---|---|---|---|
| fbl | FBgn0000591 | Espl | 0 | 2 | intron | | l(1)sc (pubmed) |
| ppcs | FBgn0005660 | Ets21c | 0 | 2 | upstream | | |
| Ppcdc | FBgn0005660 | Ets21c | 0 | 1 | upstream | | |
| CG5828 | FBgn0005660 | Ets21c | 0 | 2 | upstream | | |
| ppcs | FBgn0005658 | Ets65A | 0 | 1 | upstream | | |
| Ppcdc | FBgn0005658 | Ets65A | 0 | 1 | upstream | | |
| CG5828 | FBgn0005658 | Ets65A | 0 | 1 | upstream | | |
| ppcs | FBgn0039225 | Ets96B | 0 | 1 | upstream | | |
| CG5828 | FBgn0039225 | Ets96B | 0 | 1 | upstream | | |
| ppcs | FBgn0004510 | Ets97D | 0 | 2 | upstream | Myc | |
| CG5828 | FBgn0004510 | Ets97D | 0 | 2 | upstream | Myc | |
| fbl | FBgn0005659 | Ets98B | 0 | 1 | upstream | | |
| ppcs | FBgn0005558 | ey | 0 | 3 | intron | da | toy (pubmed) |
| Ppcdc | FBgn0005558 | ey | 0 | 1 | upstream | da, hth | toy (pubmed) |
| Ppat-Dpck | FBgn0005558 | ey | 0 | 2 | upstream | da, hth | |
| fbl | FBgn0005558 | ey | 0 | 1 | upstream | | |
| ppcs | FBgn0037475 | Fer1 | 0 | 2 | intron | | |
| Ppcdc | FBgn0037475 | Fer1 | 0 | 2 | upstream | | |
| CG5828 | FBgn0037475 | Fer1 | 0 | 2 | upstream | | |
| ppcs | FBgn0038402 | Fer2 | 0 | 1 | upstream | | |
| CG5828 | FBgn0038402 | Fer2 | 0 | 2 | upstream | | |
| ppcs | FBgn0037937 | Fer3 | 1 | 0 | upstream | da | |
| Ppcdc | FBgn0037937 | Fer3 | 0 | 2 | upstream | da | |
| Ppat-Dpck | FBgn0037937 | Fer3 | 2 | 0 | upstream | da | |
| CG5828 | FBgn0037937 | Fer3 | 1 | 0 | upstream | da | |
| ppcs | FBgn0000659 | fkh | 0 | 1 | upstream | | |
| ppcs | FBgn0000659 | fkh | 0 | 1 | intron | | |

| | | | | | | | |
|---|---|---|---|---|---|---|---|
| Dpck | FBgn0000659 | fkh | 0 | 3 | upstream | | |
| fbl | FBgn0000659 | fkh | 0 | 2 | intron | | |
| fbl | FBgn0001078 | ftz-f1 | 0 | 1 | upstream | D | |
| ppcs | FBgn0032223 | GATAd | 0 | 2 | upstream | | |
| ppcs | FBgn0032223 | GATAd | 0 | 1 | intron | | |
| Ppat-Dpck | FBgn0032223 | GATAd | 0 | 1 | upstream | | |
| CG5828 | FBgn0032223 | GATAd | 0 | 1 | upstream | | |
| fbl | FBgn0032223 | GATAd | 0 | 2 | intron | | |
| Ppcdc | FBgn0038391 | GATAe | 1 | 0 | upstream | | |
| ppcs | FBgn0261703 | gce | 0 | 1 | upstream | Met | Met (pubmed) |
| Ppcdc | FBgn0261703 | gce | 0 | 1 | upstream | | |
| Ppat-Dpck | FBgn0261703 | gce | 0 | 1 | upstream | | |
| ppcs | FBgn0004618 | gl | 0 | 1 | upstream | | |
| Dpck | FBgn0004618 | gl | 0 | 2 | upstream | | |
| ppcs | FBgn0259211 | grh | 1 | 0 | upstream | Rel | |
| Ppat-Dpck | FBgn0259211 | grh | 1 | 0 | upstream | | |
| CG5828 | FBgn0259211 | grh | 1 | 0 | upstream | | |
| Dpck | FBgn0001138 | grn | 0 | 1 | upstream | | |
| Ppat-Dpck | FBgn0001148 | gsb | 0 | 1 | upstream | | |
| ppcs | FBgn0001150 | gt | 0 | 1 | upstream | ttk | |
| Dpck | FBgn0001150 | gt | 0 | 1 | upstream | | |
| CG5828 | FBgn0001150 | gt | 1 | 2 | upstream | | |
| fbl | FBgn0001150 | gt | 0 | 1 | intron | ttk | |
| Ppcdc | FBgn0001168 | h | 1 | 0 | upstream | | |
| Ppat-Dpck | FBgn0001168 | h | 1 | 0 | upstream | dpn | ac, sc, l(1)sc (pubmed) |
| Dpck | FBgn0001168 | h | 0 | 1 | upstream | | |
| CG5828 | FBgn0001168 | h | 2 | 0 | upstream | | ac, sc, l(1)sc (pubmed) |

| | | | | | | | |
|---|---|---|---|---|---|---|---|
| Ppat-Dpck | FBgn0032209 | Hand | 0 | 2 | upstream | | |
| fbl | FBgn0032209 | Hand | 0 | 1 | intron | | |
| ppcs | FBgn0001180 | hb | 0 | 13 | upstream | | Kr (pubmed) |
| ppcs | FBgn0001180 | hb | 0 | 5 | intron | | Kr (pubmed) |
| Ppcdc | FBgn0001180 | hb | 0 | 1 | upstream | | |
| Ppcdc | FBgn0001180 | hb | 0 | 3 | intron | | |
| Ppat-Dpck | FBgn0001180 | hb | 0 | 6 | upstream | | |
| Dpck | FBgn0001180 | hb | 0 | 16 | upstream | | |
| CG5828 | FBgn0001180 | hb | 0 | 19 | upstream | | Kr (pubmed) |
| fbl | FBgn0001180 | hb | 0 | 3 | upstream | | |
| fbl | FBgn0001180 | hb | 0 | 3 | intron | | |
| fbl | FBgn0001185 | her | 0 | 1 | intron | | |
| Ppcdc | FBgn0027788 | Hey | 0 | 1 | upstream | | |
| Ppat-Dpck | FBgn0027788 | Hey | 0 | 1 | upstream | dpn | |
| ppcs | FBgn0001204 | hkb | 0 | 2 | intron | | |
| Ppcdc | FBgn0001204 | hkb | 0 | 2 | upstream | | |
| Dpck | FBgn0001204 | hkb | 0 | 2 | upstream | | |
| fbl | FBgn0261283 | HLH106 | 0 | 1 | intron | | |
| Ppat-Dpck | FBgn0011277 | HLH4C | 0 | 2 | upstream | | |
| Ppat-Dpck | FBgn0022740 | HLH54F | 0 | 3 | upstream | da | |
| Ppcdc | FBgn0002609 | HLHm3 | 0 | 2 | upstream | h | |
| Ppat-Dpck | FBgn0002609 | HLHm3 | 0 | 2 | upstream | ac, sc, h | |
| Ppcdc | FBgn0002631 | HLHm5 | 0 | 1 | upstream | da | |
| Ppat-Dpck | FBgn0002631 | HLHm5 | 0 | 3 | upstream | da | |
| Ppat-Dpck | FBgn0002633 | HLHm7 | 0 | 2 | upstream | ac, sc, da | |
| ppcs | FBgn0002733 | HLHmbeta | 0 | 1 | upstream | da, eg | |
| Ppcdc | FBgn0002733 | HLHmbeta | 0 | 1 | upstream | da, tap | |

| | | | | | | | |
|---|---|---|---|---|---|---|---|
| Ppat-Dpck | FBgn0002733 | HLHmbeta | 0 | 1 | upstream | sc, da | sc (pubmed) |
| Dpck | FBgn0002733 | HLHmbeta | 0 | 2 | upstream | | |
| CG5828 | FBgn0002733 | HLHmbeta | 0 | 2 | upstream | sc, da | sc (pubmed) |
| ppcs | FBgn0002734 | HLHmd | 0 | 2 | upstream | | |
| ppcs | FBgn0002734 | HLHmd | 0 | 1 | intron | | |
| Ppcdc | FBgn0002734 | HLHmdelta | 0 | 2 | upstream | | |
| Ppat-Dpck | FBgn0002734 | HLHmdelta | 0 | 3 | upstream | | |
| ppcs | FBgn0002735 | HLHmg | 0 | 2 | upstream | da | |
| ppcs | FBgn0002735 | HLHmg | 0 | 1 | intron | da | |
| Ppcdc | FBgn0002735 | HLHmg | 0 | 1 | upstream | da | |
| Ppat-Dpck | FBgn0002735 | HLHmgamma | 0 | 3 | upstream | sc, da, dpn, ac | |
| ppcs | FBgn0004914 | Hnf4 | 0 | 1 | upstream | | |
| Ppcdc | FBgn0004914 | Hnf4 | 0 | 1 | upstream | | |
| CG5828 | FBgn0004914 | Hnf4 | 0 | 1 | upstream | | |
| ppcs | FBgn0261239 | Hr39 | 0 | 1 | intron | | EcR (pubmed) |
| ppcs | FBgn0000448 | Hr46 | 0 | 2 | upstream | toy | |
| ppcs | FBgn0000448 | Hr46 | 0 | 2 | intron | toy | |
| Ppcdc | FBgn0000448 | Hr46 | 0 | 2 | upstream | toy | |
| ppcs | FBgn0034012 | Hr51 | 0 | 1 | intron | Eip75B | |
| Ppat-Dpck | FBgn0034012 | Hr51 | 0 | 1 | upstream | | |
| CG5828 | FBgn0034012 | Hr51 | 0 | 2 | upstream | | |
| fbl | FBgn0034012 | Hr51 | 0 | 1 | upstream | Eip75B | |
| ppcs | FBgn0015239 | Hr78 | 0 | 1 | upstream | | |
| Ppcdc | FBgn0015239 | Hr78 | 0 | 1 | upstream | | |
| Ppat-Dpck | FBgn0015239 | Hr78 | 1 | 0 | upstream | | |
| CG5828 | FBgn0015239 | Hr78 | 1 | 1 | upstream | | |
| Dpck | FBgn0037436 | Hr83 | 0 | 2 | upstream | kni | |

| | | | | | | | |
|---|---|---|---|---|---|---|---|
| fbl | FBgn0037436 | Hr83 | 0 | 1 | intron | | |
| Ppcdc | FBgn0001235 | hth | 1 | 0 | upstream | ey | |
| Ppat-Dpck | FBgn0001235 | hth | 1 | 0 | upstream | ey | |
| ppcs | FBgn0039350 | jigr1 | 0 | 2 | upstream | | |
| Ppat-Dpck | FBgn0039350 | jigr1 | 0 | 2 | upstream | | |
| Dpck | FBgn0039350 | jigr1 | 0 | 2 | upstream | | |
| CG5828 | FBgn0039350 | jigr1 | 0 | 2 | upstream | | |
| fbl | FBgn0039350 | jigr1 | 0 | 1 | intron | | |
| ppcs | FBgn0027339 | jim | 0 | 6 | upstream | | |
| Ppcdc | FBgn0027339 | jim | 0 | 1 | upstream | | |
| Ppat-Dpck | FBgn0027339 | jim | 0 | 3 | upstream | | |
| Dpck | FBgn0027339 | jim | 0 | 1 | upstream | | |
| CG5828 | FBgn0027339 | jim | 0 | 15 | upstream | | |
| fbl | FBgn0027339 | jim | 0 | 4 | upstream | | |
| fbl | FBgn0001291 | Jra | 0 | 1 | intron | kay | kay (pubmed) |
| ppcs | FBgn0001291 | kay | 0 | 3 | upstream | Stat92E, kay | pnt, kay (pubmed) |
| ppcs | FBgn0001297 | kay | 0 | 3 | upstream | vri | |
| Dpck | FBgn0001291 | kay | 0 | 1 | upstream | kay | kay (pubmed) |
| Dpck | FBgn0001297 | kay | 0 | 1 | upstream | | |
| fbl | FBgn0001297 | kay | 0 | 1 | intron | vri, Jra | Jra (pubmed) |
| ppcs | FBgn0011236 | ken | 0 | 2 | upstream | | |
| ppcs | FBgn0011236 | ken | 0 | 1 | intron | | |
| Ppcdc | FBgn0011236 | ken | 0 | 1 | upstream | | |
| CG5828 | FBgn0011236 | ken | 0 | 1 | upstream | Trl | |
| fbl | FBgn0011236 | ken | 0 | 2 | upstream | | |
| ppcs | FBgn0013469 | klu | 0 | 3 | intron | | |
| Ppcdc | FBgn0013469 | klu | 0 | 7 | upstream | | H (pubmed) |

| | | | | | | | |
|---|---|---|---|---|---|---|---|
| Dpck | FBgn0013469 | klu | 0 | 3 | upstream | | H (pubmed) |
| CG5828 | FBgn0013469 | klu | 0 | 4 | upstream | | H (pubmed) |
| ppcs | FBgn0001320 | kni | 0 | 3 | upstream | | knrl (pubmed) |
| Dpck | FBgn0001320 | kni | 0 | 2 | intron | Hr83 | knrl (pubmed) |
| ppcs | FBgn0001323 | knrl | 0 | 3 | upstream | CG5953 | kni (pubmed) |
| Dpck | FBgn0001323 | knrl | 0 | 2 | intron | CG5953 | kni (pubmed) |
| ppcs | FBgn0001325 | Kr | 0 | 1 | upstream | | Eip75B, hb (pubmed) |
| CG5828 | FBgn0001325 | Kr | 1 | 0 | upstream | | hb (pubmed) |
| Ppat-Dpck | FBgn0002561 | l(1)sc | 0 | 2 | upstream | da | da, h (pubmed) |
| CG5828 | FBgn0002561 | l(1)sc | 0 | 2 | upstream | da | da, h (pubmed) |
| fbl | FBgn0002561 | l(1)sc | 0 | 1 | upstream | | |
| ppcs | FBgn0086910 | l(3)neo38 | 0 | 1 | upstream | | |
| ppcs | FBgn0086910 | l(3)neo38 | 0 | 1 | intron | | |
| Ppat-Dpck | FBgn0086910 | l(3)neo38 | 0 | 1 | upstream | | |
| CG5828 | FBgn0040918 | Lag1 | 0 | 1 | upstream | | |
| ppcs | FBgn0039039 | lmd | 0 | 2 | upstream | | |
| CG5828 | FBgn0039039 | lmd | 0 | 2 | upstream | | |
| Ppcdc | FBgn0005630 | lola | 0 | 1 | upstream | | |
| Dpck | FBgn0005630 | lola-PJ | 0 | 1 | upstream | | |
| Dpck | FBgn0040765 | luna | 0 | 2 | upstream | | |
| CG5828 | FBgn0040765 | luna | 0 | 1 | upstream | | |
| ppcs | FBgn0017578 | Max | 0 | 1 | upstream | Myc | Myc (pubmed) |
| Ppat-Dpck | FBgn0017578 | Max | 0 | 1 | upstream | Myc | Myc (pubmed) |
| Dpck | FBgn0023215 | Max | 0 | 1 | upstream | Max | Myc (pubmed) |
| fbl | FBgn0023215 | Max | 0 | 1 | upstream | Max | Myc (pubmed) |
| ppcs | FBgn0011655 | Med | 1 | 0 | upstream | | |
| Ppcdc | FBgn0011655 | Med | 3 | 0 | upstream | | |

| | | | | | | | |
|---|---|---|---|---|---|---|---|
| Ppat-Dpck | FBgn0011655 | Med | 3 | 0 | upstream | | |
| Dpck | FBgn0011655 | Med | 3 | 0 | upstream | | |
| CG5828 | FBgn0011655 | Med | 2 | 0 | upstream | | |
| ppcs | FBgn0037207 | Mes2 | 0 | 2 | upstream | CG12768 | |
| fbl | FBgn0037207 | Mes2 | 0 | 1 | upstream | | |
| fbl | FBgn0037207 | Mes2 | 0 | 1 | intron | | |
| ppcs | FBgn0002723 | Met | 0 | 1 | upstream | usp, EcR, gce | gce, br (pubmed) |
| ppcs | FBgn0023076 | Met | 0 | 2 | upstream | cyc | |
| ppcs | FBgn0002723 | Met | 0 | 1 | intron | usp, EcR, gce | gce, br (pubmed) |
| ppcs | FBgn0023076 | Met | 0 | 3 | intron | cyc | |
| ppcs | FBgn0032940 | Mio | 0 | 1 | upstream | | |
| ppcs | FBgn0039509 | Mio | 0 | 1 | upstream | | |
| Ppat-Dpck | FBgn0032940 | Mio | 0 | 1 | upstream | | |
| Ppat-Dpck | FBgn0039509 | Mio | 0 | 1 | upstream | | |
| ppcs | FBgn0262656 | Myc | 2 | 1 | upstream | Max, Ets97D | Max (pubmed) |
| Ppcdc | FBgn0262656 | Myc | 1 | 0 | upstream | | |
| Ppat-Dpck | FBgn0262656 | Myc | 0 | 1 | upstream | Max | Max (pubmed) |
| Dpck | FBgn0262656 | Myc | 2 | 0 | upstream | Max | prd, Max (pubmed) |
| CG5828 | FBgn0262656 | Myc | 1 | 0 | upstream | Ets97D | |
| fbl | FBgn0262656 | Myc | 1 | 0 | upstream | Max | Max (pubmed) |
| Ppcdc | FBgn0002922 | nau | 0 | 2 | upstream | | |
| ppcs | FBgn0030505 | NFAT | 0 | 2 | intron | | pnr (pubmed) |
| ppcs | FBgn0085424 | nub | 0 | 1 | upstream | pdm2 | |
| ppcs | FBgn0085424 | nub | 0 | 11 | intron | pdm2 | |
| ppcs | FBgn0002985 | odd | 0 | 1 | upstream | | |
| fbl | FBgn0032651 | Oli | 0 | 1 | upstream | | |
| ppcs | FBgn0003002 | Opa | 0 | 1 | upstream | croc | |

| | | | | | | | |
|---|---|---|---|---|---|---|---|
| Ppat-Dpck | FBgn0003028 | ovo | 0 | 1 | upstream | | |
| ppcs | FBgn0004394 | pdm2 | 0 | 6 | intron | nub | |
| Ppcdc | FBgn0016694 | Pdp1 | 0 | 6 | upstream | | |
| Dpck | FBgn0016694 | Pdp1 | 0 | 2 | upstream | | |
| Dpck | FBgn0003053 | peb-F5-7 | 0 | 1 | upstream | | |
| fbl | FBgn0003053 | peb-F5-7 | 0 | 1 | intron | | Jra (pubmed) |
| Ppcdc | FBgn0002521 | pho | 0 | 1 | upstream | Sp1 | |
| Dpck | FBgn0002521 | pho | 0 | 4 | upstream | Sp1 | |
| CG5828 | FBgn0002521 | pho | 0 | 1 | upstream | Sp1 | |
| fbl | FBgn0002521 | pho | 0 | 1 | intron | Sp1 | |
| ppcs | FBgn0035997 | phol | 0 | 1 | intron | | |
| Ppcdc | FBgn0035997 | phol | 0 | 2 | upstream | | |
| Dpck | FBgn0035997 | phol | 0 | 6 | upstream | | |
| fbl | FBgn0035997 | phol | 0 | 2 | upstream | | |
| ppcs | FBgn0003117 | pnr | 0 | 1 | upstream | tin, tup | NFAT (pubmed) |
| CG5828 | FBgn0003117 | pnr | 0 | 1 | upstream | tup | |
| fbl | FBgn0003117 | pnr | 0 | 3 | intron | | |
| ppcs | FBgn0003118 | pnt | 0 | 2 | upstream | | aop (pubmed) |
| CG5828 | FBgn0003118 | pnt | 0 | 2 | upstream | | aop (pubmed) |
| Dpck | FBgn0003145 | prd | 1 | 0 | upstream | bin, croc | Myc (pubmed) |
| ppcs | FBgn0014018 | Rel | 0 | 2 | intron | br, grh, sqz | |
| ppcs | FBgn0004795 | retn | 0 | 2 | upstream | | |
| ppcs | FBgn0004795 | retn | 0 | 2 | intron | | |
| Ppcdc | FBgn0004795 | retn | 0 | 2 | upstream | | |
| ppcs | FBgn0003254 | rib | 0 | 1 | upstream | ttk, br | |
| ppcs | FBgn0003254 | rib | 0 | 1 | intron | ttk, br | |
| Ppat-Dpck | FBgn0003254 | rib | 0 | 2 | upstream | br | |

| | | | | | | | |
|---|---|---|---|---|---|---|---|
| Dpck | FBgn0003254 | rib | 0 | 2 | upstream | br | |
| CG5828 | FBgn0003254 | rib | 0 | 3 | upstream | br | |
| fbl | FBgn0003254 | rib | 0 | 2 | upstream | ttk, br | |
| fbl | FBgn0003254 | rib | 0 | 1 | intron | ttk, br | |
| ppcs | FBgn0259172 | rn | 0 | 2 | upstream | | |
| Ppat-Dpck | FBgn0259172 | rn | 0 | 6 | upstream | | |
| Dpck | FBgn0259172 | rn | 0 | 2 | upstream | | |
| CG5828 | FBgn0259172 | rn | 0 | 13 | upstream | | |
| ppcs | FBgn0003300 | run | 0 | 1 | upstream | | ttk (pubmed) |
| ppcs | FBgn0013753 | run | 0 | 1 | upstream | run | |
| ppcs | FBgn0003300 | run | 0 | 1 | intron | | ttk (pubmed) |
| ppcs | FBgn0013753 | run | 0 | 1 | intron | run | |
| Ppcdc | FBgn0003300 | run | 0 | 1 | upstream | H | |
| Ppcdc | FBgn0013753 | run | 0 | 1 | upstream | run | |
| Dpck | FBgn0003300 | run | 0 | 1 | upstream | H | |
| Dpck | FBgn0013753 | run | 0 | 1 | upstream | run | |
| fbl | FBgn0003300 | run | 0 | 1 | intron | | ttk (pubmed) |
| fbl | FBgn0013753 | run | 0 | 1 | intron | run | |
| Ppcdc | FBgn0037672 | sage | 0 | 2 | upstream | | |
| Ppat-Dpck | FBgn0000287 | salr-F3-5 | 0 | 2 | upstream | | |
| Dpck | FBgn0000287 | salr-F3-5 | 0 | 2 | upstream | | |
| Ppat-Dpck | FBgn0004170 | sc | 0 | 2 | upstream | da | ac, da, h (pubmed) |
| CG5828 | FBgn0004170 | sc | 0 | 2 | upstream | da | ac, da, tup, h (pubmed) |
| fbl | FBgn0004170 | sc | 0 | 1 | upstream | | ac (pubmed) |
| ppcs | FBgn0002573 | sens | 0 | 1 | upstream | CG5953 | |
| Ppcdc | FBgn0002573 | sens | 0 | 1 | upstream | CG5953 | Dl (pubmed) |
| ppcs | FBgn0051632 | sens2 | 0 | 1 | upstream | | |

| | | | | | | | |
|---|---|---|---|---|---|---|---|
| Ppcdc | FBgn0051632 | sens2 | 0 | 1 | upstream | | |
| ppcs | FBgn0003396 | shn-F1-2 | 0 | 1 | intron | | |
| Ppcdc | FBgn0003396 | shn-F1-2 | 0 | 1 | upstream | | |
| Ppcdc | FBgn0032741 | Side | 0 | 2 | upstream | | |
| ppcs | FBgn0005638 | slbo | 0 | 1 | upstream | | Stat92E (pubmed) |
| Ppat-Dpck | FBgn0005638 | slbo | 0 | 1 | upstream | | |
| fbl | FBgn0005638 | slbo | 0 | 2 | upstream | | |
| ppcs | FBgn0003430 | slp1 | 0 | 2 | upstream | | |
| ppcs | FBgn0003430 | slp1 | 0 | 1 | intron | | |
| Ppcdc | FBgn0003430 | slp1 | 0 | 1 | upstream | | |
| Ppat-Dpck | FBgn0003430 | slp1 | 0 | 1 | upstream | | |
| fbl | FBgn0003430 | slp1 | 0 | 5 | intron | | |
| fbl | FBgn0003430 | slp1 | 0 | 1 | upstream | | |
| ppcs | FBgn0004567 | slp2 | 0 | 2 | upstream | | |
| ppcs | FBgn0004567 | slp2 | 0 | 1 | intron | | |
| Ppcdc | FBgn0004567 | slp2 | 0 | 1 | upstream | | |
| Ppat-Dpck | FBgn0004567 | slp2 | 0 | 1 | upstream | | |
| fbl | FBgn0004567 | slp2 | 0 | 6 | intron | | |
| fbl | FBgn0004567 | slp2 | 0 | 2 | upstream | | |
| ppcs | FBgn0004892 | sob | 0 | 1 | upstream | | |
| ppcs | FBgn0005612 | Sox14 | 0 | 1 | intron | | |
| Ppat-Dpck | FBgn0005612 | Sox14 | 0 | 2 | upstream | | |
| ppcs | FBgn0020378 | Sp1 | 0 | 1 | upstream | | |
| Ppcdc | FBgn0020378 | Sp1 | 0 | 1 | upstream | pho | |
| Dpck | FBgn0020378 | Sp1 | 0 | 6 | upstream | pho | |
| CG5828 | FBgn0020378 | Sp1 | 0 | 3 | upstream | pho | |
| fbl | FBgn0020378 | Sp1 | 0 | 3 | upstream | pho | |

| | | | | | | | |
|---|---|---|---|---|---|---|---|
| ppcs | FBgn0010768 | sqz | 0 | 25 | upstream | Rel | |
| ppcs | FBgn0010768 | sqz | 0 | 20 | intron | Rel | |
| Ppcdc | FBgn0010768 | sqz | 0 | 1 | intron | dimm | |
| Ppat-Dpck | FBgn0010768 | sqz | 0 | 9 | upstream | | |
| Dpck | FBgn0010768 | sqz | 0 | 23 | upstream | dimm | |
| CG5828 | FBgn0010768 | sqz | 0 | 56 | upstream | | |
| fbl | FBgn0010768 | sqz | 0 | 7 | upstream | | |
| fbl | FBgn0010768 | sqz | 0 | 2 | intron | | |
| ppcs | FBgn0003499 | sr | 0 | 1 | intron | | |
| Ppcdc | FBgn0003499 | sr | 0 | 6 | upstream | | |
| ppcs | FBgn0003507 | srp | 0 | 1 | upstream | | |
| Ppcdc | FBgn0003507 | srp | 0 | 2 | upstream | | |
| Ppat-Dpck | FBgn0003507 | srp | 0 | 1 | upstream | | |
| Dpck | FBgn0003507 | srp | 0 | 1 | upstream | | |
| fbl | FBgn0003507 | srp | 0 | 1 | upstream | | |
| fbl | FBgn0003507 | srp | 0 | 1 | intron | | |
| ppcs | FBgn0016917 | STAT92E | 0 | 2 | intron | toy, ttk | slbo (pubmed) |
| ppcs | FBgn0033782 | sug | 0 | 1 | upstream | | |
| Dpck | FBgn0033782 | sug | 0 | 1 | upstream | | |
| CG5828 | FBgn0033782 | sug | 0 | 1 | upstream | | |
| ppcs | FBgn0003651 | svp | 0 | 1 | upstream | | |
| Ppcdc | FBgn0003651 | svp | 0 | 1 | upstream | | |
| CG5828 | FBgn0003651 | svp | 0 | 1 | upstream | | |
| ppcs | FBgn0041092 | tai | 0 | 1 | upstream | usp, ab, EcR | |
| ppcs | FBgn0041092 | tai | 0 | 2 | intron | usp, ab, EcR | |
| Ppcdc | FBgn0023076 | tai | 0 | 1 | upstream | cyc | |
| Ppcdc | FBgn0041092 | tai | 0 | 1 | upstream | usp, ab | |

| | | | | | | | |
|---|---|---|---|---|---|---|---|
| Ppcdc | FBgn0015550 | tap | 0 | 2 | upstream | | |
| ppcs | FBgn0264075 | tgo | 1 | 0 | intron | | |
| ppcs | FBgn0004666 | tgo | 0 | 4 | upstream | tgo | tgo (pubmed) |
| ppcs | FBgn0015014 | tgo | 0 | 10 | upstream | | |
| ppcs | FBgn0015542 | tgo | 0 | 4 | upstream | | |
| ppcs | FBgn0262139 | tgo | 0 | 2 | upstream | tgo | tgo (pubmed) |
| Ppat-Dpck | FBgn0264075 | tgo | 1 | 0 | upstream | | |
| Ppat-Dpck | FBgn0003513 | tgo | 0 | 1 | upstream | tgo | Dll, tgo (pubmed) |
| Ppat-Dpck | FBgn0015014 | tgo | 0 | 3 | upstream | | |
| Dpck | FBgn0004666 | tgo | 0 | 2 | upstream | D, tgo | tgo (pubmed) |
| Dpck | FBgn0015014 | tgo | 0 | 1 | upstream | | |
| fbl | FBgn0015014 | tgo | 0 | 1 | upstream | | |
| ppcs | FBgn0004110 | tin | 0 | 1 | intron | pnr | |
| Ppcdc | FBgn0004110 | tin | 0 | 1 | upstream | | |
| Ppcdc | FBgn0000964 | tj | 0 | 2 | upstream | | |
| CG5828 | FBgn0000964 | tj | 0 | 1 | upstream | | |
| Dpck | FBgn0003720 | tll | 0 | 1 | upstream | | |
| fbl | FBgn0003720 | tll | 0 | 1 | upstream | | |
| ppcs | FBgn0019650 | toy | 0 | 2 | intron | Stat92E | ey (pubmed) |
| Ppcdc | FBgn0019650 | toy | 0 | 2 | upstream | | ey (pubmed) |
| Ppat-Dpck | FBgn0013263 | Trl | 1 | 0 | upstream | | |
| CG5828 | FBgn0013263 | Trl | 1 | 0 | upstream | CG12155, ken | |
| ppcs | FBgn0003870 | ttk | 0 | 1 | intron | gt, Stat92E, rib | run, aop (pubmed) |
| fbl | FBgn0003870 | ttk | 0 | 3 | upstream | gt, D, rib | ac, run (pubmed) |
| ppcs | FBgn0003896 | tup | 0 | 1 | upstream | pnr | |
| Ppcdc | FBgn0003896 | tup | 0 | 2 | upstream | | |
| CG5828 | FBgn0003896 | tup | 0 | 2 | upstream | pnr | sc (pubmed) |

| | | | | | | |
|---|---|---|---|---|---|---|
| ppcs | FBgn0029711 | Usf | 0 | 1 | intron | |
| ppcs | FBgn0003964 | usp | 0 | 1 | upstream | Met, tai, EcR |
| Ppcdc | FBgn0003964 | usp | 0 | 1 | intron | tai |
| Ppat-Dpck | FBgn0003964 | usp | 0 | 1 | upstream | EcR |
| ppcs | FBgn0003986 | Vnd | 0 | 2 | intron | |
| Ppcdc | FBgn0003986 | Vnd | 0 | 1 | intron | |
| fbl | FBgn0003986 | Vnd | 0 | 1 | upstream | |
| ppcs | FBgn0016076 | vri | 0 | 2 | upstream | kay |
| fbl | FBgn0016076 | vri | 0 | 2 | upstream | kay |
| ppcs | FBgn0021872 | Xbp1 | 0 | 1 | upstream | |
| ppcs | FBgn0261113 | Xrp1 | 0 | 1 | upstream | |
| Ppat-Dpck | FBgn0036126 | Xrp1 | 0 | 1 | upstream | Xrp1, crc |
| Ppat-Dpck | FBgn0261113 | Xrp1 | 0 | 1 | upstream | |
| fbl | FBgn0036126 | Xrp1 | 0 | 4 | upstream | Xrp1 |
| fbl | FBgn0261113 | Xrp1 | 0 | 4 | upstream | |
| ppcs | FBgn0004050 | z | 0 | 1 | intron | |
| Ppat-Dpck | FBgn0004050 | z | 1 | 0 | upstream | |
| fbl | FBgn0004050 | z | 0 | 1 | upstream | |
| Ppcdc | FBgn0004053 | zen | 0 | 1 | upstream | |
| Ppat-Dpck | FBgn0004606 | zfh1 | 0 | 1 | upstream | |
| CG5828 | FBgn0004606 | zfh1 | 1 | 0 | upstream | |
| fbl | FBgn0004606 | zfh1 | 0 | 1 | upstream | |
| fbl | FBgn0004606 | zfh1 | 0 | 1 | intron | |
| ppcs | FBgn0259789 | zld | 2 | 1 | intron | |
| ppcs | FBgn0259789 | zld | 1 | 1 | upstream | |
| Ppat-Dpck | FBgn0259789 | zld | 1 | 0 | upstream | |
| Dpck | FBgn0259789 | zld | 1 | 0 | upstream | |

| | | | | | |
|---|---|---|---|---|---|
| CG5828 | FBgn0259789 | zld | 1 | 0 | upstream |
| fbl | FBgn0259789 | zld | 1 | 0 | upstream |

**Supplementary table3. Genotypes used in this study**

| **Figure** | |
|---|---|
| 1, 2c, d, h, S4b | *w1118* |
| 2e, f, g, I, j, k; 3a, c, d; S1e, f, g, j; S2a, b c, d, f, g | *CG31272 (MT)-GAl4, Tub-GAL80TS > +* |
| | *CG31272-GAl4, Tub-GAL80TS > CG5828 (dPANK4)-RNAi* |
| S1b | *esg-GAL4, tub-GAL80TS > +* |
| | *esg-GAL4, tub-GAL80TS > Fbl-RNAi* |
| S1h, i | *CG31272-GAl4, Tub-GAL80TS > +* |
| | *CG31272-GAl4, Tub-GAL80TS > CG5828* |
| 3e | *esg-GAL4, tub-GAL80TS > +* |
| | *esg-GAL4, tub-GAL80TS > Hmgcr-RNAi* |
| | *esg-GAL4, tub-GAL80TS > Qm-RNAi* |
| | *esg-GAL4, tub-GAL80TS > beta GGT-I-RNAi* |
| 3f | *CG31272-GAl4, Tub-GAL80TS > +* |
| | *CG31272-GAl4, Tub-GAL80TS > Drip-RNAi* |
| S2e | *esg-GAL4, tub-GAL80TS > +* |
| | *esg-GAL4, tub-GAL80TS > Mof-RNAi* |
| | *esg-GAL4, tub-GAL80TS > Hat1-RNAi* |
| | *esg-GAL4, tub-GAL80TS > Gcn5-RNAi* |
| S2i | *tsh-GAl4, Tub-GAL80TS > +* |
| | *tsh-GAl4, Tub-GAL80TS > Drip-RNAi* |
| 4a, 5f | *esg-GAL4, tub-GAL80TS > UAS-GFP* |
| | *esg-GAL4, tub-GAL80TS > UAS-GFP, UAS-yki3SA* |
| 4b | *esg- LexA, tub-GAL80TS > +; CG31272>+* |
| | *esg- LexA, tub-GAL80TS > LexAop-yki3SA-GFP 2nd; CG31272>+* |
| | *esg- LexA, tub-GAL80TS > LexAop-yki3SA-GFP 2nd; CG31272>Smvt-RNAi* |
| 4c, d, e, f, g, I, j, k, l, m; S3b, c, d, e, m, l | *esg- LexA, tub-GAL80TS > +; CG31272>+* |
| | *esg- LexA, tub-GAL80TS > LexAop-yki3SA-GFP 2nd; CG31272>+* |
| | *esg- LexA, tub-GAL80TS > LexAop-yki3SA-GFP 2nd; CG31272>Fbl-RNAi* |
| | *esg- LexA, tub-GAL80TS > LexAop-yki3SA-GFP 2nd; CG31272>CG5828* |
| S3f, g, h, I, j, k | *esg- LexA, tub-GAL80TS > +; CG31272>+* |
| | *esg- LexA, tub-GAL80TS > LexAop-yki3SA-GFP 3rd; CG31272>+* |

| | |
|---|---|
| | *esg- LexA, tub-GAL80TS > LexAop-yki3SA-GFP 3rd; CG31272>Fbl-RNAi* |
| | *esg- LexA, tub-GAL80TS > LexAop-yki3SA-GFP 3rd; CG31272>CG5828* |
| | *CG31272-GAl4, Tub-GAL80TS > +* |
| 5a | *CG31272-GAl4, Tub-GAL80TS > Myc* |
| 5c | *CG31272-GAl4, Tub-GAL80TS > Myc-HA* |
| | *CG31272-GAl4, Tub-GAL80TS > +* |
| S4c, d | *CG31272-GAl4, Tub-GAL80TS > Myc-RNAi* |
| | *esg- LexA, tub-GAL80TS > +; CG31272>+* |
| | *esg- LexA, tub-GAL80TS > LexAop-yki3SA-GFP 2nd; CG31272>+* |
| 5g, h, I, j, k, l, m n , S4e, f, g, h | *esg- LexA, tub-GAL80TS > LexAop-yki3SA-GFP 2nd; CG31272>Myc-RNAi* |
| | *esg- LexA, tub-GAL80TS > +; CG31272>+* |
| | *esg- LexA, tub-GAL80TS > LexAop-yki3SA-GFP 3rd; CG31272>+* |
| S4i, j, k, l, m, n | *esg- LexA, tub-GAL80TS > LexAop-yki3SA-GFP 3rd; CG31272>Myc-RNAi* |
| | *CG31272-GAl4, Tub-GAL80TS > +* |
| 5o | *CG31272-GAl4, Tub-GAL80TS > PvR[Act]* |
| | *esg- LexA, tub-GAL80TS > +; CG31272>+* |
| | *esg- LexA, tub-GAL80TS > LexAop-yki3SA-GFP 2nd; CG31272>+* |
| 5p | *esg- LexA, tub-GAL80TS > LexAop-yki3SA-GFP 2nd; CG31272>Pvr-RNAi* |

| Supplementary table4. Primers used in this study | | |
|---|---|---|
| | | **RT-qPCR primers** |
| dPANK4 (CG5828) | Forward | TTACAGATCCCTGGCTGAGAC |
| | Reverse | CCACCTTGTGTCCTCATCCG |
| Fbl | Forward | TTCTCTTCGCCGATCTGCATA |
| | Reverse | GAACTGCTGCTTTTCGCTTTTTA |
| FASN1 | Forward | GACATGGTCAACGATGATCCC |
| | Reverse | ACCGAAGAACTGTTGGTCAAAG |
| ACC | Forward | ACAAGATGAAGAACCATGCCAT |
| | Reverse | TTCGCGGGACTTCTGTTGC |
| AcCoAS | Forward | CCATGATTCTGGAGCTGCCTA |
| | Reverse | GCCTTCAGGTACAGGGGTTTC |
| Atpcl | Forward | TTTCCACAGTAAATTCCACGACA |
| | Reverse | GGCGCTTGATAAGTTGATCGG |
| Gcn5 | Forward | GGTGGAAACAAGAGGACCAGTG |
| | Reverse | CCAAATTCTCACTGCTTGGA |
| Elp3 | Forward | AATTCTGCTTCCAAAGCTGAGG |
| | Reverse | GCCGGGACAATAGACGCATA |
| Hat1 | Forward | TGGTAGACTTTAAGCTGATCCGT |
| | Reverse | CTCCCCGAAAATCTGGTGGG |
| Mof | Forward | GAGCCAACCGATGCGTACA |
| | Reverse | TCCTCCGAAATGGGACTGATG |
| Nej | Forward | ATGATGGCCGATCACTTAGACG |
| | Reverse | GATTTGTGGTTACACCGGAGG |
| Fpps | Forward | GCAACGCCTGATCTCTACCAG |
| | Reverse | TTGGAGCGTCGATAAGGTTCT |
| Hmgcr | Forward | GCTGCACTGCCGTACTGTA |
| | Reverse | AATGCCCAGCACATATTTGGA |
| Qm | Forward | TAAATGCGGCCAACTATGCAC |
| | Reverse | CATCAGCTTGTAATCCGACTCG |
| beta GGT-I | Forward | ATGGCCTCGCACGATAACAC |
| | Reverse | GCAATGAGTTTAGCACATCCAGG |
| CG13200 | Forward | GCATATGCGACAAAGTGGGCC |
| | Reverse | AACATTCACCGCAAGGGCTCC |
| RP49 | Forward | AAGAAGCGCACCAAGCACTTCATC |
| | Reverse | TCTGTTGTCGATACCCTTGGGCTT |
| | | **ChIP-qPCR primers** |
| dPANK4 (CG5828) | Forward | CCAGCTGAGGTGTGCTGG |
| | Reverse | TTGAACAGTCTTATTGCAACTATCG |

| | | |
|---|---|---|
| Fbl | Forward | GGTGCACATAAAATGTGTGGGA |
| | Reverse | ATCGAAAAGCGCAGTGTTGG |
| Tll (Neg) | Forward | CCTTCTTGAATTTCCAGGTCGC |
| | Reverse | CGTCTTGTCCACCACACAGA |
| | | **CG5828 into TOPO clone** |
| dPANK4 (CG5828) | Forward | caccATGTACAGTAGCAGCTTGCTGCCG |
| | Reverse | CTAGCTGGGCGCAGCCGGTTCGAA |
| | | **CG5828 into pGEX-4T2 InFusion** |
| dPANK4 (CG5828) | Forward | TCCCCAGGAATTCCCATGTACAGTAGCAGCTTGCTGCC |
| | Reverse | CGCTCGAGTCGACCCCTAGCTGGGCGCAGCCGG |
| dPANK4 (CG5828)-D209A | Forward | GGTCTTTGTGGCGAACAGCGGCG |
| | Reverse | ACGGCGCACTTGTGT |
| dPANK4 (CG5828)-D245A | Forward | TGCTCTAAATGCGGTGACCAGCG |
| | Reverse | GGTTCACTGTTGGCG |
